# Working memory operations emerge from dynamic changes in neural subspace geometry

**DOI:** 10.64898/2026.08.31.748237

**Authors:** Aniol Santo-Angles, M Gyurkovics, K Jaworska, JM Palva, G Thut, S Palva

**Affiliations:** Neuroscience Center, Helsinki Institute of Life Science HiLIFE, University of Helsinki, Finland; Centre for Cognitive Neuroimaging, School of Psychology and Neuroscience, University of Glasgow, Glasgow, United Kingdom; School of Psychology, University of East Anglia, Norwich, United Kingdom; Centre de Recherche Cerveau et Cognition (CerCo), CNRS UMR5549 and Université de Toulouse, Toulouse, France; Department of Neuroscience and Biomedical Engineering, Aalto University, Espoo, Finland

## Abstract

Working memory representations of multiple items are maintained within quasi-orthogonal neural subspaces, a mechanism thought to reduce interference between memory contents. Previous studies in both non-human and human primates have demonstrated the existence of such subspaces, but it remains unclear how cognitive operations are performed on the information they encode. One hypothesis proposes that these operations are implemented through dynamic changes in subspace geometry, yet direct empirical evidence for this idea in the context of working memory is lacking. Furthermore, the neural mechanisms underlying such geometric reconfigurations remain unknown, particularly the role of oscillatory and aperiodic neural activity. Here, we addressed these questions using simultaneous magneto– and electroencephalography (M/EEG) recordings from healthy human participants performing a multi-item, multi-feature delayed match-to-sample visual WM task. Neural subspaces were estimated using dimensionality reduction techniques, and their geometry was characterized by quantifying the orthogonality between subspaces, distances between memory representations and subspace shapes. Participants maintained orientation and shape information simultaneously while task demands required either deprioritizing one feature or updating the representation of the prioritized feature. We found that subspace geometry was flexibly reshaped according to task demands: prioritization expanded representational subspaces, whereas de-prioritization shrunk them. In addition, subspaces representing different feature domains were generally oblique and became more orthogonal only under high cognitive demands. Finally, these task-dependent changes in subspace geometry were largely driven by interactions between aperiodic neural activity and slow oscillations in the delta and theta frequency bands.

## INTRODUCTION

Working memory (WM), the ability to hold and manipulate information over short periods ^1,2^, is supported by neural activity that maintains memory representations in the absence of external stimuli, distributed across large-scale brain networks ^3,4^. There is ongoing debate as to whether this neural activity is sustained throughout the retention period ^5–7^, is organized into transient oscillatory events ^8–10^, or emerges from an interplay between activity-dependent and activity-silent mechanisms depending on task demands ^11–14^. An emerging perspective based on population-level analyses of multi-neuron recordings reframes WM as an emergent property of dynamic neural population activity, in which transient, heterogeneous single-neuron responses are organized into stable, sustained, low-dimensional population geometries that preserve memory content ^14–17^. A growing body of work has focused on characterizing WM-related low-dimensional subspaces, also referred to as neural manifolds, that maintain representations of multiple stimuli of the same feature dimension, such as color or spatial location. In multi-item WM, individual items are maintained in distinct low-dimensional subspaces that reduce interference and support behaviorally relevant coding ^18,19^. When multiple items are held simultaneously, their representations tend to occupy quasi-orthogonal subspaces, which can reorganize when attentional priority shifts, becoming more aligned as previously relevant items lose priority ^18^. In sequential tasks, items are encoded in distinct subspaces according to ordinal position ^19^, and these representations can dynamically remap—for example, switching subspaces when the recall order is reversed ^20^. A recent MEG study in humans showed that memory contents are maintained in quasi-orthogonal neural subspaces organized by ordinal position in sequential WM tasks, and that the geometry of these subspaces is behaviorally relevant ^21^, extending earlier findings in non-human primates ^19^. These studies provide a mechanistic account of how the brain can simultaneously maintain multiple items of the same type while limiting mutual interference. Classical attractor-based WM models explain the maintenance of a single item through persistent activity sustained by recurrent excitatory connectivity within neural populations, where stable attractor states preserve information over time despite ongoing neural noise ^7,22,23^. However, these models often exhibit catastrophic interference when multiple items must be stored simultaneously, as overlapping population representations compete with one another, degrading the fidelity of individual memories and leading to rapid loss of stored information ^24^. Importantly, it remains unclear whether WM-related subspaces generalize to situations in which multiple items from different stimulus domains are maintained simultaneously. Are memory representations organized into subspaces only when they are maintained within the same neuronal populations (e.g., multiple stimuli of the same feature dimension), as a mechanism to reduce mutual interference? If so, subspaces encoding different stimulus features (e.g., orientation and shape) should not exhibit systematic orthogonalization.

Alternatively, WM representations may be organized into subspaces regardless of content type, reflecting a more general computational principle. Previous studies showing that memory representations can be dissociated from sensory and motor representations through subspace structure ^25,26^ are consistent with this latter possibility. In the present study, we address this issue directly by characterizing the geometry of neural subspaces encoding memory representations of orientations and polygonal shapes.

Another open question concerns the relationship between subspace dynamics, task demands, and behavioral performance. Prior work has shown that deprioritizing a memory item by shifting attention away from it during the retention period reduces the orthogonality between neural subspaces, increasing their overlap ^18^. However, this effect may arise from a transition in representational structure from a two-item to a one-item state, reflecting a release or pruning of representations rather than an active reconfiguration of subspace geometry. In parallel, several studies have linked subspace geometry to behavior, showing that error trials are associated with greater overlap between subspaces ^19,21^ and reduced separation between memory representations within subspaces ^18,21^. Despite these findings, the relationship between cognitive operations and subspace dynamics remains unclear. It has been proposed that neural computations are implemented through changes in trajectories within low-dimensional neural manifolds ^27,28^. However, whether cognitive operations such as prioritization or substitution of memory items could originate from dynamical changes in subspace geometry is unknown. In the present study, we test whether cognitive operations actively reshape subspace geometry and whether such changes are behaviorally relevant.

A central unresolved question is how the geometry of WM-related neural subspaces relates to the different components of neural activity, and in particular whether these representational structures are primarily shaped by aperiodic/broadband activity or by narrow-band oscillatory dynamics. While there is now substantial evidence that WM can be described in terms of low-dimensional subspaces ^18–21^, the nature of the neural signals that instantiate these geometries is still largely unresolved. Most studies on the neural underpinnings of WM-related subspace geometry have relied on spike trains in non-human primates ^18–20^ or broadband source modelled MEG data in humans ^21^. In a recent intracranial recording study in the macaque prefrontal cortex during sequential WM, ^29^ found that transient increases in theta-band power carried information about stimulus identity, while theta– spike coupling selectively tracked WM subspaces that encoded the ordinal position of items in the sequence. This finding might indicate that distinct components of WM geometry may be supported by separable oscillatory mechanisms. More broadly, this view aligns with a large body of work implicating neural oscillations in WM, where different bands support distinct aspects of memory processing ^30–32^. Theta oscillations, particularly in frontomedial regions, have been implicated in higher-order cognitive control and coordination of distributed WM processes ^33,34^, as well as in functions such as cyclic sampling of visual space in the frontal eye fields ^35^ and the encoding and maintenance of sequential information in the hippocampus ^36–40^. In contrast, alpha oscillation amplitude has traditionally been interpreted as a mechanism of functional inhibition, with increased alpha amplitude suppressing the processing of task-irrelevant information and thereby protecting relevant sensory and mnemonic representations ^41,42^. More recent evidence, however, challenges the notion that alpha simply indexes neural suppression. Alpha amplitude and long-range alpha synchronization has also been linked to the active selection and prioritization of relevant information and to the modulation of representations held in memory ^43–48^. Beta and gamma amplitudes have been linked to the active maintenance of memory contents content in MEG ^31,49,50^, intracranial EEG ^36^ and non-human primate LFP ^8,9,51^, as well as WM operations including prioritization, read-out, and post-response updating, often expressed through transient beta bursts ^8,9,51,52^. Those fast oscillations often interact with slower rhythms that help organize and integrate information ^9,37,53–55^. However, how these multiscale oscillatory dynamics are linked to the formation and control of WM-related subspaces remains an open question. At the same time, growing evidence has highlighted the importance of aperiodic neural activity and its potential to confound or reshape interpretations of narrowband oscillatory dynamics ^56–61^. Aperiodic components reflect the scale-free, 1/f structure of neural activity and are thought to index global neural excitability as well as the balance between excitation and inhibition in cortical circuits ^62,63^. In the context of WM, aperiodic activity has been increasingly linked to behaviorally relevant variability. Early evidence showed that aperiodic spectral components vary systematically with age-related differences in WM performance ^64^. During WM delay periods, posterior aperiodic activity contralateral to attended items exhibits reliable reductions in broadband power relative to pre-stimulus baseline, and these changes track behavioral accuracy ^56^. Extending this work, ^65^ demonstrated that increases in 1/f slope steepness over fronto-central electrodes during the delay period are associated not only with WM capacity, but also with performance across a broader set of complex span tasks. In summary, these findings raise the possibility that some of the neural dynamics attributed to oscillatory mechanisms in WM, especially to frontal theta activity ^66,67^, may partially reflect, or be modulated by, underlying aperiodic changes in cortical activity, underscoring the need to disentangle these components when relating neural population dynamics to WM-related subspaces.

In the present study, we addressed whether the geometry of low-dimensional neural subspaces organizing memory contents is dynamically reshaped by changing task demands, and whether subspace organization and dynamics are related to ongoing neural oscillations or aperiodic activity. Participants initially encoded gratings (orientation) and polygons (shape) and maintained them across a delay period (see Figure 1A, upper panels). A subsequent cue instructed participants either to maintain all stored information (control condition), deprioritize one feature while prioritizing the other (inhibition condition), or update one memory item by replacing it with a newly presented stimulus while also deprioritizing the non-updated feature (updating condition) (Figure 1A, lower panels). By keeping the sensory input identical across conditions while varying the required cognitive operations, we isolated how attentional prioritization and updating reshape the geometry of neural representations in working memory space. Concurrent magneto– and electroencephalography (M/EEG) were recorded while participants performed the task. We hypothesized that (1) memory representations of shape and orientation would be organized within low-dimensional neural subspaces; (2) subspaces encoding different stimulus features would exhibit an orthogonal geometry; (3) subspace geometry would be dynamically modulated by task demands, such that (3.1) orthogonality between subspaces following the cue would increase in the control condition but decrease in the inhibition and update conditions, when discrimination between features is no longer required, and (3.2) the separability of memory representations within prioritized subspaces would increase (subspace expansion), whereas the separability of representations within deprioritized subspaces would decrease (subspace shrinkage); (4) these task-dependent modulations would be reduced or absent during incorrect trials; and (5) subspaces geometry would be shaped by theta-band oscillatory activity.

**Figure 1.**
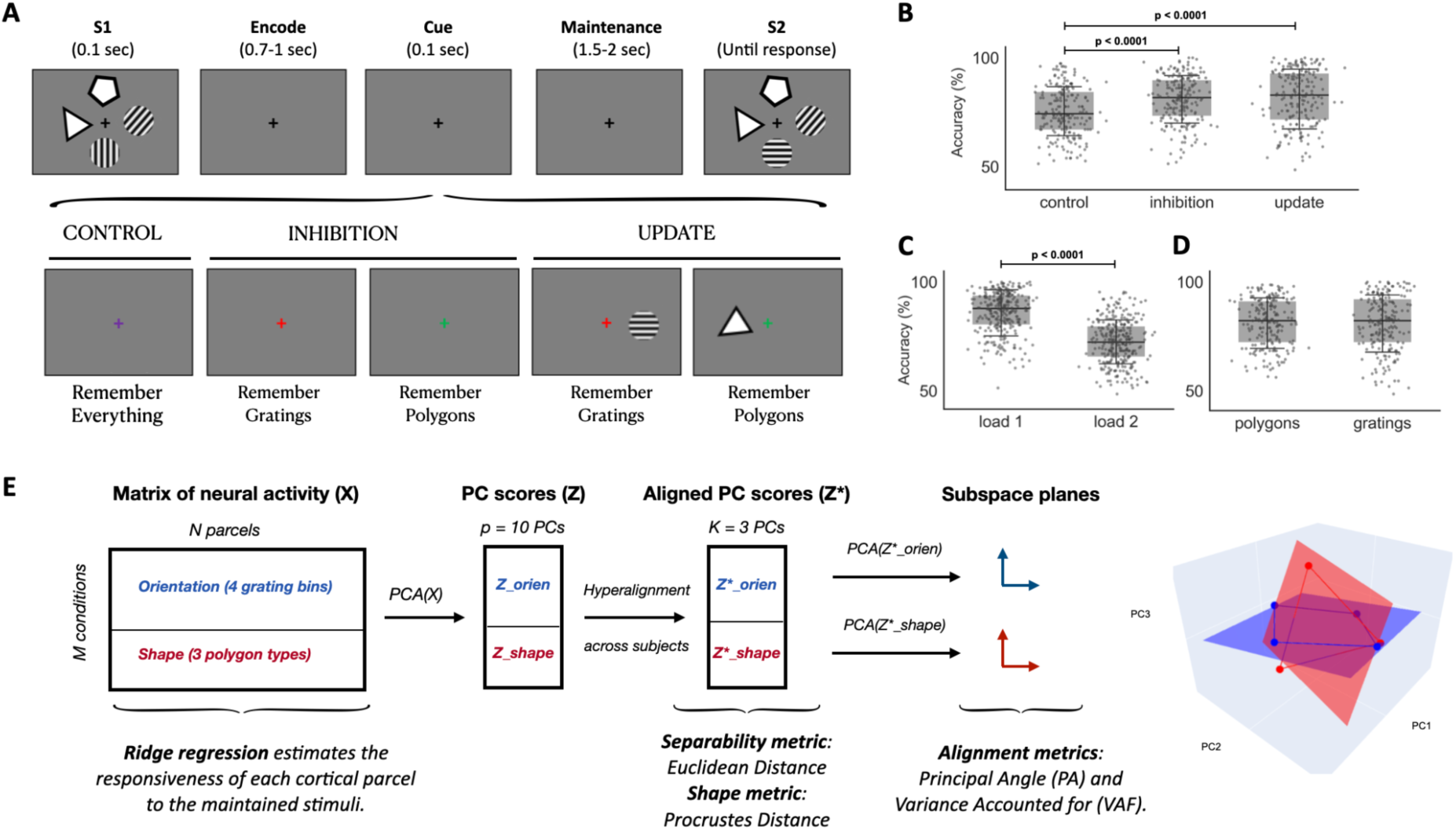
Task design, behavioral performance, and geometric analysis of neural subspaces. (A) Task design. (B–D) Behavioral performance. Boxplots show accuracy across (B) experimental conditions (control, inhibition, update), (C) load levels (1 vs. 2), and (D) relevant stimulus features (polygons vs. gratings) in the inhibition and update conditions. Individual subject data are overlaid as points. P-values were obtained from linear mixed-effects models. (E) Geometric analysis of neural subspaces. Schematic of the analysis pipeline. The neural activity matrix X was organized by stimulus conditions, with M = 7 conditions as rows and N = 400 brain parcels (channels) as columns. To estimate low-dimensional neural subspaces, we applied PCA along the channel dimension and projected X onto the top p = 10 principal components (PCs), yielding the PC-score matrix Z (M = 7 conditions × p = 10 PCs). Z was then hyperaligned across subjects to obtain a common representational space (see below). From the aligned matrix Z* (M = 7 × p = 10), we retained the top k = 3 PCs to define the neural subspaces, yielding a 7 × 3 matrix. This matrix was subsequently partitioned by feature into orientation (M = 4 conditions × k = 3 PCs) and shape (M = 3 conditions × k = 3 PCs) subspaces. Finally, we applied PCA separately to each feature-specific subspace and defined its geometric plane using the two leading PCs. See Methods for details.

## RESULTS

All participants performed the task above chance. Binomial tests confirmed that accuracy exceeded chance level (50%) when pooling trials across all conditions and loads (83 ± 8% [63, 95]; all p < 0.05). Considering loads separately, performance was above chance for load 1 (89 ± 8% [66, 98]; all p < 0.05) and load 2 (78 ± 8% [60, 92]; all p < 0.05). Across conditions, all participants performed above chance in control (86 ± 7% [66, 96]) and inhibition (81 ± 8% [59, 95]), while one participant did not exceed chance in update (81 ± 10% [39, 96]; 49 participants p < 0.05). Considering the combination of condition and load, one participant performed at chance in update load 1 (89 ± 12% [26, 99]; 49 participants p < 0.05) and update load 2 (73 ± 10% [52, 92]; 49 participants p < 0.05), and three participants performed at chance in control load 2 (69 ± 8% [52, 86]; 47 participants p < 0.05). All subjects were included in the main analyses.

Behavioral accuracy differed across conditions (Figure 1B). A linear mixed-effects model revealed a significant main effect of condition (inhibition > control: t = 8.973, p < 0.001; update > control: t = 9.264, p < 0.001). Post-hoc Tukey comparisons confirmed that accuracy was lower in the control condition compared with both inhibition (mean difference = 5.42, p < 0.001) and update (mean difference = 5.60, p < 0.001), whereas accuracy did not differ between the inhibition and update conditions (mean difference = 0.18, p = 0.988). There was also a significant main effect of load, with accuracy being lower in load 2 than load 1 (t = –27.047, p < 0.001) (Figure 1C). Finally, in inhibition and update trials (excluding control trials), the effect of which stimulus feature became relevant after the cue (polygons vs. gratings) was not significant (t = 0.311, p = 0.756), indicating that behavioral accuracy did not differ depending on which feature remained relevant (Figure 1D).

### Geometric analysis of neural subspaces

To test hypotheses 1-4 positing that the contents of WM for multiple stimulus features are maintained simultaneously in low-dimensional neural subspaces specific to each feature and that this is differentially expressed as a function of task demands and behavioural performance, we applied a geometric analysis of neural subspaces (Figure 1E). This approach has previously been used to characterize subspaces encoding multiple stimuli of the same feature in intracranial recordings from non-human primates ^18,19^ and MEG in humans ^21^. Here, we used this framework to provide a novel characterization of the neural subspaces encoding two distinct stimulus features: orientation and shape. We analyzed source-reconstructed MEG data from a sample of 50 healthy participants, parcellated into 400 cortical parcels ^68^. We constructed time-resolved, subject-specific stimulus-by-channel matrices, with rows representing distinct stimulus identities of each feature (specific orientations or shapes) and columns representing cortical parcels. Low-dimensional subspaces were estimated by projecting these matrices onto the leading three Principal Components (PC), which together explained 87 ± 5% of the variance (range: 70–99%; PC1: 54 ± 13%, PC2: 21 ± 7%, PC3: 12 ± 5%). These results support the low dimensionality of WM-related subspaces (hypothesis 1).

We next examined the spatial topography by mapping the PCs onto the cortical surface, summarizing the eigenvalue-weighted squared eigenvectors across participants, time windows, and conditions using a hierarchical statistical model (see Methods). We squared the eigenvector loadings to quantify their amplitude, rather than their direction, because the sign of PCA eigenvectors is arbitrary and can be reversed without changing the underlying subspace. We first assessed whether contributions to the subspaces were restricted to a subset of parcels or distributed across the cortex. Contributions were significantly greater than zero in all 400 parcels for each of the three leading PCs. Specifically, t-statistics were positive across all parcels for PC1 (mean = 16, SD = 2, range = 11–22), PC2 (mean = 20, SD = 3, range = 11–29), and PC3 (mean = 18, SD = 2, range = 10–26). This indicates that the neural subspaces were broadly distributed across the cortex rather than being driven by a restricted set of cortical parcels. We next characterized relative parcel contributions by z-scoring component contributions across parcels within each participant. Positive values therefore indicated above-average contributions, whereas negative values indicated below-average contributions. The resulting parcel-wise t-statistics revealed distinct spatial topographies across the three PCs (see Methods). PC1 showed relatively greater contributions in posterior medial regions, particularly the posterior cingulate/precuneus, as well as somatomotor regions, whereas relatively lower contributions were concentrated in visual and prefrontal regions (Figure 2G). PC2 showed relatively greater contributions in frontal and anterior cingulate regions, whereas relatively lower contributions were concentrated in visual and parietal regions (Figure 2H). PC3 showed relatively greater contributions in visual and prefrontal regions, particularly medial prefrontal areas, whereas relatively lower contributions were observed in somatomotor and salience/ventral attention regions, including lateral fronto-insular/opercular areas, as well as posterior medial regions encompassing the precuneus and posterior cingulate cortex (Figure 2I).

**Figure 2.**
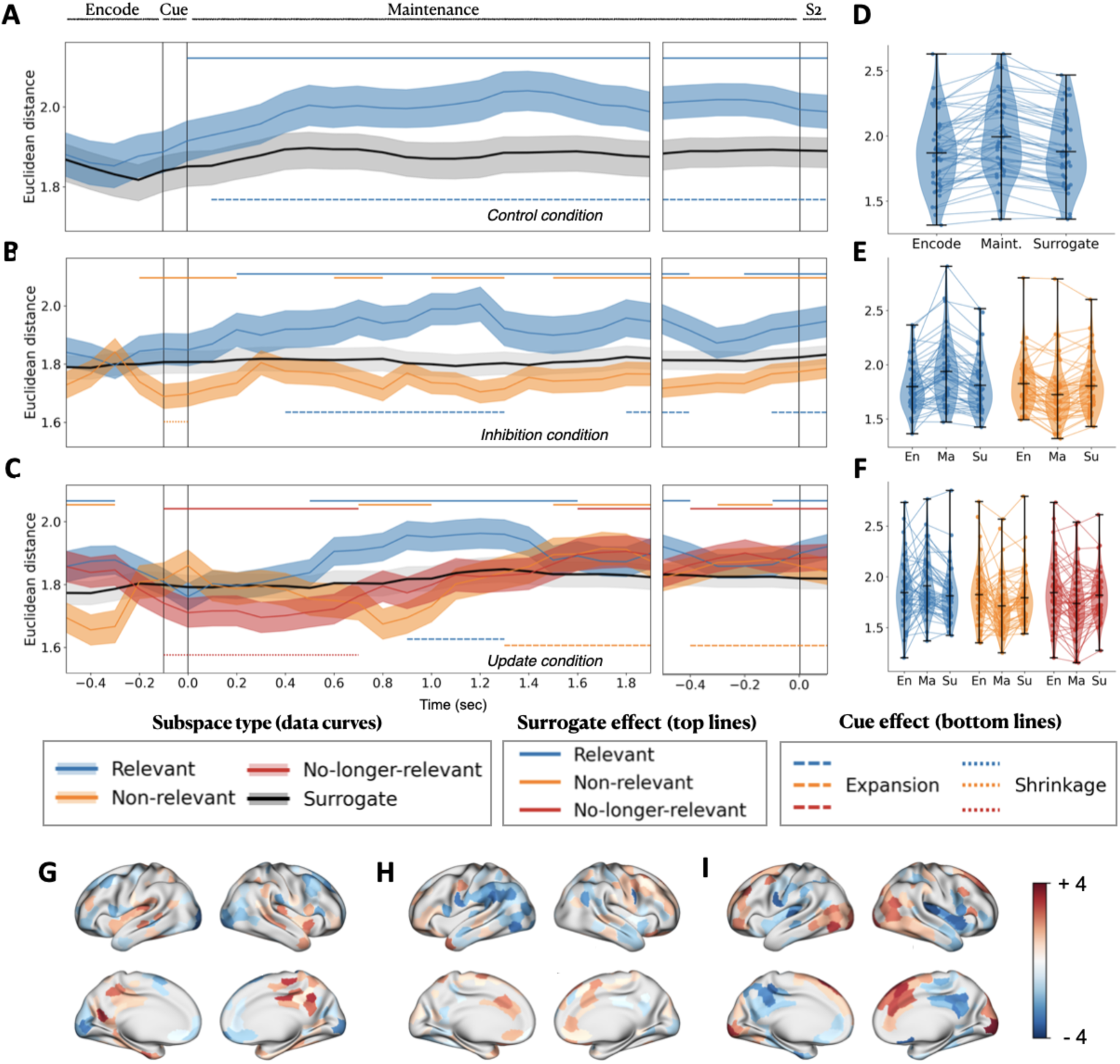
Separability of memory representations within feature-specific neural subspaces, quantified as the Euclidean distance between rows in the Z* matrices of aligned PC scores (y-axis), combining stimulus loads 1 and 2. A–C) Time-resolved separability for A) control, B) inhibition, and C) update conditions. Separate panels reflect different jitter-corrected maintenance duration (1.5-2 sec). Solid vertical lines mark cue, maintenance and S2 onsets (left to right). The x-axis shows time in milliseconds, representing the onset of 0.4 sec time windows in 0.1 sec steps (e.g., x = 0 ms corresponds to the interval [0, 0.4] sec). Colored solid lines represent mean Euclidean distances across subjects, with blue denoting relevant features, orange non-relevant features, and red no-longer-relevant features. Black solid lines represent separability estimated from surrogate data, averaged across 1,000 surrogate iterations. For visualization purposes, only the maintenance period is shown; the full sequence of trial events is provided in Figure S2. Only the surrogate trace for the relevant-feature subspace is shown here; surrogate traces for the other feature-specific subspaces are provided in Figures S3 and S4. Shaded regions indicate the standard error of the mean (SEM) across subjects. Horizontal bars at the top indicate time points at which separability significantly differed from surrogate data, whereas bars at the bottom indicate time points showing a significant cue effect (dependent-samples, two-tailed t-test; FDR-corrected, p<0.05). D–F) Violin plots of Euclidean distance for D) control, E) inhibition, and F) update conditions, averaged across time windows exhibiting a significant effect relative to surrogate data. Distances are shown separately for the encoding period (En), maintenance period (Ma), and surrogate data during the maintenance period (Su). Individual dots represent individual subjects, and lines connect observations from the same subject. G–I) Cortical maps showing the spatial topography of the three leading PCs (PC1–PC3), based on eigenvalue-weighted squared eigenvector contributions during the maintenance period. Maps show parcel-wise t-statistics summarizing relative component contributions across subjects, time windows, and conditions. For visualization, maps were thresholded to display the 50% of cortical parcels with the largest absolute t-statistics, while preserving the sign of the t-statistic; positive values indicate above-average relative contributions and negative values indicate below-average relative contributions. These t-statistics are uncorrected and are shown for descriptive visualization only, and the threshold should not be interpreted as a statistical significance threshold. G) PC1, H) PC2, and I) PC3. En, encoding delay period; Ma, maintenance delay period; Su, surrogate data.

Having established the dimensionality and spatial organization of the feature-specific neural subspaces, we next characterized their geometry. To enable comparisons of subspaces across participants despite individual differences in their neural representational spaces, subspaces were hyperaligned across participants to define a common representational space ^69,70^. Hyperalignment was performed on the PC-score matrix Z (M = 7 conditions × p = 10 PCs). Following hyperalignment, we retained the leading *k* = 3 PCs for downstream analyses; we refer to these as the subspaces (see Methods). Varying *k* and *p* around these values produced similar results, indicating that the main findings are robust to the specific choice of parameters (see Supplementary Material). We then quantified the geometry of subspaces by a) the alignment between subspaces, quantified by the principal angle (PA) and the variance accounted for (VAF), which capture the degree of orthogonality versus parallelism between subspaces (hypothesis 2 and 3.1); b) the separability of memory representations within each subspace, measured as the Euclidean distance between stimulus-specific representations (hypothesis 3.2) ^21^; and c) shape metrics describing the global configuration of the representations within each subspace (hypothesis 3.2) ^69^ (see Figure 1E for an overview of the geometric pipeline). To assess how subspace geometry was modulated by task demands (hypothesis 3), including changes in subspace orthogonality (hypothesis 3.1) and representational separability (hypothesis 3.2), we distinguished two complementary effects. The cue effect captured changes in geometric variables induced by the cue, quantified as the difference between the post-cue delay period (maintenance) and the average pre-cue period (encode). The surrogate effect assessed whether geometric variables during the pre-cue (encode) and post-cue (maintenance) delay periods differed from chance levels estimated using surrogate data (1,000 iterations). When both effects were significant, we reported the overall change in geometry; when only one effect was significant, we specified the effect driving the observed change.

#### Alignment

Subspaces were generally oblique (semi-orthogonal), although with substantial inter-subject variability (Figure S1 in Supplementary Material). Principal angles (PA), averaged over time within each participant and then across participants, were 64° ± 10 (range: 34–84) in the control condition, 60° ± 10 (30–80) in the inhibition condition, and 63° ± 10 (31–80) in the update condition. VAF analyses led to the same qualitative conclusion, with higher VAF values indicating greater alignment (i.e., lower orthogonality) between subspaces (Control: 0.62 ± 0.12 [0.31 0.89]; Inhibition: 0.66 ± 0.12 [0.34 0.96]; Update: 0.64 ± 0.11 [0.36 0.95]). Although these descriptive values indicate that orientation and shape were represented in semi-orthogonal subspaces (hypothesis 2), comparison with surrogate data showed that subspace orthogonality did not exceed chance levels before the cue (Figure S1). Following the cue, subspaces became significantly more orthogonal than expected by chance (surrogate effect) only in the control condition when participants maintained two stimuli per feature (load 2; Figure S1B,E), supporting Hypothesis 3.1. This effect was not observed when participants maintained one stimulus per feature (load 1; Figure S1A,C). Moreover, this increase in orthogonality was absent in incorrect trials, supporting hypothesis 4 (Figure S1C,F). No robust cue– or surrogate-related effects were observed in the inhibition or update conditions, aside from brief isolated significant time points that likely reflected random fluctuations (Figure S1). Together, these findings indicate that subspace orthogonality is dynamically modulated by task demands and is behaviorally relevant, but emerging only under the highest WM demands, when all four memory items had to be maintained following the cue.

#### Separability

We investigated the effect of cue-driven prioritization on the geometry of memory representations by measuring the Euclidean distance between memory representations within each subspace, which served as a measure of representational separability. To assess how separability was modulated by task demands (hypothesis 3.2), we used the cue and surrogate effects described above. We interpreted the resulting increase or decrease in separability as subspace expansion or shrinkage, respectively.

In the control condition, where both stimulus features remain relevant throughout the trial, subspaces expanded after the cue (Figure 2A,D). This pattern holded when analyzing loads 1 and 2 separately (Figure S5C–D). Considering features individually, gratings subspace expanded after the cue, whereas polygons remained above chance level throughout the delay and did not show a cue-related increase (Figure S5A–B).

In the inhibition condition (Figure 2B,E), relevant subspace expanded after the cue, while non-relevant subspace shrunk after the cue, mainly driven by the surrogate effect. When analyzed separately by feature, the expansion of the relevant subspace and shrinkage on non-relevant subspace was observed for both polygons and gratings (Figure S6A–B). Considering loads individually, these effects remained significant for load 2 (Figure S6D). For load 1, however, we observed the opposite pattern, expansion of non-relevant and shrinkage of relevant subspace (Figure S6C). The effect observed at load 1 was not attributable to any single stimulus feature; it was evident when polygons and gratings were analyzed independently (Figure S7).

In the update condition (Figure 2C,E), relevant subspaces expanded later in the maintenance period, rather than immediately after the cue as observed in other conditions. Notably, in this condition, the items represented in the relevant subspaces were updated during cue presentation. Non-relevant subspaces briefly shrank (surrogate effect) at the same time that relevant subspaces expanded, then expanded again toward the end of the maintenance period, near S2. The no-longer-relevant subspace, only present in the update condition, encodes the feature that remains relevant after the cue, but contains the original stimuli presented during S1 rather than the updated stimuli presented alongside the cue. This subspace showed significant shrinkage immediately after the cue, followed by a surrogate-driven expansion near S2. These patterns were consistent across stimulus features and loads (Figure S8). The observed patterns of subspace expansion and shrinkage were consistent across individual subjects within each experimental condition (Figure 2E–F).

We next assessed whether these effects of separability were behaviorally relevant (hypothesis 4) by examining the same analyses on incorrect trials. To minimize confounds arising from the smaller number of incorrect trials, we restricted this analysis to the inhibition and update conditions at load 2 (Table 1). To ensure that representations from correct and incorrect trials were expressed in a common representational space, we projected the incorrect-trial data onto the PC basis derived from the correct-trial data and applied the hyperalignment transformation matrices estimated from correct trials (see Methods). This procedure ensured that both correct and incorrect representations were embedded in the same representational space, allowing any performance-related differences in geometric variables to be attributed to representational geometry rather than misalignment between representational spaces. The overall expansion of task-relevant subspaces and shrinkage of non-relevant (and no-longer-relevant) subspaces observed in the inhibition and update conditions was absent in incorrect trials, supporting the behavioral relevance of subspace modulations (Figure 3).

**Figure 3.**
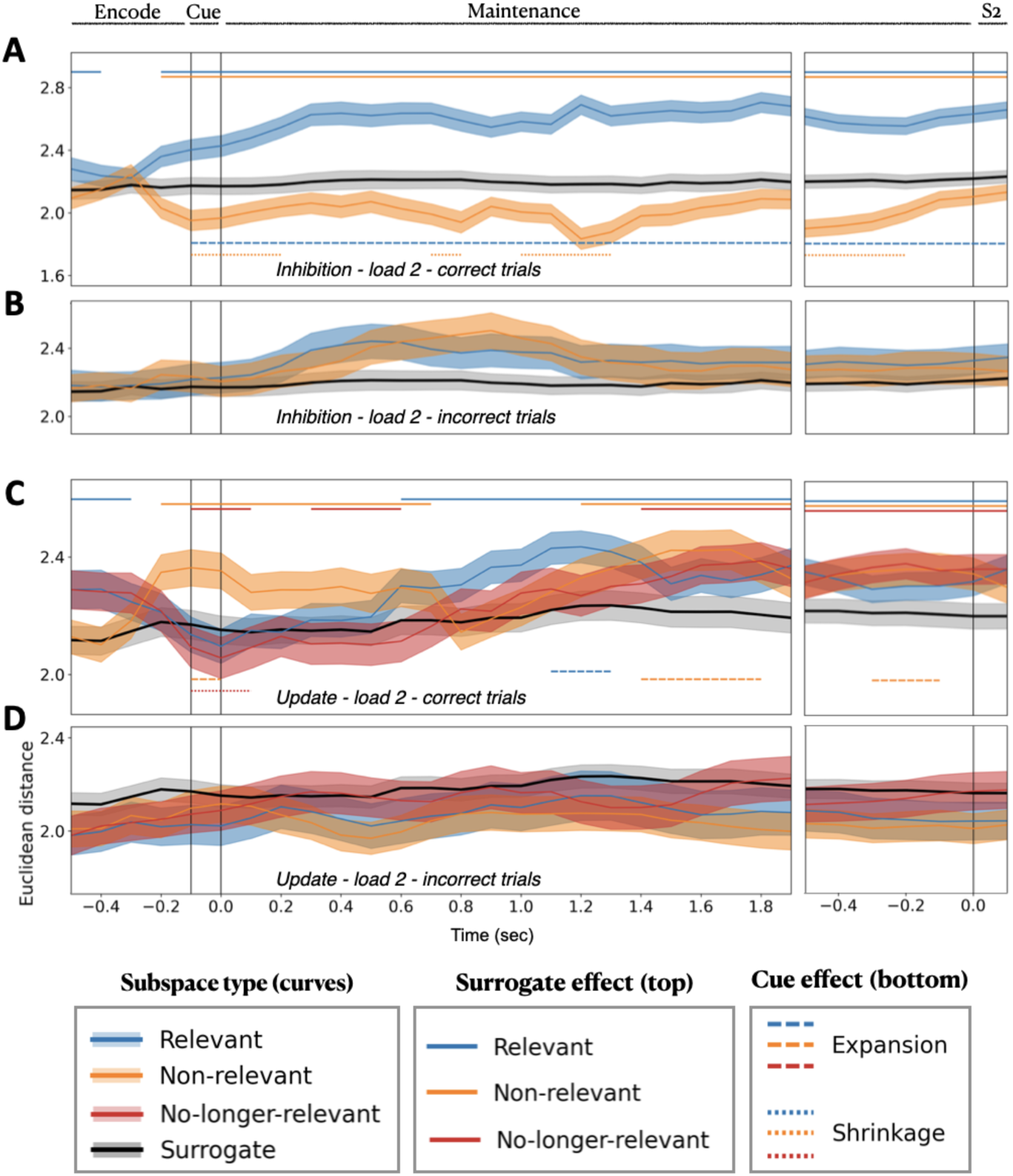
Effect of performance. Separability of memory representations within feature-specific neural subspaces for stimulus load 2, shown separately for correct and incorrect trials. A–B) Inhibition condition (A: correct trials; B: incorrect trials). C–D) Update condition (C: correct trials; D: incorrect trials). The y-axis scales are identical within A–B and C–D, respectively, to facilitate comparison between correct and incorrect trials. Panel layout, time axis, and significance markers follow the same conventions as in Figure 2.

**Table 1.** Summary of trial counts by condition and load. Values represent the mean ± standard deviation and range across participants for the total number of trials, as well as the number of correct and incorrect trials, separately for each condition and load.

| Condition | Valid trials | Correct trials | Incorrect trials |
| --- | --- | --- | --- |
| Control - load 1 | 149.4 ± 5.2 [128, 152] | 123.2 ± 14.7 [93, 148] | 26.3 ± 13.4 [4, 52] |
| Control - load 2 | 149.4 ± 5.2 [128, 152] | 102.7 ± 13.1 [75, 130] | 46.7 ± 11.6 [22, 70] |
| Inhibition - load 1 | 298.9 ± 10.4 [256, 304] | 257.1 ± 28.5 [172, 298] | 41.7 ± 26.1 [3, 116] |
| Inhibition - load 2 | 298.9 ± 10.4 [256, 304] | 226.9 ± 27.6 [165, 278] | 72.0 ± 25.2 [24, 123] |
| Update - load 1 | 298.9 ± 10.4 [256, 304] | 266.8 ± 37.0 [76, 301] | 32.1 ± 34.4 [3, 212] |
| Update - load 2 | 298.9 ± 10.4 [256, 304] | 218.4 ± 30.9 [151, 280] | 80.5 ± 28.6 [24, 137] |

#### Shape metrics

To further characterize the temporal dynamics of neural subspaces (hypothesis 3.2), we computed the Procrustes distance between the subspaces at every pair of time points, yielding a time-generalization matrix. In this context, the Procrustes distance quantifies changes in the global geometry (i.e., subspace shape) of the neural subspaces over time after removing differences due to rotation, reflection, and translation. Larger distances indicate greater changes in subspace shape between time points. We observed that the inhibition and update conditions showed higher Procrustes distances after the cue compared with the control condition, suggesting stronger shifts in the global geometry of subspaces, whereas the control condition remained more stable (Figure 4A–B). When directly contrasting inhibition and update, we found higher distances in inhibition during the second half of the maintenance period (Figure 4C). These reconfigurations were cue-driven, as no differences were observed before the cue. When examining subspaces by cue-driven relevance, a similar pattern emerged, with control subspaces remaining more stable than other conditions after the cue, regardless of whether they were compared with relevant or non-relevant subspaces (Figure S9). These results indicate that the expansion and shrinkage patterns observed in separability (Euclidean distances in PC scores) were not driven by misalignment of subspaces across time. Instead, the elevated Procrustes distances in the inhibition and update conditions suggest that cue-driven processing was accompanied by broader reconfigurations of representational geometry, reflecting changes in the overall shape of neural subspaces beyond simple increases or decreases in separability. Together, these findings suggest that prioritization and updating of WM representations involve not only changes in the distances between representations but also a restructuring of their global geometric organization.

**Figure 4.**
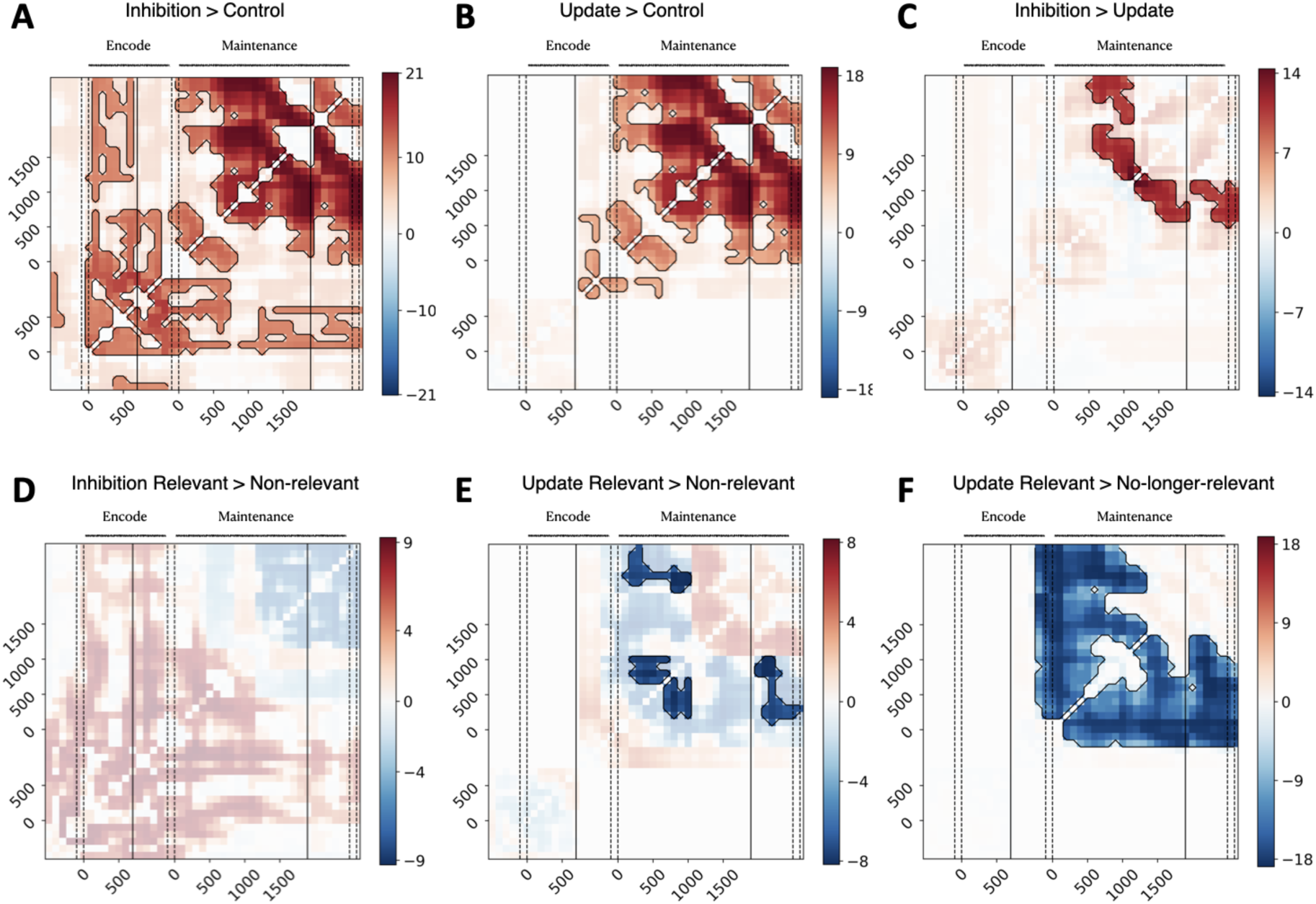
Time-generalization of Procrustes distances, showing t-statistics for multiple contrasts with cluster-based permutation correction for multiple comparisons. Red clusters indicate significant positive differences (e.g., A > B means distances are higher in condition A than B, reflecting greater stability of the global configuration in B), whereas blue clusters indicate significant negative differences. Highlighted colors show corrected significant effects, while the overlaid background displays uncorrected values. Panels correspond to the contrasts: A) Inhibition > Control, B) Update > Control, C) Inhibition > Update, D) Inhibition Relevant > Non-relevant, E) Update Relevant > Non-relevant, and F) Update Relevant > No-longer-relevant. Vertical dashed lines indicate onset and offset of S1, cue, and S2. Time is locked to the onset of the encode and maintenance periods, respectively. Black vertical lines mark block boundaries corresponding to different jitter-corrected delay durations (encode: 700–1000 ms; maintenance: 1500–2000 ms).

### Oscillatory vs aperiodic geometry

The task-related modulation of neural subspace geometry described above, which is based on the analysis of broadband source data, was re-examined using spectrally decomposed data to test for a contribution of narrowband oscillatory versus aperiodic dynamics (hypothesis 5). Broadband data were decomposed into three data-driven frequency bands, namely theta (2.00–5.75 Hz), alpha (6.46 – 13.06 Hz) and beta/gamma (14.69 – 60.00 Hz) (see Methods for details about subject-level and group-level boundaries on data-driven frequency bands). Neural subspaces were estimated for each frequency band, computing the so-called narrowband (NB) separability. We then predicted our main findings on separability (Figure 2) from NB separabilities (Figure 5). Theta-band separability was predictive across conditions (all conditions: t = 11.9, p < 0.0001; all individual conditions p < 0.0001), whereas alpha was predictive in the control (t = 5.0, p < 0.0001) and update conditions (t = 3.0, p = 0.006); while beta/gamma was predictive in update condition (t = 3.2, p = 0.003) (Figure 5A, model 1). Next, we extended the model by including an shared broadband separability predictor (Figure 5B, model 2). This predictor was derived by applying PCA to the NB separabilities and extracting a broadband-like PC. NB predictors were then residualized with respect to this broadband component. In this model, the shared broadband predictor showed a strong positive effect (all conditions: t = 12.6, p < 0.0001; all individual conditions p < 0.0001). Among NB predictors, only theta remained significant (all conditions: t = 3.9, p = 0.0006; inhibition: t = 5.9, p < 0.0001; update: t = 2.9, p =0.01). In contrast, alpha and beta/gamma predictors were no longer predictive, except for the negative association in inhibition condition (t = –3.4, p =0.002). In both models, covariates of no interest were significant. Specifically, load showed a strong effect across all conditions (all conditions: t = 6.8, p < 0.0001; all individual conditions p < 0.0001), as well as stimulus feature (all conditions: t = 3.04, p = 0.006; inhibition: t = 4.4, p = 0.0002). Visual inspection of the separability time courses further supported the linear modeling results (Figure S10-12). The temporal dynamics of separability for the shared broadband component and the theta band closely matched the task-related separability patterns observed in our main analyses, consistent with the linear model results. In contrast, although separability in the alpha and beta/gamma bands also varied over the course of the task, their temporal profiles did not consistently correspond to the separability dynamics observed in the main analyses.

**Figure 5.**
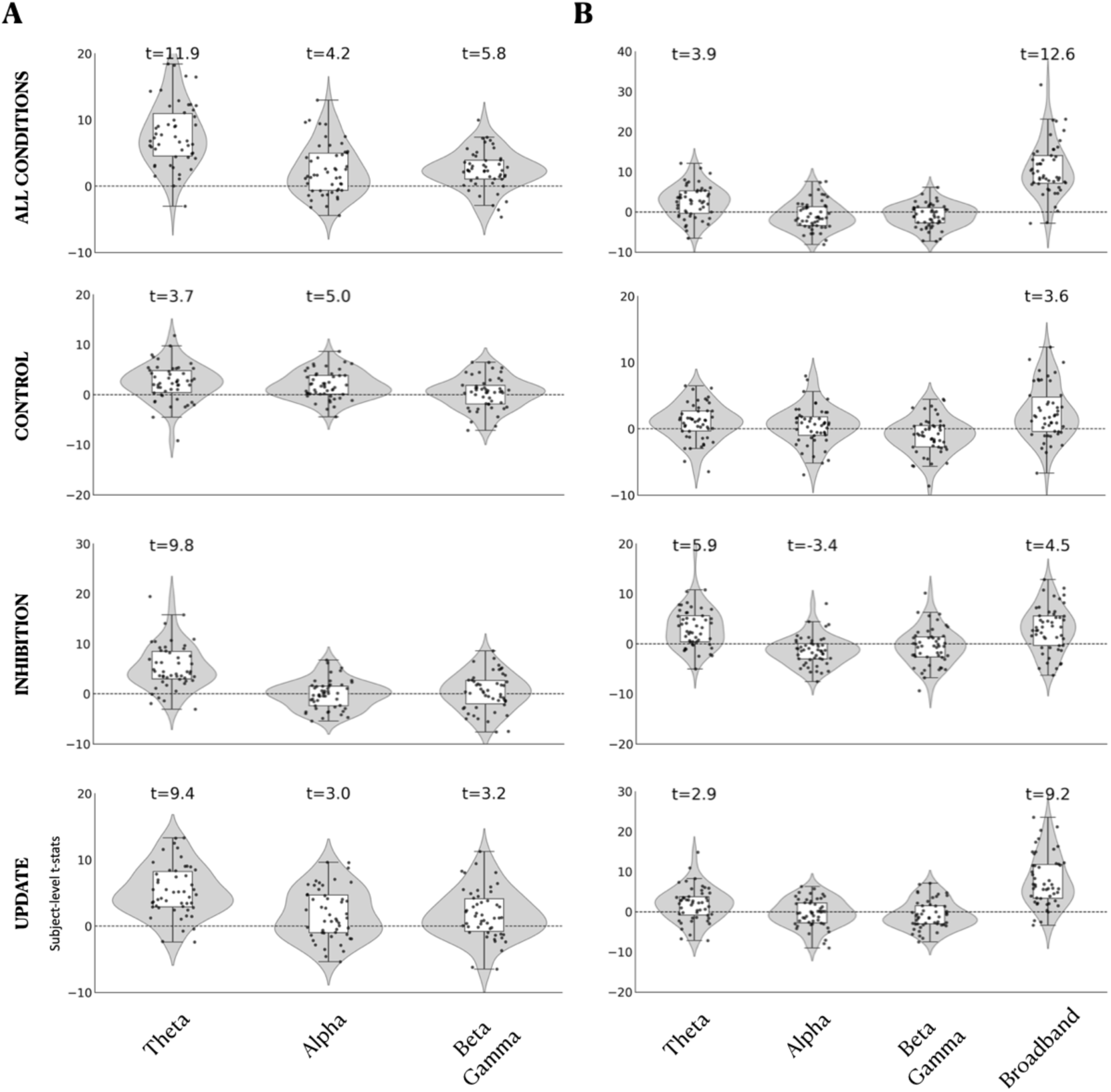
Oscillatory vs broadband geometry. Results from hierarchical linear models predicting our main findings on separability from narrowband oscillatory components (A, model 1), and from shared broadband component and residualized narrowband oscillatory components (B, model 2). Results derived from the real part of the complex Morlet wavelets, and frequency bands defined based on clustering-derived frequency boundaries at the subject level. Predictor variables are shown on the x-axis. First-level t-statistics are shown on the y-axis, and second-level t-statistics are reported in the text for regressors that remained significant after FDR correction (p < 0.05, two-tailed).

To further evaluate the influence of subspace relevance, we repeated the analysis after partitioning separability into relevant and non-relevant subspaces following the cue, fitting the same model separately to each subset (Figure S13). Shared broadband separability remained a significant predictor in the inhibition condition for both relevant (t = 3.2, p =0.007) and non-relevant subspaces (t = 3.3, p =0.01), and in the update condition for non-relevant subspaces (t = 8.1, p < 0.0001). In contrast, theta separability was no longer significant in the inhibition condition after separating subspaces by relevance. In the update condition, however, theta separability remained significant in the relevant subspaces (t = 3.1, p =0.01).

The results described above were obtained using subject-specific frequency band boundaries (Figure 5) but using group-level frequency bands yielded qualitatively similar results (Figure S14). In addition, the reported analyses were based on spectral decomposition of the real part of the complex Morlet wavelet time series, thereby preserving both phase alignment and power fluctuations (Figure 5). When narrowband signals were instead derived from amplitude envelopes, the same overall pattern of results was observed (Figure S15). Finally, to assess whether the results were influenced by performing hyperalignment separately for each frequency band, we repeated the analysis using the hyperalignment transformation derived from our main analysis. This manipulation produced no qualitative changes in the results, but showed stronger statistical evidence (Figure S16).

### Supplementary analyses

The main findings were robust to a range of analytical choices, including the use and implementation of hyperalignment, the dimensionality of the hyperalignment space and neural subspaces, brain parcellation resolution, and source-modelling procedures. Although some effects were reduced or no longer detectable under alternative configurations—particularly in higher-dimensional subspaces or without hyperalignment—the overall pattern of task-related changes in subspace separability was largely preserved. These analyses indicate that our findings are not driven by a specific choice of analysis parameters or preprocessing pipeline, while also showing that hyperalignment and fidelity weighting enhance sensitivity to subtle, low-dimensional changes in subspace geometry. Detailed results of these robustness analyses, including the corresponding statistical results and figures, are provided in the Supplementary Material (Figure S17-33).

## DISCUSSION

We show here that low-dimensional subspaces organizing memory representations are dynamically shaped by task demands. First, subspace alignment (e.g., principal angle between subspaces) increased under high cognitive demands, suggesting that subspace orthogonalization is recruited selectively when required rather than serving as a general organizational principle (hypothesis 2). Second, separability between memory representations within those subspaces was also modulated by task demands (hypothesis 3). Specifically, subspaces expanded when items became task-relevant and shrunk when items became task-irrelevant. We further confirmed that these representations were low-dimensional (hypothesis 1), as most of the variance was captured by only a few components (here, three PCs). Importantly, the task-dependent modulation disappeared when subspaces were defined using a larger number of PCs, indicating that the effect was specific to the low-dimensional structure (hypothesis 1). The modulation was behaviorally relevant (hypothesis 4), as it was present in correct trials, whereas there was no evidence of modulation in incorrect trials. The main effects were also robust to variations in preprocessing, indicating that the observed effects were stable across different approaches. Finally, we found that task-dependent modulation of subspace dynamics was explained by slow oscillatory activity in the delta–theta frequency range (hypothesis 5). In addition, however, shared broadband activity made a substantial independent contribution, indicating that subspace geometry is shaped by both oscillatory and broadband components of the neural signal.

Our findings of low-dimensional and behaviorally relevant WM-related subspaces aligns well with previous studies of intracranial recordings in non-human primates ^18–20,71^ as well as non-invasive MEG studies in humans ^21^. We extend this work by showing that prioritization and manipulation of memory contents (inhibition and updating) actively reshape subspace geometry. This is consistent with ^18^, who reported that deprioritizing one item increases the alignment between neural subspaces, making them less orthogonal (i.e., more parallel). However, such effects could reflect a passive consequence of releasing active maintenance constraints rather than an active reconfiguration of representational geometry. In contrast, our results suggest a functionally meaningful reorganization: prioritization selectively expands relevant subspaces while compressing irrelevant ones, a pattern that may enhance robustness to neural noise and improve representational separability. Importantly, we show that these effects emerge as time-resolved, within-trial dynamics rather than solely as static differences across conditions or memory loads. This suggests that subspace geometry is continuously modulated by cognitive operations on behaviorally relevant timescales. More broadly, these findings resonate with classical WM models based on attractor dynamics, where the separation and stability of attractor states directly shape memory performance ^22,23^, as demonstrated in single-item WM tasks using intracranial recordings in primates ^7^. Our results further support the idea that cognitive operations— and computation more generally—are implemented through dynamic reconfiguration of neural manifolds, potentially by rerouting population trajectories within these low-dimensional spaces ^27,28^. Our findings also generate a testable prediction. If cognitive operations in WM are implemented through transformations of WM-related subspaces, as suggested here, then perturbation approaches such as TMS or tDCS/tACS that improve behavioral performance should produce specific and constrained changes in subspace geometry. In particular, performance improvements should be associated with greater orthogonality between subspaces and/or increased separability of memory representations within each subspace, thereby enhancing the robustness and discriminability of neural codes. In contrast, interventions that impair performance may produce a broader range of geometric disruptions. Because successful readout by downstream neural populations likely depends on an optimal balance of dimensionality, alignment, and representational separation, performance deterioration could arise from multiple forms of subspace disorganization rather than a single stereotyped pattern. Future studies could test these predictions directly by combining causal neural perturbations with analyses of subspace geometry.

We also extend previous literature by showing that WM-related subspaces support the simultaneous maintenance of different stimulus features, not just multiple items of the same feature, as described previously ^18–21,29,71^. However, we found that orthogonality between subspaces encoding different features was not a default organizational property, but emerged only under increased cognitive demand. The alignment between feature-specific subspaces, reflecting the degree to which they are orthogonal versus parallel, was quantified using principal angles (PA) and variance accounted for (VAF). Neither measure exceeded chance levels estimated from surrogate data. This finding contrasts with previous MEG studies of sequential working memory, in which subspaces encoding spatial locations across ordinal positions in sequences of three and four items showed significantly above-chance alignment throughout the entire delay period ^21^. Our findings suggest that any apparent orthogonality between orientation and shape representations likely reflects sensor-level mixing of signals due to linear projection of partially distinct neural sources, rather than a structured representational organization at the level of neural population codes. From a coding perspective, this is expected: orthogonal subspaces are primarily required when multiple items are represented within the same or overlapping neural populations to prevent mutual interference. In contrast, when representations are supported by largely distinct neuronal populations, such orthogonalization is not strictly necessary. Consistent with this interpretation, we observed PA values near ∼60°, indistinguishable from a null distribution of random projections, suggesting quasi-orthogonality driven by mixing at the sensor level rather than structured neural alignment. Nevertheless, subspace orthogonality increased following the cue in the control condition, but only under high memory load (two items per feature). This indicates that subspace geometry is not fixed, but flexibly engaged as a function of task demands. We speculate that this flexibility reflects the involvement of shared frontoparietal and attentional control systems that support multiple feature representations, leading to context-dependent interactions between otherwise partially segregated neural populations.

Our findings contribute to the debate of the computational role of oscillatory vs aperiodic activity (hypothesis 5). We observed that WM-related subspaces emerge from a combination of shared broadband activity and theta oscillations. The shared broadband component explained the largest proportion of the geometric effects. This aligns with prior reports of WM-related subspaces in spiking activity ^18,19^ and broadband source-reconstructed MEG data ^21^, and supports a contribution of broadband, non-oscillatory activity to the geometry of neural subspaces. However, an important limitation is that the shared broadband component identified here is only an approximation of the conventional aperiodic (1/f) component of neural signals ^56^. Thus, our results should not be interpreted as demonstrating a specific contribution of the canonical 1/f component per se. Rather, they indicate that the broadband, non-oscillatory component isolated by our analysis makes a substantial contribution to WM-related representational geometry. The second strongest contributor to the geometric effects was the theta oscillatory component, which showed a pattern similar to that of the shared broadband component, with a positive effect that was consistent across task conditions and irrespective of subspace relevance. This finding aligns with intracranial findings in nonhuman primates showing that theta–spike interactions support memory subspaces encoding ordinal sequence positions ^29^. Our findings extend this observation by suggesting that theta-related geometry is a more general feature of WM representations rather than being restricted to sequential memory tasks. In contrast, alpha and beta/gamma components were not associated with subspace geometry. This may appear surprising given the established role of alpha oscillations in functional inhibition ^41,42,72^ and WM maintenance ^44,48,50^; as well as evidence linking beta and gamma oscillations to the maintenance, readout, and expression of WM representations ^8,9,31,36,49,51,52,73,74^. However, representational geometry captures the organization and separability of mnemonic states rather than the amount of information they contain or the mechanisms governing their access and maintenance. Under this interpretation, alpha oscillations may regulate which representations are prioritized or protected from interference, whereas beta and gamma oscillations may support the activation, maintenance, and communication of mnemonic content. These processes could modulate the strength, precision, or signal-to-noise ratio of neural representations of memory contents without systematically increasing or decreasing distances between memory states and, therefore, without strongly determining the large-scale geometry of the representational space recovered by our analyses. Importantly, these effects may also depend on the aspect of oscillatory activity considered. Our analyses focused on amplitude information from parcel-level source-reconstructed, frequency-filtered time series, and therefore did not directly capture large-scale synchronization between brain regions. Although dimensionality-reduction approaches applied to amplitude signals can capture patterns related to functional connectivity based on amplitude envelope correlations —because shared directions of variance reflect coordinated fluctuations across regions— this correspondence does not extend straightforwardly to phase-based synchronization. In particular, our analyses do not address whether the geometry of WM representations is also expressed in the coordination of oscillatory phase across distributed regions. It is therefore possible that faster oscillations contribute to the geometry of phase-based representational subspaces even when their influence is less apparent in the geometry derived from amplitude fluctuations. Future studies combining geometric analyses of amplitude and phase-based connectivity, ideally across multiple frequency bands, could determine whether oscillatory synchronization provides an additional mechanism through which memory representations are organized and communicated across distributed neural populations. In summary, our results suggest a functional dissociation across spectral components in WM. Aperiodic activity and theta oscillations were the primary contributors to representational geometry, indicating that they may define the large-scale organization and separability of mnemonic states. In contrast, faster oscillations were not associated with geometry despite their well-established roles in attentional control and memory maintenance, suggesting that they contribute to WM through mechanisms other than shaping representational structure. We therefore propose that aperiodic activity and theta oscillations provide a content-independent scaffold for WM representations, whereas faster oscillations support the encoding, maintenance, and selective expression of mnemonic content within this scaffold, potentially through cross-frequency interactions across distributed cortical networks ^33,34^. Future decoding and connectivity analyses will be required to test whether slow and aperiodic activity define the geometry of representational state space, while faster oscillations contribute content-specific information and modulate representational gain.

The current study also contributes to the literature on the distinction between passive maintenance (short-term memory, STM) and active manipulation of memory contents (working memory, WM) ^1,75,76^. Our finding on the expansion of relevant subspaces (increased Euclidean distance) after the cue in both control condition (maintenance of memory contents without manipulation) as well as inhibition/update conditions (maintenance and manipulation of contents) supports the dynamic code of WM, even when there is no task-induced manipulation of memory contents ^14,16,17,77^. These findings might suggest that no qualitative changes operate at the level of neural geometry between passive maintenance and active manipulation of memory contents. However, separability offers a partial characterization of subspaces. It has recently been suggested that shape metrics provide a more complete characterization of subspace geometry ^69^. Here, we used Procrustes distance to quantify the global geometry of subspaces, and observed that subspaces were indeed more stable in the control condition than in the inhibition and update conditions. This difference held whether we compared control subspaces against all inhibition/update subspaces or only against the task-relevant ones, indicating that the effect was not driven by non-relevant subspaces. This finding suggests that active manipulation of memory representations induces a stronger reorganization of subspace geometry. Specifically, manipulation is associated with a more dynamic reconfiguration of neural coding (inhibition and update conditions), whereas maintenance alone relies on relatively more stable representations (control condition). These results are consistent with previous studies ^3^. Using a WM task designed to control for confounds such as strategy use and domain-specific skills, ^78^ showed that phase–amplitude cross-frequency coupling distinguishes maintenance from manipulation processes. Manipulation was characterized by stronger and more broadly distributed network activation than maintenance. In a study of intracranial EEG in humans, ^79^ observed that WM prioritization was associated with a transformation in the format of WM representations. While category-specific representations in ventral visual cortex remained stable across encoding and maintenance, representations in prefrontal cortex were reorganized into a distinct task-dependent format during maintenance, providing evidence that prioritized memory contents are supported by dynamic representational transformations rather than static neural codes. Studies in non-human primates further support the distinction between passive storage and active manipulation. ^80^ showed that memory representations in lateral prefrontal cortex morphed dynamically after the presentation of a distractor, ^18^ observed that prioritization and control in WM lead to reconfiguration of low-dimensional population subspaces, and ^8,9,51^ reported that manipulation or flexible use of memory contents increases dynamics in prefrontal and parietal population activity, often through oscillatory bursts. In summary, our findings are consistent with previous evidence that manipulating memory contents involves greater neural dynamics than passive maintenance, and extend these findings by suggesting that such dynamics reflect the reorganization of neural subspace geometry.

Another notable finding concerns the temporal dynamics of subspace reconfiguration. We observed that the timing of subspace expansion and shrinkage differed across task conditions. In the control condition, task-relevant memory representations expanded immediately following the cue. In contrast, both inhibition and update conditions showed a delayed onset of expansion. In the inhibition condition, relevant expansion was preceded by the shrinkage of non-relevant subspaces encoding the deprioritized feature domain; while in the update condition, relevant expansion was preceded by the shrinkage of no-longer-relevant subspace, encoding the outdated stimulus of the prioritized feature. Critically, the delay in expansion of relevant subspace was more pronounced in the update condition, around 800 ms after cue offset, and it was closely coupled to the transient shrinkage of the no-longer-relevant subspace. We speculate that this temporal dissociation reflects the multi-stage organization of WM prioritization, in which changes to memory content precede the prioritization of updated information. This interpretation is consistent with influential models proposing that WM operations unfold in temporally distinct stages, where the selection of memory contents precedes their transformation into task-relevant representations ^81,82^. In the current study, we observed that these stages are reflected in dissociable effects on the geometry and dynamics of neural subspaces. Moreover, the cue-induced modulation of separability between non-relevant and no-longer-relevant subspaces did not persist throughout the entire maintenance period but instead returned toward pre-cue levels close to the response probe. This pattern suggests a release from active inhibition once updating is completed, consistent with transient attentional modulation of WM representations. This interpretation aligns with evidence against sustained attentional effects during the delay period in WM tasks ^81,83,84^.

Some limitations should be taken into consideration. First, we focused our attention on the geometry of subspaces encoding different stimulus features. However, we did not address whether such subspaces coexist with subspaces encoding several items of the same stimulus feature. In the current study, we could define subspaces based on the spatial location of stimuli, since all items appeared in one out of four predefined spatial locations. Unfortunately, the information of the precise spatial location of each stimulus was not recorded during data collection, preventing us from testing the interplay of subspaces within– and between-features. This is a future direction, we hypothesise that the non-orthogonal subspaces between feature domains that we observed should coexist with orthogonal subspaces within each feature domain, specially when two gratings and two polygons are presented simultaneously. We also expected the degree of orthogonality between within-feature subspaces, quantified by alignment metrics, would be behaviorally relevant, as observed previously^21^. Future studies should explore the hierarchical organization of subspaces. Second, our findings are constrained by choices made during preprocessing and the geometric analysis pipeline. To ensure that our results were not driven by these methodological decisions, we systematically varied key preprocessing and analysis steps and confirmed that the main effects remained stable, demonstrating the robustness of our findings. The largest deviations from the main results were observed when source time series were computed without phase-based fidelity weighting ^85^. In this case, we still observed comparable effects on subspace separability, but the timing of subspace expansion and contraction was slightly shifted, suggesting a contribution of phase information to the neural dynamics underlying WM-related subspaces.

In conclusion, working memory representations of multiple items and features are organized within low-dimensional neural subspaces, whose geometry is flexibly reshaped by task demands. Prioritization expands these subspaces, while de-prioritization shrinks them, suggesting that cognitive operations on memory contents are implemented through changes in subspace geometry. Subspaces representing items from different feature domains are generally oblique, becoming more orthogonal only under high cognitive demands, which suggests that orthogonalization is recruited when necessary. Finally, subspace dynamics are largely driven by the interaction between aperiodic neural activity and slow oscillatory activity in the theta frequency band.

## METHODS

### Sample description

We recruited fifty healthy, right-handed participants (28 females; age = 24.66 ± 4.77 years) who completed two sessions of a WM task while their brain activity was recorded simultaneously with MEG and EEG. All participants had normal or corrected-to-normal vision, no history of psychiatric or neurological conditions, and were not taking medications affecting the central nervous system. Participants with any contraindications for MEG, MRI, or TMS were excluded (TMS was an integral part of a follow-up experiment with these same individuals, the data from which will not be analysed here). Ethical approval of the study was obtained from the Ethical Committee of the University of Glasgow, and all procedures were carried out in accordance with the Declaration of Helsinki.

### Task design

Participants completed a retrocued, multi-item, multi-feature delayed match-to-sample task (Figure 1A). A fixation cross was present at the centre of the screen during the whole task. On each trial, participants were presented a set of stimuli (S1) for 100 ms against a grey background. There were two types of stimuli: polygons and gratings. Polygons were uniquely drawn on each trial to have either 3, 4, or 5 vertices. Each polygon consisted of thick black borders enclosing a white area and the surface of each stimulus was in equal proportions black and white. Gratings were black and white circular patches with a spatial frequency of 1.5 Hz, with 1 of 9 orientations (from 0° to 160° in 20° steps). In S1, participants were presented with polygons only, gratings only, or both types of stimuli simultaneously. For the current study, we only analysed trials with two stimulus features presented at the same time. Memory load was manipulated by varying the number of exemplars of each stimulus feature. Participants could see either one or two exemplars per feature (e.g., 1 polygon and 1 grating; or 2 polygons and 2 gratings). Importantly, when both stimulus features were presented, the number of exemplars was always matched—participants never saw 1 polygon with 2 gratings. Throughout the manuscript, we refer to the number of stimuli per feature simply as load, with two levels: load 1 and load 2. Stimuli could appear in 4 fixed locations above, below, to the right of and to the left of the fixation cross, and positions were randomized across trials. Participants were instructed to commit the identity and the location of the presented stimuli to memory.

After the offset of S1, the screen remained blank for a variable length of time (700/800/900 or 1000 ms), this was the ‘encoding period’. After this period the retrocue was presented: the black fixation cross at the centre of the screen changed colour to either red, green, or purple for 100 ms. The meaning of the cue colors was counterbalanced across participants and explained before the experiment. Cue color indicated the task condition. In the Control condition, participants were required to maintain all stimuli from S1 in memory (purple cue; Figure 1A). In the Inhibition condition, the cue specified which stimulus feature was relevant and which became non-relevant and should be ignored, as it would not be tested in S2 (red cue = remember gratings; green cue = remember polygons; Figure 1A). In the Update condition, the cue had the same meaning as in the Inhibition condition, but was presented alongside an additional stimulus of the relevant feature. Participants had to update the original stimulus in that position with the one presented with the cue (Figure 1A). Hence, in the Update condition, following the cue, stimuli could be classified into three categories: relevant, the stimulus feature that participants were required to maintain after the cue; no-longer-relevant, the stimulus belonging to the relevant feature that was replaced by the new stimulus presented with the cue and should now be ignored; and non-relevant, the stimulus feature that was cued to be ignored. For updating, gratings changed by 90° in orientation from S1 to S2, whereas polygons were replaced with a new polygon that differed in the number of vertices from the original (e.g., a polygon with 4 vertices could be updated to one with 3 or 5, but not 4).

After a maintenance period jittered between 1500 and 2000 ms during which the screen remained blank, participants were presented with a second stimulus set (S2). The task of the participants was to identify whether S1 and S2 were the same, and respond ‘same’ or ‘different’ by pressing the button on the response box at their right index finger or their left index finger, respectively. S2 could be completely identical to S1, in which case participants had to respond ‘same’. Whenever there was a difference between S2 and S1, the correct response depended on the condition. On Inhibition and Update conditions, participants had to respond ‘same’ to S2 if all relevant stimuli (e.g., gratings) were the same as in S1 even if one of the irrelevant stimuli (e.g., polygons) was different. Participants only had to respond ‘different’ if there was a difference in the relevant stimulus set. In the Update condition, participants only had to respond ‘same’ if the relevant stimuli were identical to S1 with the exception of the updated one during the cue. S2 had to contain the updated stimulus (not the original one) for a ‘same’ response to be correct. On Control condition, participants had to respond ‘different’ if any of the stimuli in S2 differed from S1. S2 remained on screen until the participants provided a response. Reaction time and accuracy were recorded. The following trials then started after a variable inter-trial interval with a duration between 1750-2250 ms. Participants performed a total of 19 blocks in total (across two sessions), with 112 trials each, for a grand total of 2,128 trials.

### Behavioral analysis

Behavioral accuracy was quantified as the proportion of correct responses across all experimental conditions and memory loads (Table 1). To determine whether accuracy exceeded chance, a one-sided binomial test was conducted for each participant, with the null hypothesis that accuracy was equal to 50% (chance level). P-values below 0.05 were considered statistically significant. Binomial tests were performed on all trials pooled across conditions, as well as separately for each load (one or two), for each condition (control, inhibition, or update), and the combination of condition and load.

To test whether behavioral accuracy differed across conditions and load levels, we fitted a linear mixed-effects model. The dependent variable was each participant’s mean accuracy (proportion of correct responses) for each condition and load combination. Condition (control, inhibition, update) and load (1 vs. 2) were included as fixed effects, and subject was included as a random intercept to account for repeated measurements. Post-hoc pairwise comparisons between conditions (update vs. inhibition) were conducted using Tukey’s Honestly Significant Difference (HSD) test. To further examine the influence of which stimulus feature remained relevant after the cue, a second model was fitted including only the inhibition and update conditions, with an additional predictor specifying the relevant feature. This analysis did not include control trials, as all stimulus features remained relevant after the cue in that condition.

### MEG acquisition and preprocessing

M/EEG data was collected over 2 sessions, each roughly 5 hours in duration. Participants then completed a variable number of blocks of the WM task inside the magnetically shielded room (MSR) of the MEG, with the aim of completing a total of 19 blocks across the two sessions. Session 1 also included one practice block. In every block, stimuli were back projected onto a screen 115 cm away from the head of the participant, using a PROPixx projector located outside the MSR. Participants were offered frequent breaks, during which they could leave the MSR. After the final block, participants washed their hair (if needed) and were debriefed. In a separate session, structural MRI data was collected from participants for source reconstruction. This session lasted approx. 1 hour, including safety screening and the actual scan itself.

Participants’ brain activity was recorded concurrently with a 64-channel EEG cap and 306-channel MEG (204 planar gradiometers, 102 magnetometers), at a sampling rate of 1000 Hz. To capture ocular artefacts, bipolar horizontal EOG channels were placed next to the outer canthi of each eye, and bipolar vertical EOG channels were placed above and below the right eye. The EEG was referenced to an electrode placed on the right mastoid during recording. An electrode on the left cheekbone served as ground. Electrode impedances were kept below 5 kΩ. Five head position indicator (HPI) coils were placed on the electrode cap for continuous head position tracking in the MEG. The locations of three fiducial points, the HPI coils, the electrodes, and additional points defining head shape were digitised using a Polhemus FASTRAK digitiser (Polhemus LTD, Vermont, USA).

MEG and EEG data were preprocessed offline using MNE-Python ^86^. First, bad channels were identified through visual inspection for both types of data. On average, 4.15 ± 2.50 channels were marked for EEG, and 4.56 ± 1.77 for MEG. Then, head motion correction was applied to the MEG data using the continuous HPI recording, followed by Maxwell filtering with temporal signal space separation to suppress extracranial noise, and to interpolate bad channels. Spherical spline interpolation was used for bad channels in the EEG. EEG data was also re-referenced to the average of all electrodes. Both types of data were then low-pass filtered with a cut-off of 270 Hz, and notch-filtered to attenuate 50 Hz line noise and its harmonics. Data across blocks within a given session were then concatenated.

Independent component analyses (ICA) were then applied to MEG and EEG data separately, using the FastICA algorithm (Hyvärinen and Oja, 2000). For ICA only, the data were downsampled to 200 Hz and band-pass filtered between 1-30 Hz. Ocular and heart-beat artefacts were then removed from the original data sets following visual inspection of independent components. On average, 2.83 ± 1.19 components were removed for EEG, and 2.97 ± 1.07 for MEG. Data was then downsampled to 500 Hz.

T1-weighted MRI scans were preprocessed using Freesurfer (http://surfer.nmr.mgh.harvard.edu/). This included automatic volumetric segmentation, surface reconstruction, and cortical parcellation into 400 parcels forming 7 networks, labelled in accordance with the Schaefer atlas ^68^. Surface-based source space was then created using MNE’s default oct6 spacing. MNE was also used to generate a 3-layer boundary element method (BEM) head model with layer-wise standard conductivity values of 0.3, 0.006, and 0.3, and the corresponding BEM solution. This head model was then coregistered manually with the points digitised at the beginning of each M/EEG session using MNE’s Coregistration GUI to align the MRI, MEG, and head coordinate frames for the calculation of the forward solution. The noise covariance matrix (NCM) was calculated using 1-second-long segments of broadband data from the ITI, filtered between 155-195 Hz. This window was chosen to avoid the notch-filtered frequency bands used for power line artifact removal. The forward solution and the inverse operator were computed using the MNE-Python implementation of the MNE inverse method, together with custom code, and were then used to transform the continuous M/EEG data to source-vertex time series with dipole orientations fixed to the pial surface normals and a 5 mm inter-dipole separation. Source time series were collapsed into the 400 parcels of the Schaefer atlas using fidelity-weighted collapsed inverse operators to maximise parcel reconstruction accuracy in each subject’s source space^85^. Briefly, fidelity weighting was estimated by simulating broadband parcel-level time series, assigning each source within a parcel the same simulated signal, projecting the resulting source activity to sensor space with the forward model, and reconstructing it back to source space with the inverse operator, and quantifying how accurately the reconstructed source and parcel signals matched the original simulated signals after the forward–inverse transformation. Fidelity was quantified using phase-based metrics (complex phase-locking values, cPLV) between the original and reconstructed signals, making the weighting primarily sensitive to preservation of phase relationships rather than signal amplitude. The simulated signals were broadband random complex time series rather than oscillations at a specific frequency, such that the resulting weights reflected broadband reconstruction fidelity and spatial leakage characteristics of the forward–inverse model. The inverse operators were then weighted to maximise the correspondence between the original and reconstructed parcel time series while reducing cross-parcel leakage.

### Geometric analysis of neural subspaces

The geometric analysis of neural subspaces was performed using the approach described by ^21^, who extended to broadband MEG data the geometric analysis methods previously applied to spike activity data ^18,19^ (Figure 1E). We created subject-specific matrices of neural activity X (M by N), where M is the number of conditions and N is the number of channels (400 cortical parcels from ^68^). Conditions corresponded to the stimuli presented during S1 and were defined within two feature spaces: an orientation feature space for gratings and a shape feature space for polygons. The shape feature space comprised polygons grouped by number of sides—triangles, rectangles, and pentagons. For the convex, low-side-count polygons used in this study, the number of sides provides a good approximation of radial frequency, defined as variation in curvature along a shape’s contour and a key feature used by the visual system to encode shape ^87,88^. The orientation feature space consisted of gratings at 0° to 160° in steps of 20°. To increase the signal to noise ratio, we pooled orientations into four bins: two bins of 40° centered around the cardinal directions 0° [160 20] and 90° [70 110], and two bins of 30° centered around the oblique orientations of 45° [30 60] and 135° [120 150]. This grouping was motivated by the visual system’s well-documented preference for cardinal orientations, as observed in both neurophysiological and psychophysical studies ^89^. Each column of X captures the responsiveness of a given parcel (channel) to the stimuli presented across trials, estimated using ridge-regularized linear regression (λ = 1) applied separately to each channel. A fixed regularization parameter was used across channels to avoid channel-specific scaling differences that could arise from data-driven λ selection, given that neural subspaces were computed from the full X matrix. Using alternative regularization values (λ = 0.01, 0.1, and 10) produced no qualitative differences in the main results (data not shown). In the ridge regression, the response variable was the channel activity averaged over a predefined time window (see below), baseline-corrected by subtracting the mean activity during a 0.5-second pre-stimulus baseline period, defined as [-0.6, –0.1] seconds relative to S1 onset. The design matrix included 7 binary predictor variables (4 orientations, 3 shapes), indicating which stimuli were presented on each trial. The resulting beta weights for the M conditions served as the entries in that channel’s column of X. Finally, each column of the X matrix was demeaned prior to geometric analysis. We computed subject-specific X matrices separately for each condition (control, inhibition, and update) and loads (1 or 2). For the inhibition and update conditions, separate X matrices were computed based on the cue-indicated relevance of stimulus features: one matrix for trials in which orientation was relevant and shape was non-relevant, and another for trials in which shape was relevant and orientation was non-relevant. For the update condition, an additional stimulus was presented alongside the cue. This stimulus replaced the item originally presented at the same spatial location in S1 and was the item probed in S2. Accordingly, one X matrix was computed before the cue, as in the other conditions. After the cue, two X matrices were computed, both of which included the feature that became irrelevant following the cue (the non-relevant subspace). The matrices differed in the second subspace they contained. One matrix combined the non-relevant subspace with the feature that remained relevant after the cue with the updated stimulus presented alongside the cue (the relevant subspace). The other combined the non-relevant subspace with the relevant feature but with the original S1 stimulus that had been replaced by the update (the no-longer-relevant subspace). The no-longer-relevant subspace was unique to the update condition and was computed only after the cue. It therefore captured the representation of the feature that remained relevant after the cue, but for the original S1 stimulus rather than the updated stimulus. Across all conditions, X matrices were computed in a time-resolved manner. For this analysis, the data were segmented into overlapping 400 ms windows with 100 ms steps, spanning all trial events. To increase the sample size for the regression models, we combined trials from both MEG sessions, as source modeling and brain parcellation accounted for potential differences in head position between sessions.

To estimate low-dimensional neural subspaces, we reduced the dimensionality of the X matrix along the channel dimension (i.e., its columns) using Principal Component Analysis (PCA). The data were projected into the subspace defined by the top 10 principal components (PCs), yielding the Z matrix of PC scores (M = 7 conditions × p = 10 PCs), which were then used to hyperalign neural subspaces across subjects (see below). After hyperalignment, from the resulting Z* matrix of aligned PC scores (M = 7 conditions × p = 10 PCs), we extracted the neural subspaces by selecting the top k = 3 principal components, resulting in the Z* matrix with dimensions M = 7 conditions × k = 3 PCs. Then, we subset the Z* matrix by feature, creating Z*_orientation (4 conditions by k components) and Z*_shape (3 conditions by k components). Finally, we computed the subspace plane for each feature subspace by taking the two leading components of another PCA on Z*_orientation and Z*_shape separately.

We quantified the geometry of these subspaces using alignment metrics, separability metrics, and shape metrics ^21,69^. Alignment between feature subspaces was quantified using two metrics: the principal angle (PA) and the variance accounted for (VAF). PA was computed as the cosine of the angle between the subspace planes. VAF was defined as the proportion of variance explained by the PCs of one feature subspace when neural data from another feature subspace were projected onto it, providing a measure of similarity in neural geometry across subspaces. Separability within each feature subspace was quantified by the Euclidean distance between memory representations, computed as the pairwise distance between rows of the Z*_<feature> matrix. Shape metrics were used to characterize the temporal dynamics of neural subspaces. Specifically, we computed the Procrustes distance between subspaces (i.e., Z* matrices of aligned PC scores) for every pair of time points, and constructed a time-generalization matrix. Procrustes distances were calculated using the scipy.linalg.orthogonal_procrustes Python function (SciPy), which computes the optimal rotation and reflection (and optionally scaling) aligning two matrices. Given two mean-centered matrices, X_i and X_j (n x m, where n is the number of conditions and m the number of PCs), the Procrustes distance is defined as the Frobenius norm of the difference after optimal transformation:

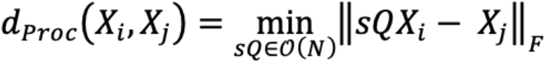

where Q is the orthogonal matrix (m x m) encodes the optimal rotation and reflection, and s is a uniform scaling factor. We computed the Procrustes distance without scaling (s = 1); this distance removes rotation and reflection but preserves the global scale of the subspace, capturing expansions, contractions, and overall geometric reconfigurations over time. Here, we used Procrustes distance as a shape metric to characterize subspace geometry. Procrustes analysis was also used independently for hyperalignment, allowing subspaces to be aligned across subjects, as described in the following paragraph. These two applications are conceptually and analytically distinct.

Hyperalignment. To enable cross-subject comparisons of neural representational geometry, we applied hyperalignment to the PC scores. This procedure aligned the Z matrices of PC scores (M = 7 conditions × p = 10 PCs) across subjects, yielding Z* matrices of aligned PC scores (M = 7 conditions × p = 10 PCs) and projecting all subjects into a common representational space. While we assume that the underlying neural circuitry supporting WM maintenance is conserved across individuals, it may be sampled or expressed differently at the sensor level. To address this, we applied hyperalignment ^69,70^, a method that aligns subject-specific representational geometries using generalized Procrustes analysis ^90^. This approach involves optimizing transformations—including translation, rotation, reflection, and optionally scaling—to minimize differences in the spatial configuration and orientation of neural activity patterns across subjects. By aligning these feature spaces, we recovered a shared low-dimensional structure that enables direct comparison of neural representations across individuals, while accounting for distortions introduced by differences in sampling, signal-to-noise ratio, and spatial localization. We performed hyperalignment on the PC scores derived from the top 10 components, treating each condition as a sample and each PC as a feature. Importantly, we omitted scaling to preserve the relative magnitudes of variance captured by each component. We aligned PC scores with a dimensionality of 10 for two main reasons. First, to ensure consistency across analyses, we wanted to apply a single hyperalignment transformation regardless of the number of dimensions used in downstream analyses. This avoids introducing variability that could arise from performing separate hyperalignment procedures for each dimensionality. Second, using only 3 features for hyperalignment is generally suboptimal, as it may not capture enough of the shared structure across subjects. Conversely, using too many features—such as the full set of 400 channels in the X matrix— risks overfitting and aligning noise rather than meaningful neural signals. A 10-dimensional space provides a balance between preserving signal and avoiding overfitting.

To establish chance-level benchmarks for the geometric measures, we generated surrogate data by jointly shuffling the rows and columns of the X matrices to create null distributions of the geometric variables —alignment (PA and VAF) and separability (Euclidean distance)— following the approach described in ^21^ This procedure was repeated 1,000 times, and the resulting geometric variables were averaged across iterations. For surrogate data, the hyperalignment transformation matrices estimated from the empirical data were applied unchanged to all iterations. This shuffling procedure preserves the global covariance structure and overall variance of the neural data, while selectively disrupting the correspondence between individual stimuli and their positions within the low-dimensional manifold. As a result, the surrogate distributions reflect the geometric structure expected by chance in the absence of stimulus-specific organization.

The main geometric analysis described above was performed using correct trials only, both because correct trials provide a more reliable definition of the representational space and to avoid inflating noise in its estimation. We then asked whether the geometric effects observed in correct trials were also present in incorrect trials (Figure S34). Importantly, this was not a direct statistical comparison between correct and incorrect trials, because stimuli could not be reliably matched across performance conditions, particularly for load-2 trials. We therefore evaluated incorrect-trial subspaces within the representational space defined by correct trials in a two-step procedure, following (Santo-Angles et al. 2025). First, we constructed the X matrices for incorrect trials using the same procedure, but computed their PC scores (Z matrix) by projecting them onto the leading eigenvectors obtained from the PCA of correct trials. Thus, incorrect-trial representations were evaluated within the same representational space that was defined by correct trials. Second, rather than performing hyperalignment separately on incorrect-trial data, we applied the transformation matrices obtained from hyperalignment of the correct-trial data. This procedure ensured that the analysis of incorrect trials was not affected by differences in the definition or alignment of the representational spaces across trial types. Notably, because the PC basis and hyperalignment transformations were derived from correct trials, this approach could, in principle, bias the analysis toward detecting geometric effects in incorrect trials. Therefore, the absence of an effect in incorrect trials provides a conservative indication that the effects observed are not simply a consequence of the analysis pipeline.

We examined the spatial topography of the PCs by quantifying the variance contribution of each component at each cortical parcel. For each participant, time point, and output, the parcel-wise contribution of each of the first three PCs was calculated as the squared eigenvector loading multiplied by its corresponding eigenvalue, thereby quantifying the variance accounted for by each component at each parcel while avoiding dependence on the arbitrary sign of PCA eigenvectors. We focused on the maintenance period and used a hierarchical aggregation procedure to summarize these parcel-wise component contributions across subjects, time points, and conditions. Specifically, we first averaged the component contributions across maintenance time points and outputs within each participant and then summarized these values across participants at each parcel using a one-sample t-statistic. This first analysis was performed on the unstandardized component contributions, allowing us to assess whether the component contributed to the neural representation across parcels in absolute terms, without removing differences in the overall magnitude of their contributions. We subsequently performed a second, complementary analysis after z-scoring the component contributions across parcels within each participant. This normalization removed between-parcel differences in the overall magnitude of component contributions and allowed us to characterize the relative spatial topography of each component, namely which parcels contributed more or less than the participant-specific parcel average. The resulting z-scored values were again summarized across participants at each parcel using an one-sample t-statistic. Positive t-statistic values therefore indicate parcels where the component made a greater-than-average contribution relative to the other parcels, whereas negative values indicate parcels where its contribution was lower than average. Importantly, these hierarchical procedures were used as descriptive frameworks to aggregate and summarize the spatial distribution of component contributions across subjects, time points, and conditions, rather than as a basis for statistical inference or hypothesis testing of the spatial maps. For visualization of the relative spatial topography, we retained the 50% of parcels with the largest absolute group-level t-statistics from the z-scored analysis, while preserving the sign of the t-statistic. This highlighted clusters of parcels showing the strongest relative component contributions without interpreting the threshold as a statistical significance criterion.

We performed a series of supplementary analyses to assess the robustness of our findings to the geometric analysis pipeline. Specifically, we repeated the main analysis under several alternative configurations: (1) without hyperalignment; (2) varying the number of principal components used to subset the Z matrix of PC scores prior to hyperalignment p=[5, 7, 15, 20, 100, 200, 400], instead of the 10 PCs used in the main analysis, (3) applying an iterative hyperalignment procedure in which all subjects were aligned to a single reference subject, repeated across all subjects and averaged across iterations, rather than aligning subjects to the group-average PC scores, as implemented in the main analysis, following ^90^; (4) using different numbers of PCs to estimate neural subspaces (aligned PC-score matrices Z*; k=[2, 4, 5, 6, 8, 10]); (5) varying the number of brain parcels N = [200, 300]; and (6) introducing variations in the source modelling approach. Specifically, we repeated the analysis after removing the fidelity-weighting step from the inverse solution, while keeping all other aspects of the custom pipeline unchanged. In this variant, source estimates were obtained using the inverse operators without reconstruction-based weighting. In addition, we repeated source reconstruction using the standard MNE-Python pipeline, using both MNE and dSPM inverse methods, to evaluate whether the observed effects depended on the choice of inverse solution.

### Statistical analysis

Statistical analyses were performed in a time-resolved manner to assess two effects on the geometry of neural subspaces: cue effect and the surrogate effect. The cue effect was evaluated at each time point during the post-cue maintenance period (from cue offset to S2 onset) by comparing geometric variables (PA, VAF, and separability) to their mean values during the pre-cue encoding period (from S1 offset to cue onset). This analysis tested whether cue-driven prioritization modulates subspace geometry over time. The surrogate effect was assessed at each time point by comparing the observed geometric variables to null distributions derived from surrogate (noise-driven) data, thereby identifying time points at which subspace geometry significantly deviated from chance. Statistical significance was assessed using dependent-samples t-tests, with multiple-comparisons correction applied using the false discovery rate (FDR). Effects were considered significant at a two-tailed, FDR-corrected p<0.05. The behavioral relevance of WM-related subspaces was assessed by testing whether the effects observed in correct trials were also present in incorrect trials. Due to the limited number of incorrect trials in the other conditions, the analysis on separability was restricted to the inhibition and update conditions at load 2. Within these conditions, we tested the same cue and surrogate effects in incorrect trials as those identified in the main analyses of correct trials.

### Oscillatory analysis of neural subspaces

To assess the contribution of oscillatory dynamics to WM–related neural subspaces—originally estimated from broadband source-level time series—we extended our geometric analysis to the time– frequency domain by repeating the same subspace analysis on frequency-resolved data.

First, time–frequency representations were computed at the parcel level by convolving continuous source-space time series with a bank of 30 complex Morlet wavelets. Central frequencies were logarithmically spaced between 2 and 60 Hz, and the number of cycles was fixed to 7 for all frequencies. Wavelet convolution was performed using the CROCOpy toolbox ^91^, yielding frequency-specific analytic signals for each parcel and time point.

Second, rather than using predefined frequency bands, we derived data-driven frequency clusters from inter-parcel functional connectivity ^92^. Connectivity was estimated using orthogonalized amplitude envelope correlation (AEC) and the weighted phase-lag index (wPLI) during the time window spanning from the offset of S1 to the onset of S2. For each subject, we computed a frequency-by-frequency similarity matrix by calculating the cosine similarity between vectorized connectivity matrices for each frequency. The similarity matrices for AEC and wPLI were averaged to form a joint similarity matrix, which was then clustered using the Leiden community detection algorithm to produce subject-level frequency partitions (leidenalg.find_partition python function with resolution = 1.05) ^93^. To derive group-level clusters, a consensus (co-assignment) matrix was computed from the subject-level partitions and clustered again with Leiden (resolution = 0.9 and percentage of subjects with cluster at threshold = 0.4), yielding contiguous frequency bands that were consistent across both metrics and across subjects ^94^. To enable comparison across participants, subject-specific clusters were aligned to the group-level clusters. Specifically, for each subject cluster we identified the group-level cluster with which it showed the largest overlap in frequency bins and reassigned the subject cluster label accordingly (majority-overlap mapping). This procedure preserved subject-specific cluster boundaries while enforcing a common set of cluster labels across participants. Leiden algorithm parameters were selected based on the consistency of the number of clusters across subjects. We tuned the parameters to yield three clusters, as this solution showed the highest stability at the subject level. When parameter settings allowed a larger number of clusters, the number of clusters identified in individual subjects became highly variable, complicating the alignment between subject-level and group-level solutions. For example, when targeting five group-level clusters, the number of subject-level clusters ranged from four to nine. Therefore, we opted for a three-cluster solution, which yields 3.48 ± 0.54 [3, 5] clusters at subject-level. Group-level clusters were as follows: Cluster 1 spanned 2.00–5.75 Hz (n = 10 frequency bins; referred to as theta), Cluster 2 spanned 6.46 – 13.06 Hz (n = 7; referred to as alpha), and Cluster 3 spanned 14.69 – 60.00 Hz (n = 13; referred to as beta/gamma). As a result of the alignment of subject-level clusters to those group-level clusters, we also got the same three clusters, but the precise frequency boundaries of each cluster remained subject-specific. Subject-level clusters are summarized here by their average lower and upper frequency boundaries across participants: Cluster 1 spanned 2.00–6.18 Hz (range across subjects: 2.00–8.17 Hz; number of frequency bins = 10.4 ± 1.8, n = 6–13), Cluster 2 spanned 6.95–12.26 Hz (range: 4.04–16.51 Hz; n = 5.88 ± 2.1, n = 2–9), Cluster 3 spanned 13.78–60.00 Hz (range: 9.19–60.00 Hz; n = 13.68 ± 1.53, n = 11–17).

Third, once data-driven frequency bands were identified, we reconstructed two types of band-limited signals from the Morlet complex timeseries: (a) Complex-sum real signal. For each band, complex Morlet coefficients were averaged across frequencies and projected to the real axis. This preserves both amplitude and phase relationships, so the resulting timeseries reflects phase-aligned oscillatory components within the band. (b) Amplitude (envelope) signal. The amplitude envelopes of each frequency were averaged across the band. Phase information is discarded, so geometry captures only power fluctuations.

Fourth, we applied the geometric analysis to frequency-specific time series described above, focusing on the separability metric, which showed the strongest task-related modulation. For each data-driven frequency band, we computed the separability metric (Euclidean distance) and used two hierarchical linear models to test whether our main findings on separability (Figure 2) was better explained by narrowband (NB) oscillatory components or by shared broadband activity. In model 1, predictor variables included the NB separability metrics. In model 2, predictor variables were the separability derived from an shared broadband component, and the NB separability metrics after regressing out the shared broadband component. Both models also included load and stimulus feature as covariates of no interest. Model 2 predictors were computed as follows. PCA was applied across the separability of NB predictors, and the PC with the most distributed loadings across frequency bands was identified as a proxy for a shared broadband separability component. Specifically, we quantified the distribution of each PC by taking the absolute values of its loadings across frequency bands and computing their sum; the component with the largest summed loading magnitude was selected as the most distributed component. The time series of this component was taken as the shared broadband separability predictor. The original NB predictors were then residualized with respect to this distributed component by regressing each band-specific predictor onto the shared broadband component and retaining the residuals. These residualized predictors were used as the NB predictors in the regression model, thereby capturing variance specific to individual frequency bands after accounting for the shared broadband component. To verify the distribution of all components, we also computed two additional metrics—standard deviation of loadings and the maximum-to-sum loading ratio, with smaller values indicating greater distribution. These metrics were computed per subject and component, and group-level t-tests confirmed that the shared broadband component was significantly more distributed than the remaining components (Figure S35). In all models, we smoothed the separability predictors over time using a Gaussian kernel (σ = 2) within each subject and output condition to reduce the influence of transient fluctuations on the prediction. At the first level of the hierarchical model, we fit both models separately for each subject, treating time points within the maintenance period as independent observations. At the second level, subject-specific regression coefficients were pooled across participants and tested against zero using one-sample t-tests. Models were fit both with all conditions pooled and separately for each condition. Multiple comparisons across predictors were controlled using FDR correction.

