## Supplementary Material for "Working memory operations emerge from dynamic changes in neural subspace geometry"

### SUPPLEMENTARY RESULTS

We assessed the robustness of our results by repeating the geometric analysis under several alternative configurations, focusing on the task modulation of separability. First, when the analysis was performed without hyperalignment across subjects, none of the effects survived correction for multiple comparisons. Nonetheless, uncorrected analyses revealed a significant expansion of relevant subspaces in the Control and Inhibition conditions, whereas no such effect was observed in the Update condition (Figure S17). These effects were absent in incorrect trials (Figure S18). In contrast to the main analysis, we did not observe shrinkage of non-relevant or no-longer-relevant subspaces in correct trials when hyperalignment was omitted (Figure S17). However, in incorrect trials, we observed an expansion of non-relevant subspaces in the Inhibition condition at uncorrected level (Figure S18). Furthermore, the increase in subspace orthogonality following the cue in Control condition load 2 observed in the main analysis was also present when hyperalignment was omitted; however, this effect was captured only by VAF and not by PA (Figure S19). When considering shape metrics, Procrustes distances did not show any significant differences between conditions when hyperalignment was omitted. However, uncorrected analyses revealed a pattern qualitatively similar to the main results (Figure S20). In summary, hyperalignment across subjects enhanced our sensitivity to subtle dynamical changes in subspace configuration that were already present, but less detectable, without hyperalignment.

Second, we repeated the analysis while varying the number of principal components used to subset the Z matrix of PC scores before hyperalignment (p=[3, 5, 7, 15, 20, 100, 200, 400]), instead of the 10 PCs used in the main analysis. The number of PCs specifies the feature space used for the hyperalignment procedure. We observed that the main effects on separability remained significant with a hyperalignment dimensionality of p = 5 (Figure S21). For higher dimensionalities (p = [7, 15, 100, 200, 400]), the results were identical to those obtained with p=10 in the main analysis (Figure S22). However, with a more strongly reduced dimensionality (p=3), the results were indistinguishable from those obtained without hyperalignment (Figure S23). Inspection of the transformation matrices (p x p) from hyperalignment revealed that, at this low dimensionality, they were nearly identity matrices, leaving the PC scores almost unchanged after hyperalignment.

Third, we repeated the analysis using an alternative hyperalignment procedure. Rather than aligning subspaces to a common group template, as in (Heusser et al. 2017), we iteratively aligned all subjects to a single reference subject, repeated this procedure using each subject as the reference, and averaged the results across iterations. Under this approach, the main effects of separability remained significant; however, the expansion of subspaces in the Control condition was no longer observed (Figure S24).

Fourth, we examined whether our findings were robust to the dimensionality of the neural subspaces. Subspaces were defined using k=2,4,5,6,8,10 PCs. The main effects on separability were preserved for low-dimensional subspaces (k=2,4,5). Specifically, subspace expansion in the Control condition and expansion of relevant subspaces in the Inhibition condition were observed for k=2,4,5, whereas shrinkage of non-relevant subspaces in Inhibition was observed for k=4,5. In the Update condition, expansion of relevant subspaces occurred for k=2,4, and shrinkage of no-longer-relevant subspaces was present for k=2,4,5 (Figure S25–27). None of these effects were observed for subspaces with 6 or more PCs, indicating that the effects are confined to low-dimensional subspaces and do not extend across the full data dimensionality (Figure S28).

Fifth, we evaluated whether our findings were robust to the dimensionality of the brain parcellation. The main effects were replicated when using 200 and 300 parcels (Figures S29-30).

Sixth, we evaluated the effect of fidelity weighting, a custom preprocessing step applied during source modelling in which source time series are weighted according to their ability to recover phase information in simulated data (see Methods). When fidelity weighting was omitted while the remainder of the geometric analysis pipeline was kept unchanged, the main findings—namely, expansion of relevant subspaces and shrinkage of non-relevant subspaces following the cue—were preserved, although the temporal dynamics of these effects changed (Figure S31). In the inhibition condition, the expansion of relevant subspaces emerged later during the maintenance delay period, whereas the shrinkage of non-relevant subspaces remained largely unchanged (Figure S31B). In the update condition, the expansion of relevant subspaces was shorter in duration and reduced in magnitude, while the shrinkage of no-longer-relevant subspaces was preserved, as were the later expansions of non-relevant and no-longer-relevant subspaces (Figure S31C). When source modelling was instead performed using the standard MNE-Python pipeline, the results obtained with MNE (Figure S32) and dSPM (Figure S33) were broadly similar to those obtained with our custom pipeline without fidelity weighting. Overall, these findings suggest that downweighting source time series with poor phase-recovery fidelity enhances the sensitivity to capture task modulations of neural subspaces.

**SUPPLEMENTARY FIGURES**

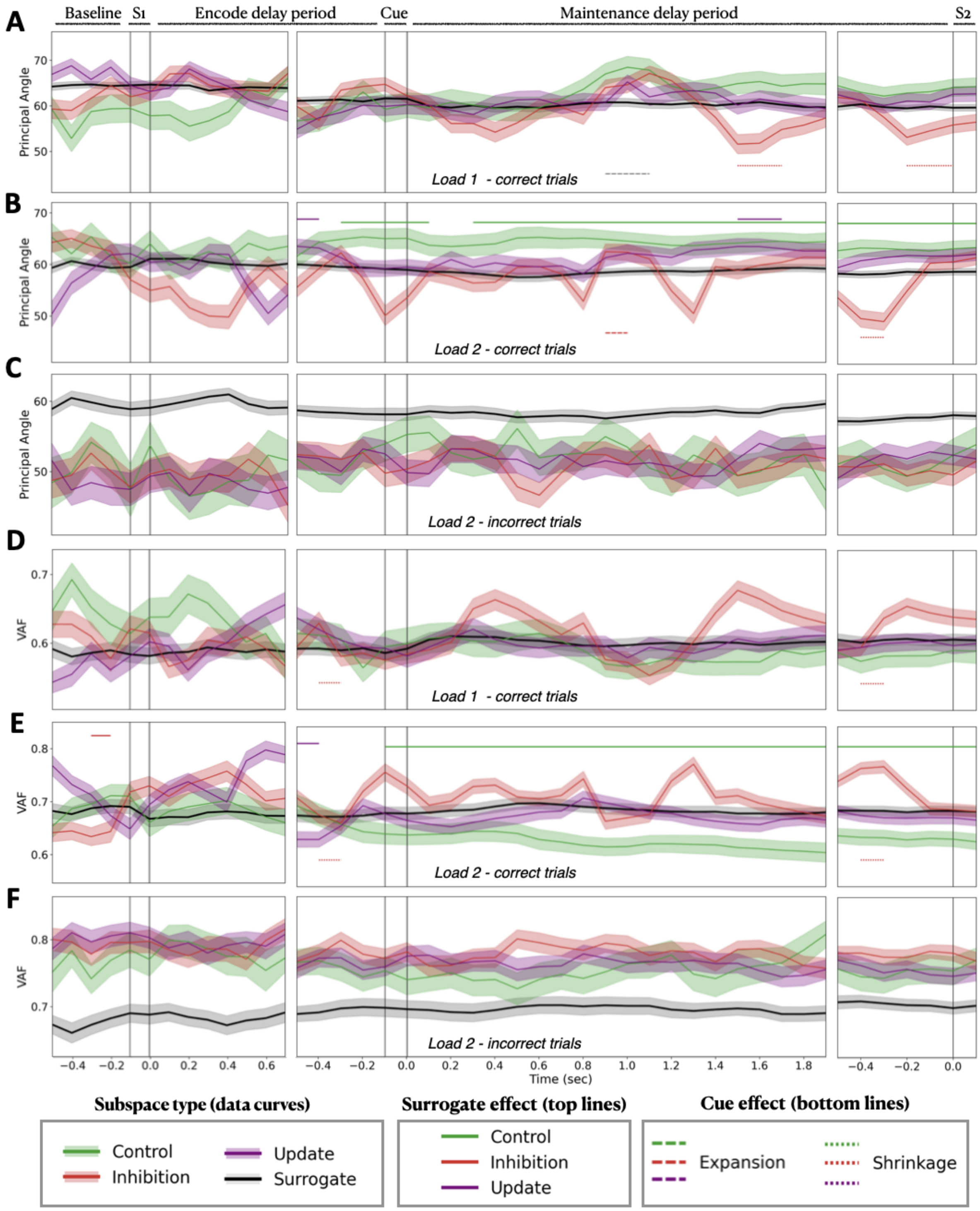


**Figure S1.** Subspace alignment quantified by Principal Angle (PA; A–C) and Variance Accounted For (VAF; D–F). (A, D) Correct trials of load 1; (B, E) correct trials of load 2; (C, F) incorrect trials of load 2. Panels correspond to different jitter-corrected delay durations (encode: 0.7–1 s; maintenance: 1.5–2 s). Solid vertical lines mark, from left to right, the onsets of S1, encode period, cue, maintenance period, and S2. The x-axis shows time in seconds, with each point representing the onset of a 0.4-s time window advanced in 0.1-s steps (e.g., x = 0 s corresponds to the interval [0, 0.4] s). Colored curves indicate the mean measure across subjects for the control, inhibition, and update conditions. Black lines indicate the corresponding measure computed from surrogate data, averaged across 1000 iterations; for visualization, the surrogate curve is shown only for the control condition. Shaded regions represent the standard error of the mean (SEM) across subjects. Horizontal bars at the top indicate time points showing a significant difference from surrogate data, whereas bars at the bottom indicate time points with a significant effect of cue (dependent-samples t-test, two-tailed, FDR-corrected, $p < 0.05$).

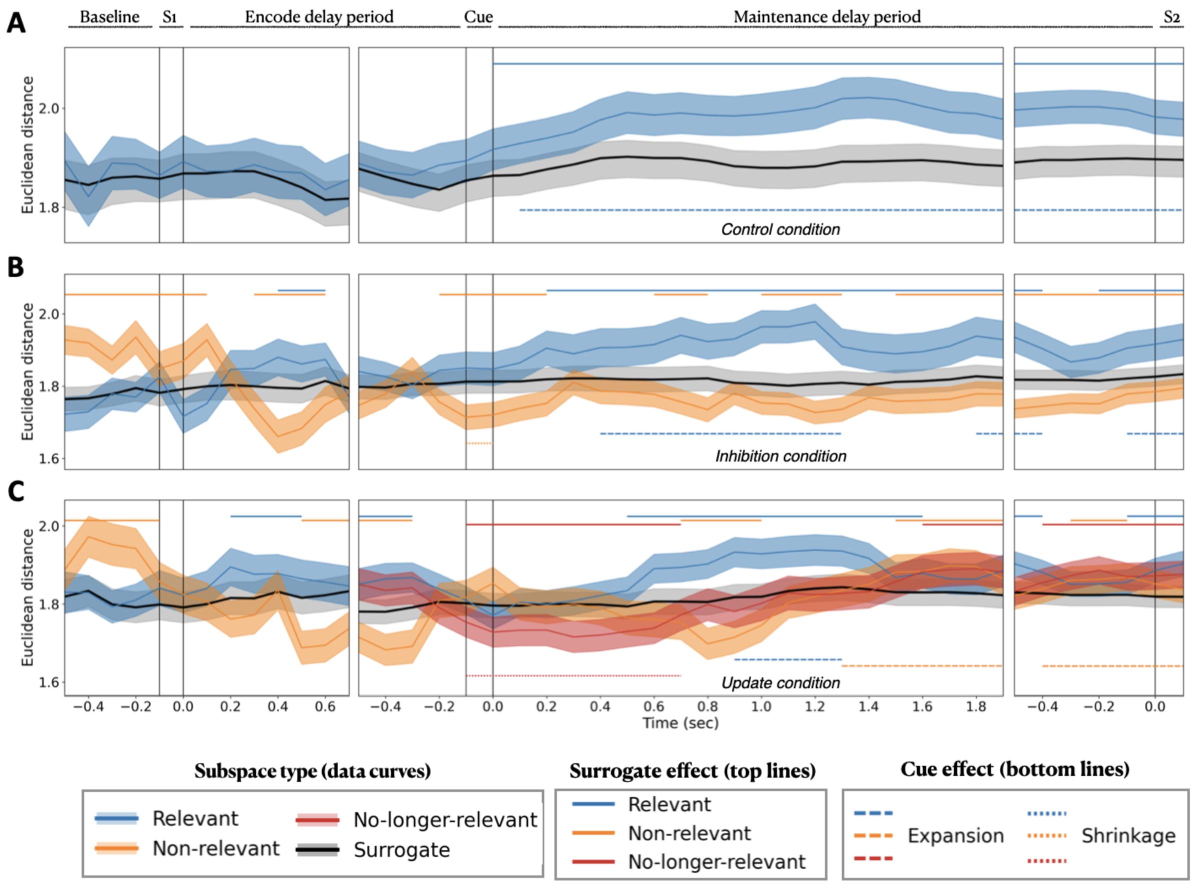


**Figure S2.** Separability of memory representations across the full trial. Separability within feature-specific neural subspaces, quantified as the Euclidean distance between rows of the Z* matrices of aligned PC scores (y-axis), combining stimulus loads 1 and 2. A–C) Time-resolved separability for A) control, B) inhibition, and C) update conditions. This figure shows the same data as Figure 2A-C, but displays the full trial time course, including all task events, whereas Figure 2 focuses on the maintenance period for visualization purposes. Panels correspond to different jitter-corrected delay durations (encode: 0.7–1 s; maintenance: 1.5–2 s). Solid vertical lines mark, from left to right, the onsets of S1, encode period, cue, maintenance period, and S2. The x-axis shows time in seconds, with each point representing the onset of a 0.4-s time window advanced in 0.1-s steps (e.g., x = 0 s corresponds to the interval [0, 0.4] s). Colored solid lines represent mean Euclidean distances across subjects, with blue denoting relevant features, orange non-relevant features, and red no-longer-relevant features. Dashed lines represent separability estimated from surrogate data, averaged across 1,000 surrogate iterations. Dashed lines represent separability estimated from surrogate data, averaged across 1,000 surrogate iterations. Shaded regions indicate the standard error of the mean (SEM) across subjects. Horizontal bars at the top indicate time points at which separability significantly differed from surrogate data, whereas bars at the bottom indicate time points showing a significant cue effect (dependent-samples, two-tailed t-test; FDR-corrected, $p<0.05$).

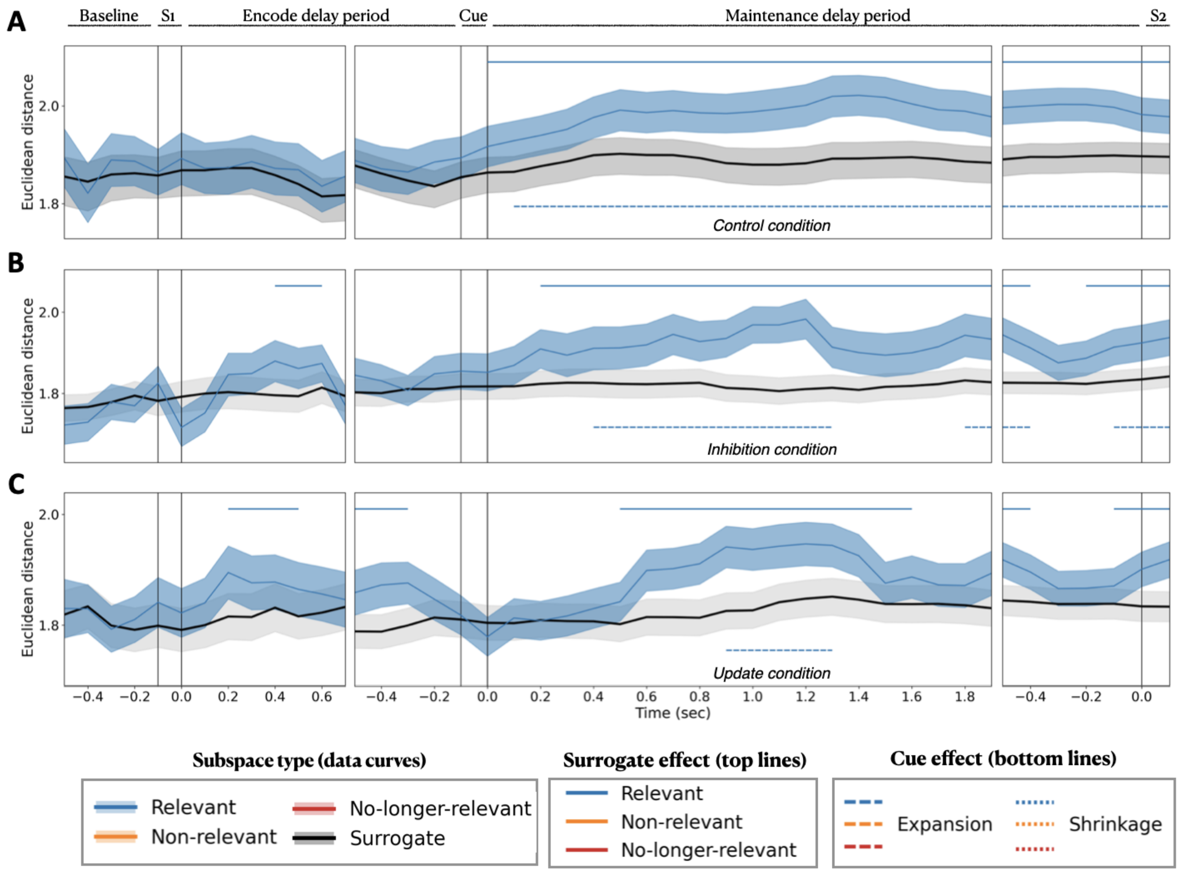


**Figure S3.** Separability of memory representations within the relevant-feature subspace. This figure shows the same data as Figure 2, displayed separately for the relevant-feature subspace. A) Relevant subspaces in Control condition, B) Relevant subspaces in Inhibition condition, and C) Relevant subspaces in Update condition. Panel layout, time axis, and significance markers follow the conventions described in Figure S2.

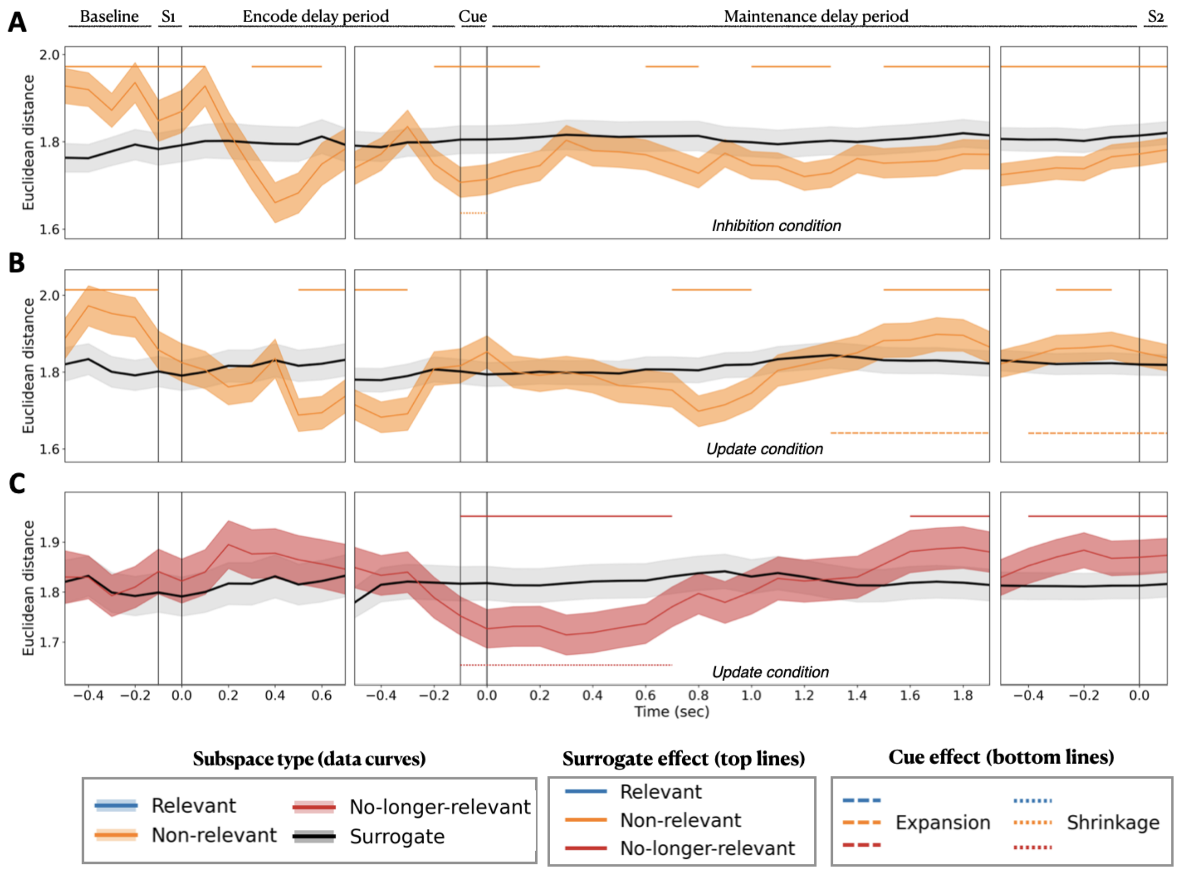


**Figure S4.** Separability of memory representations within the relevant-feature subspace. This figure shows the same data as Figure 2, displayed separately for the relevant-feature subspace. A) Non-relevant subspaces in Inhibition condition, B) Non-relevant subspaces in Update condition, and C) No-longer-relevant subspaces in Update condition. Panel layout, time axis, and significance markers follow the conventions described in Figure S2.

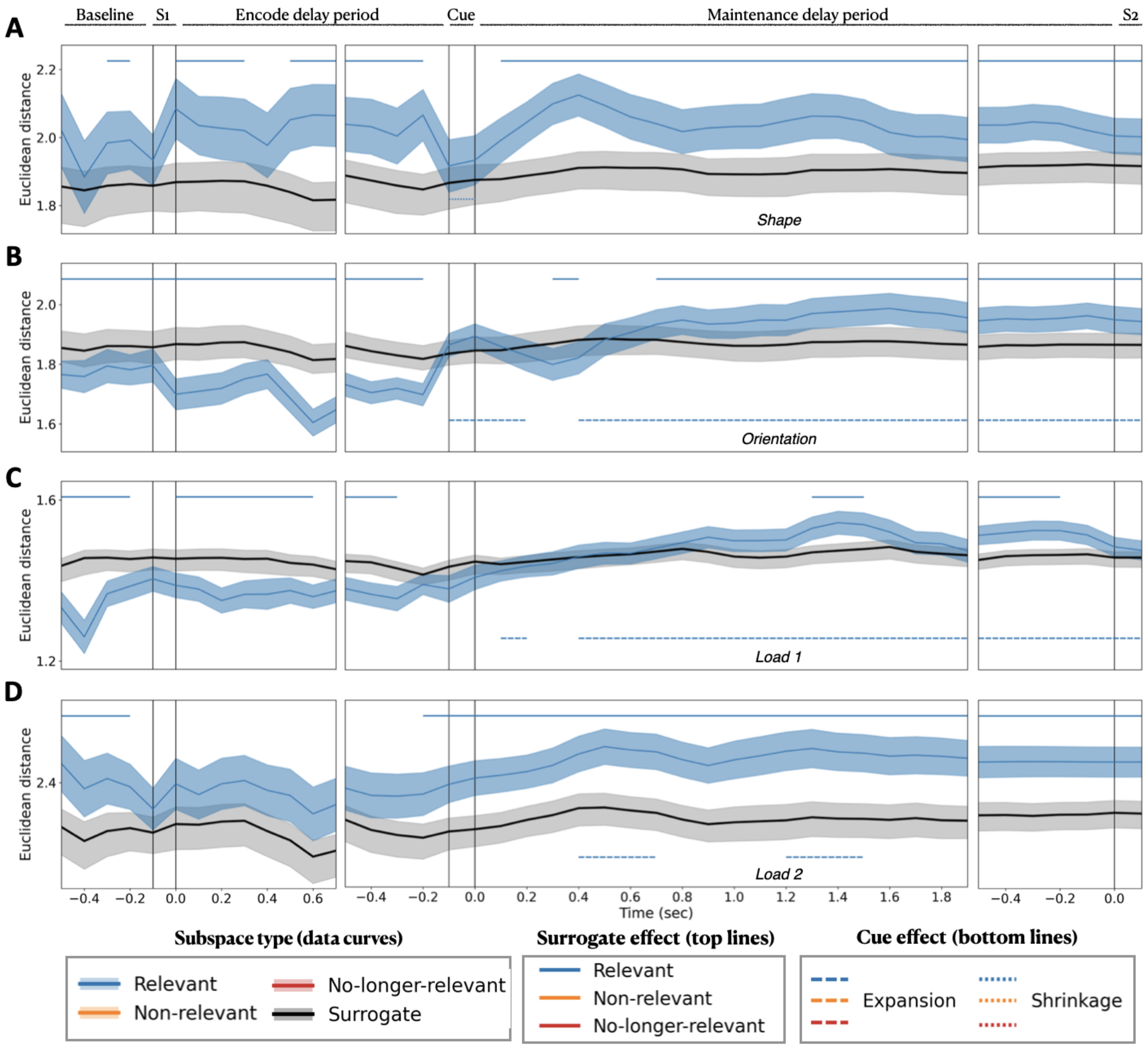


**Figure S5.** Separability of memory representations within feature-specific neural subspaces in the control condition, for A) Polygons (shape), B) Gratings (orientation), C) Stimulus load 1, and D) Stimulus load 2. Panel layout, time axis, and significance markers follow the conventions described in Figure S2.

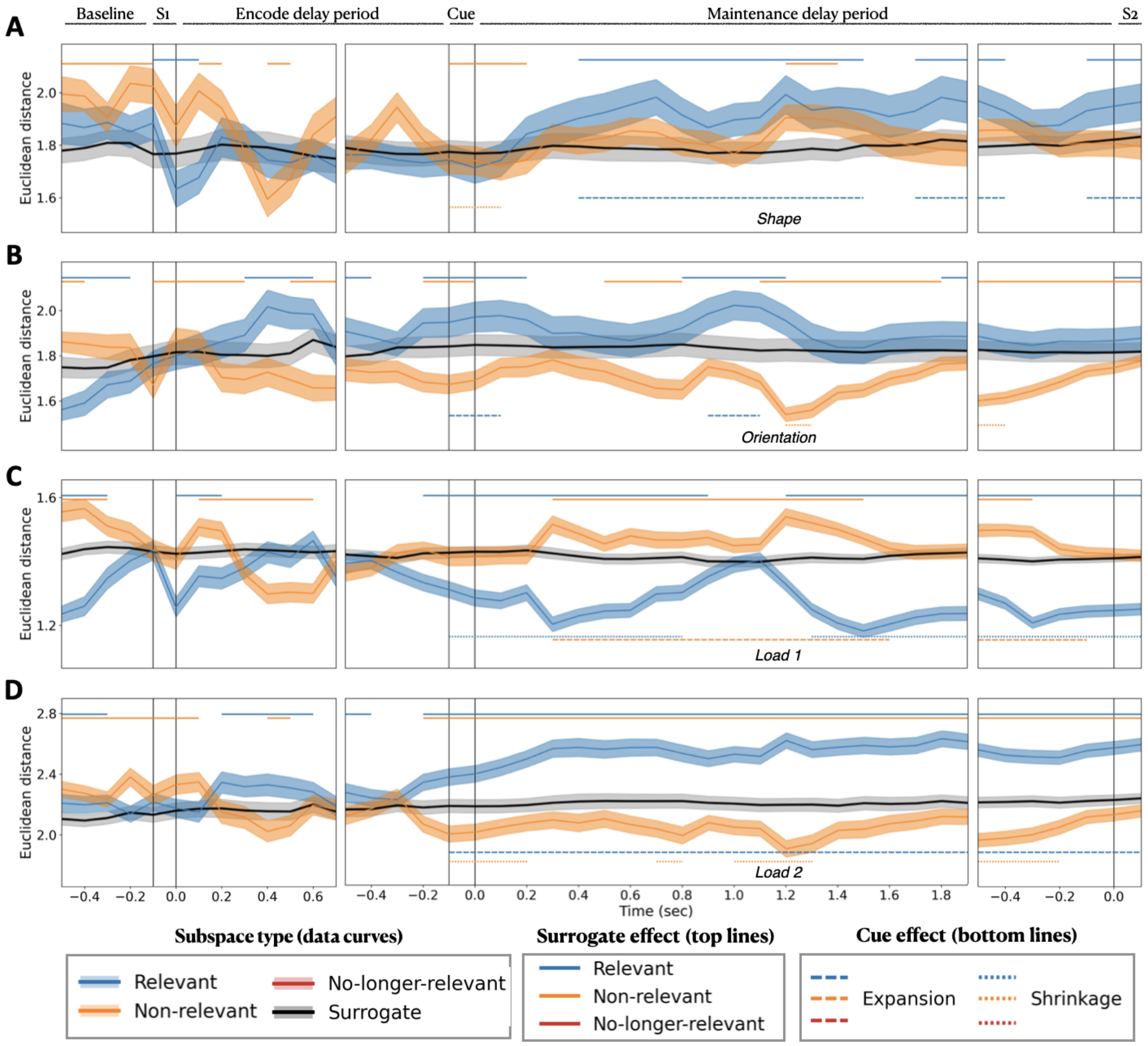


**Figure S6.** Separability of memory representations within feature-specific neural subspaces in the inhibition condition, for A) Polygons (shape), B) Gratings (orientation), C) Stimulus load 1, and D) Stimulus load 2. Panel layout, time axis, and significance markers follow the conventions described in Figure S2.

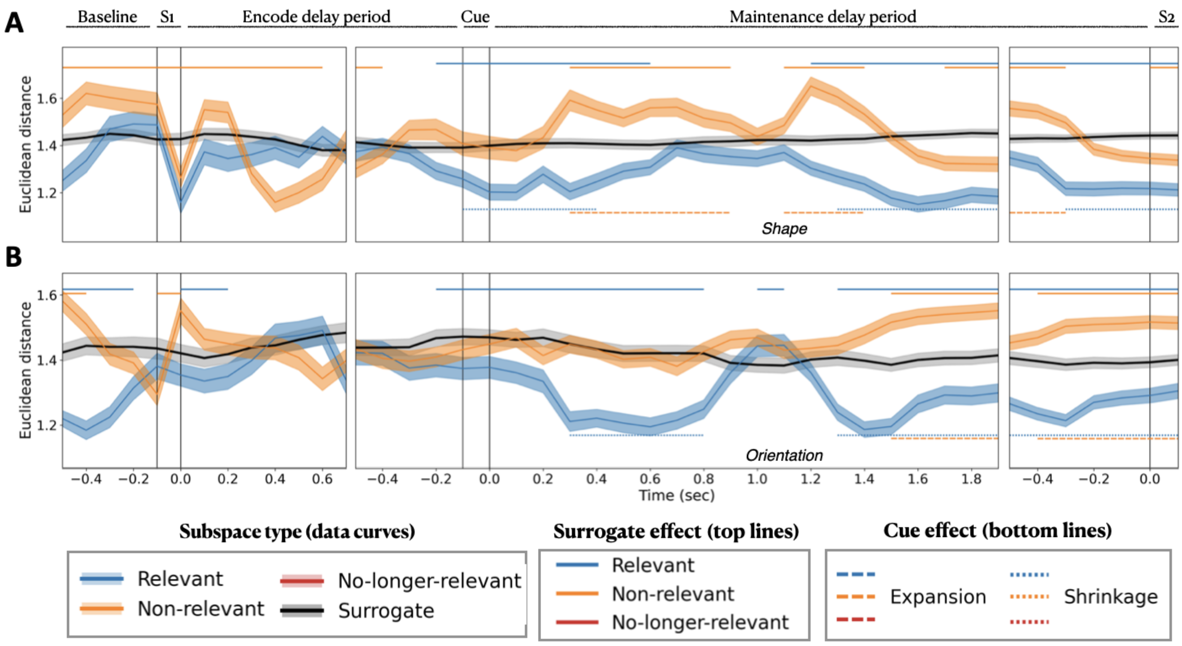


**Figure S7.** Separability of memory representations within feature-specific neural subspaces in the inhibition condition in stimulus load 1 for A) Polygons (shape), B) Gratings (orientation). Panel layout, time axis, and significance markers follow the conventions described in Figure S2.

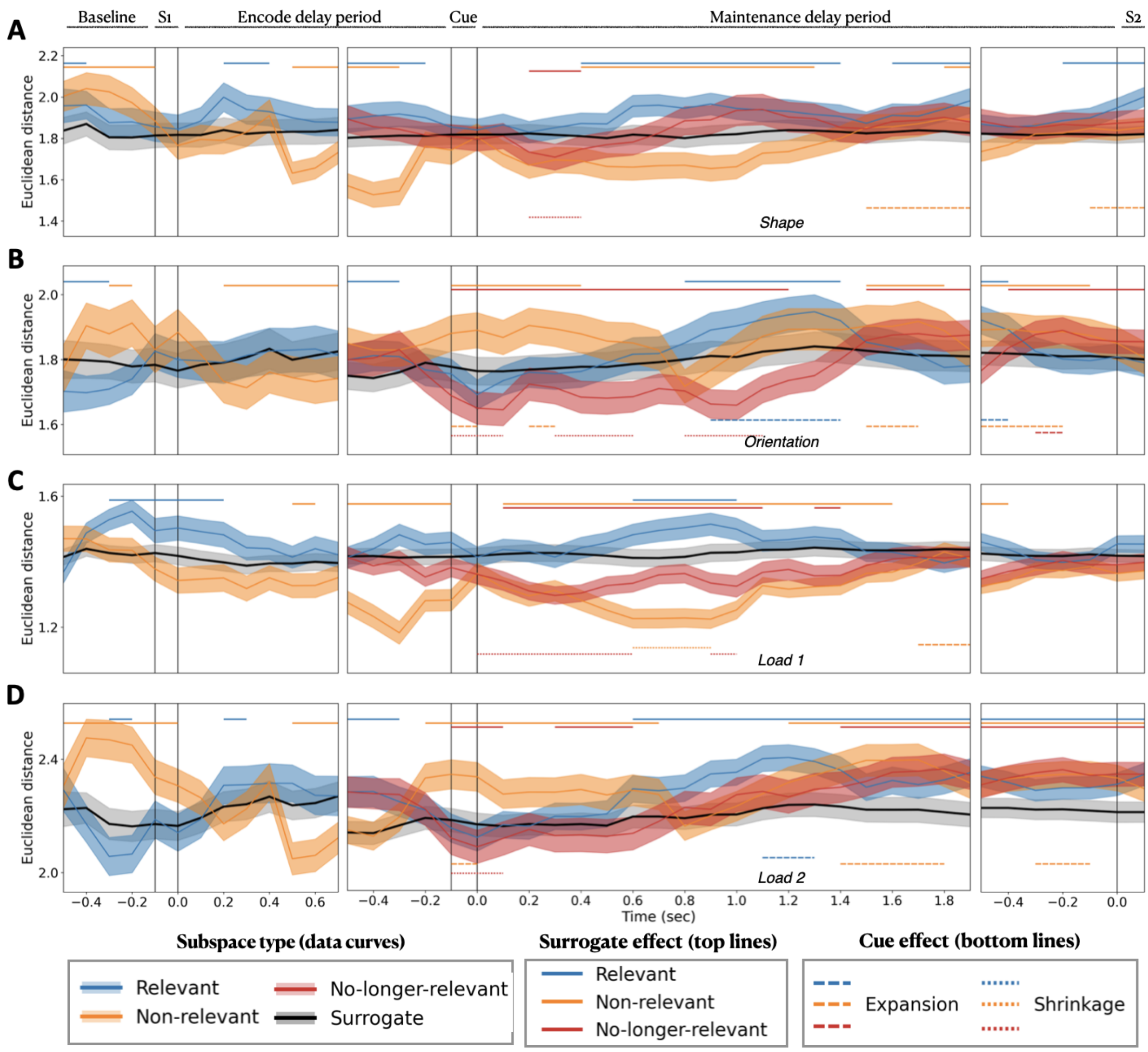


**Figure S8.** Separability of memory representations within feature-specific neural subspaces in the update condition, for A) Polygons (shape), B) Gratings (orientation), C) Stimulus load 1, and D) Stimulus load 2. Panel layout, time axis, and significance markers follow the conventions described in Figure S2.

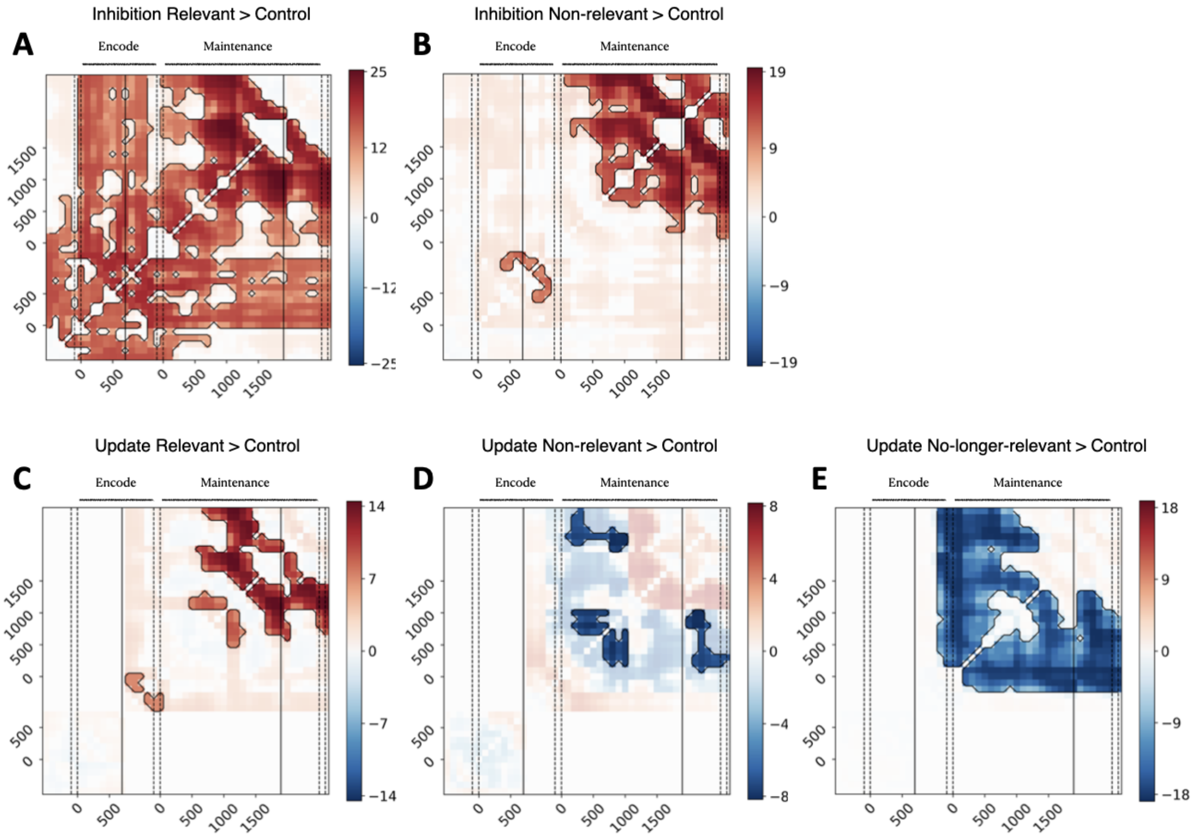


**Figure S9.** Time-generalization of Procrustes distances without scaling. Panels correspond to the contrasts: A) Inhibition Relevant > Control, B) Inhibition Non-relevant > Control, C) Update Relevant > Control, D) Update Non-relevant > Control, E) Update No-longer-relevant > Control. Panel layout, time axes, and markers follow the same conventions as in Figure 5.

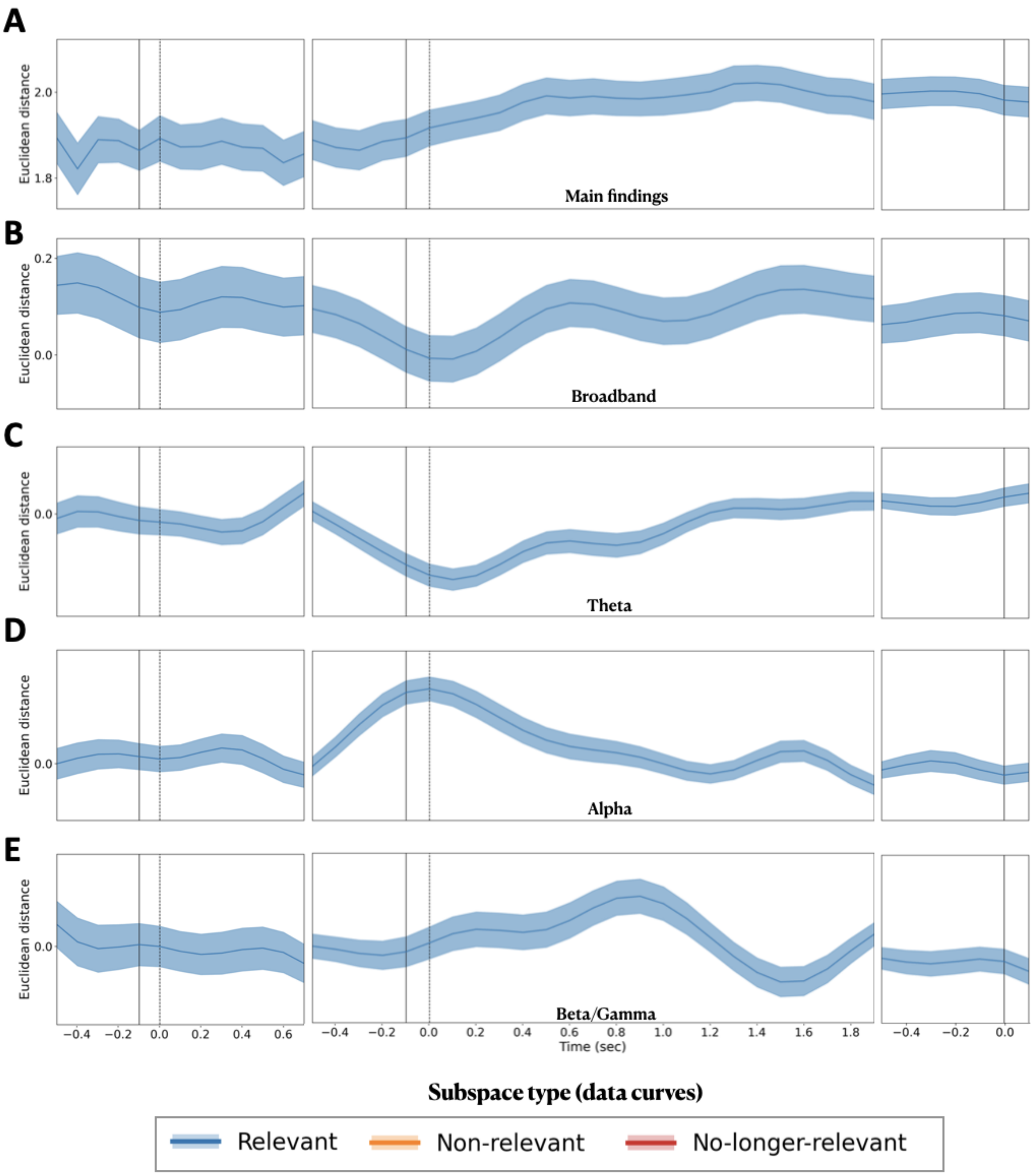


**Figure S10.** Separability of the control condition for correct trials. (A) Main findings on separability (same as Figure 2), corresponding to the response variable in the models shown in Figure 5B. (B) Broadband component, (C) theta-band component, (D) alpha-band component, and (E) beta/gamma-band component, corresponding to the predictor variables in the models shown in Figure 5B. Panel layout, time axis, and significance markers follow the conventions described in Figure S2.

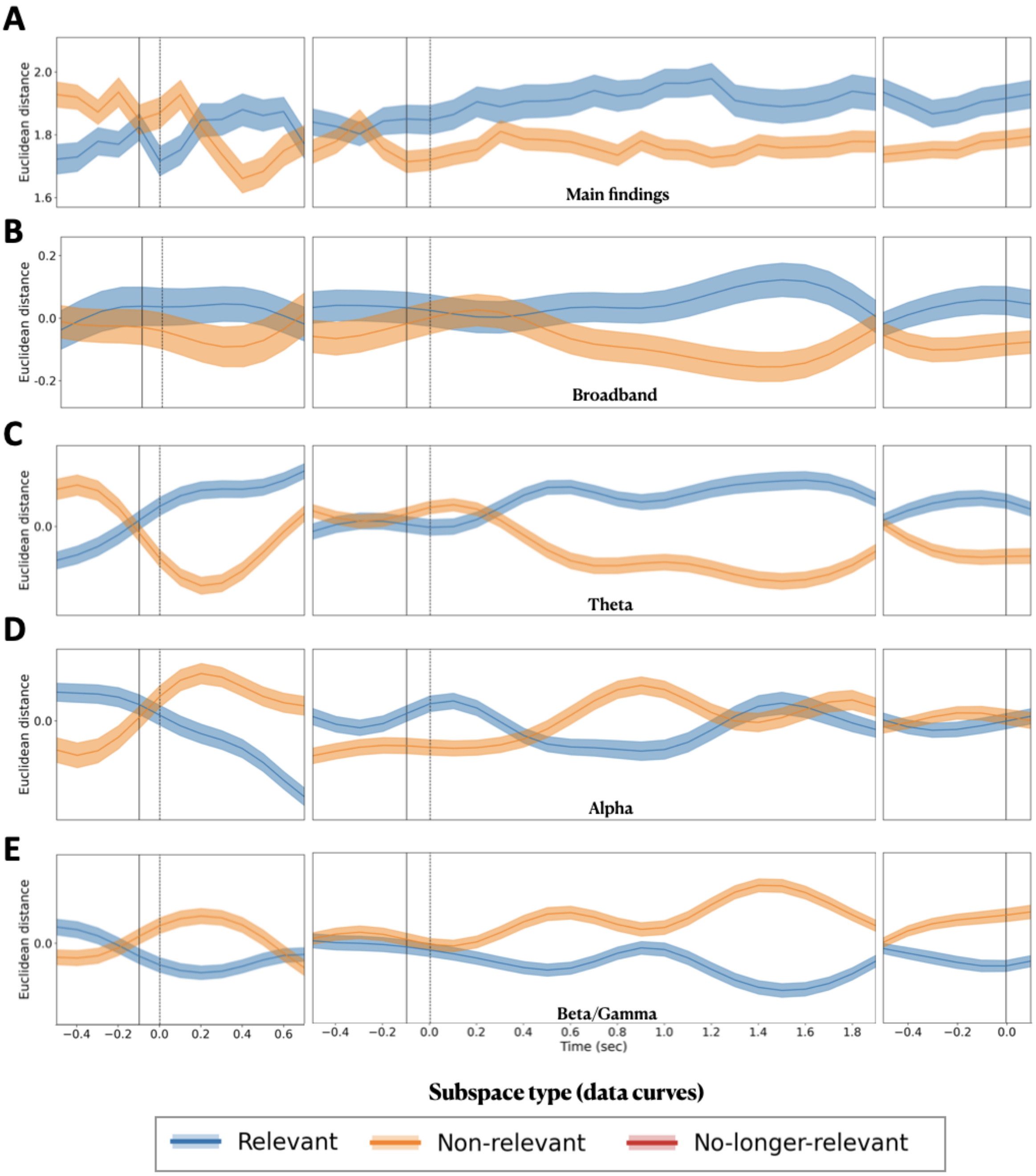


**Figure S11.** Separability of the inhibition condition for correct trials. (A) Main findings on separability (same as Figure 2), corresponding to the response variable in the models shown in Figure 5B. (B) Broadband component, (C) theta-band component, (D) alpha-band component, and (E) beta/gamma-band component, corresponding to the predictor variables in the models shown in Figure 5B. Panel layout, time axis, and significance markers follow the conventions described in Figure S2.

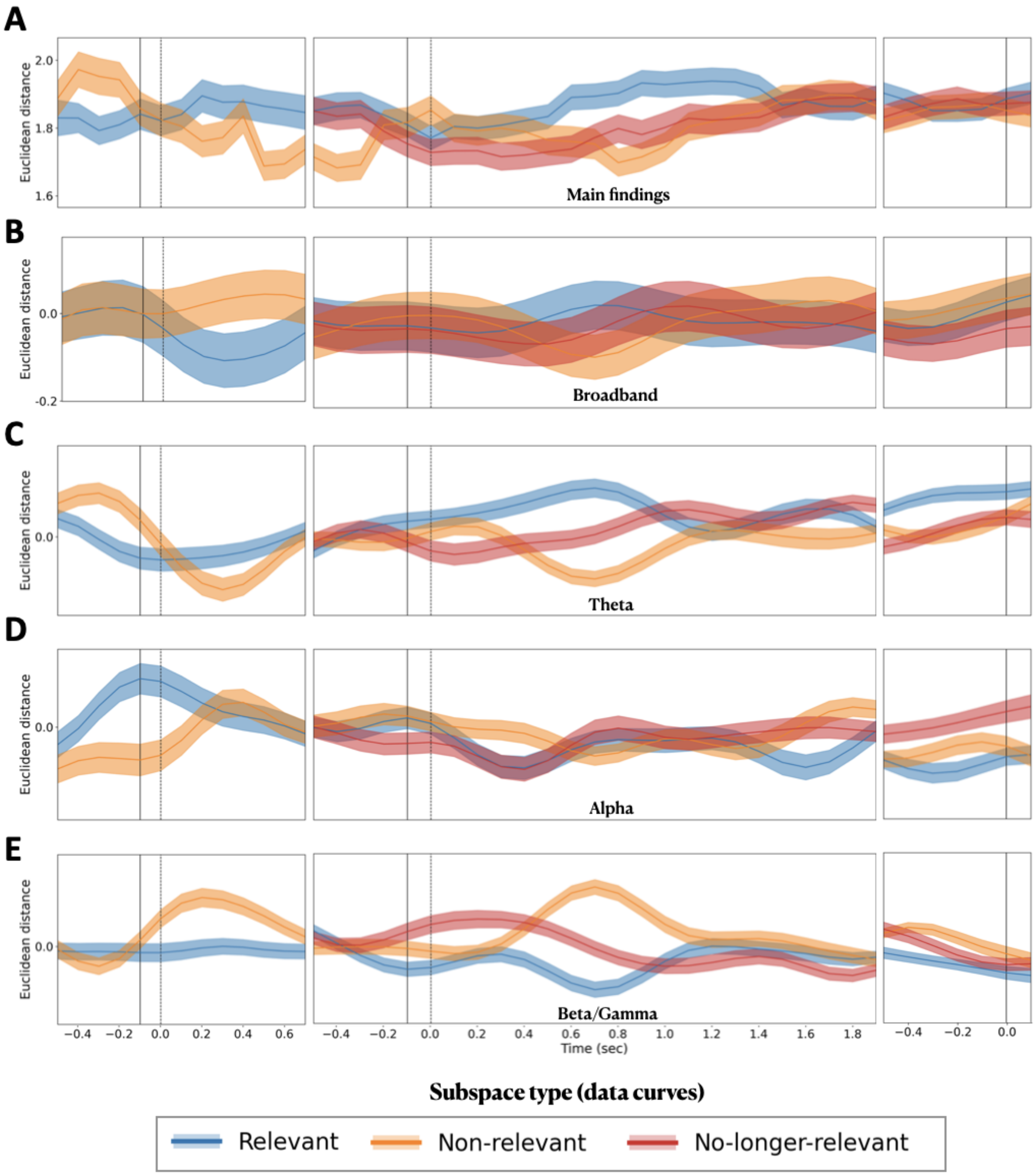


**Figure S12.** Separability of the update condition for correct trials. (A) Main findings on separability (same as Figure 2), corresponding to the response variable in the models shown in Figure 5B. (B) Broadband component, (C) theta-band component, (D) alpha-band component, and (E) beta/gamma-band component, corresponding to the predictor variables in the models shown in Figure 5B. Panel layout, time axis, and significance markers follow the conventions described in Figure S2.

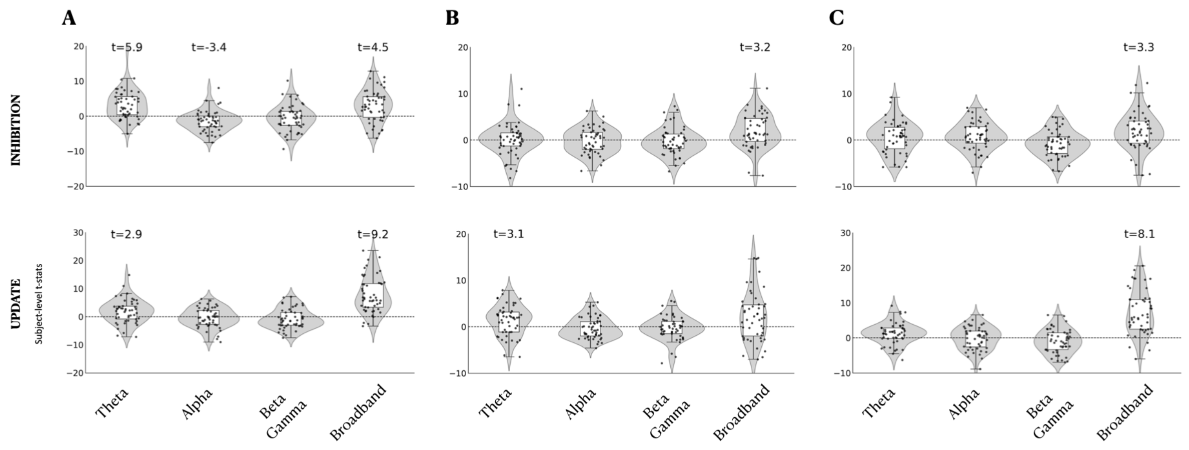


**Figure S13.** Hierarchical linear models predicting main findings on separability from the broadband component and residualized narrowband (NB) oscillatory components are shown. Panel A shows results including both relevant and non-relevant subspaces (also shown in Figure 5 for comparison), panel B shows only relevant subspaces, and panel C shows only non-relevant subspaces. Results were derived from the real part of complex Morlet wavelets, with frequency bands defined using clustering-derived group-level frequency boundaries. Predictor variables are on the x-axis; first-level t-statistics are on the y-axis. Second-level t-statistics reported in the text correspond to regressors that remained significant after FDR correction ($p < 0.05$, two-tailed; † indicates a trend toward significance).

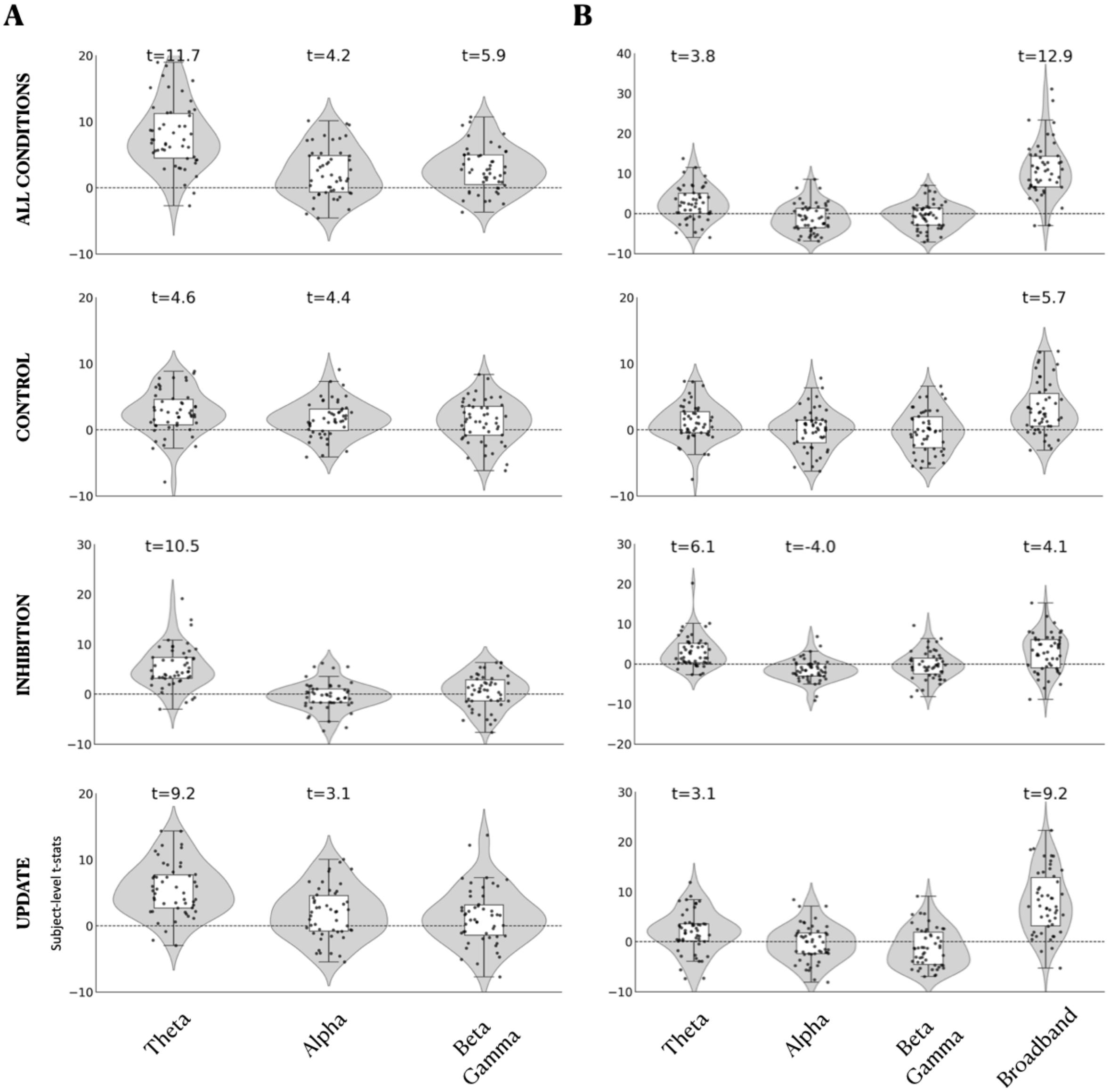


**Figure S14.** Oscillatory vs broadband geometry. Results from hierarchical linear models (A) predicting the main findings on separability from NB oscillatory components (model 1), and (B) predicting the main findings on separability from broadband component and residualized NB oscillatory components (model 2). Results derived from the real part of the complex Morlet wavelets, and frequency bands defined based on clustering-derived frequency boundaries at the group level. Predictor variables are shown on the x-axis. First-level t-statistics are shown on the y-axis, and second-level t-statistics are reported in the text for regressors that remained significant after FDR correction ($p < 0.05$, two-tailed).

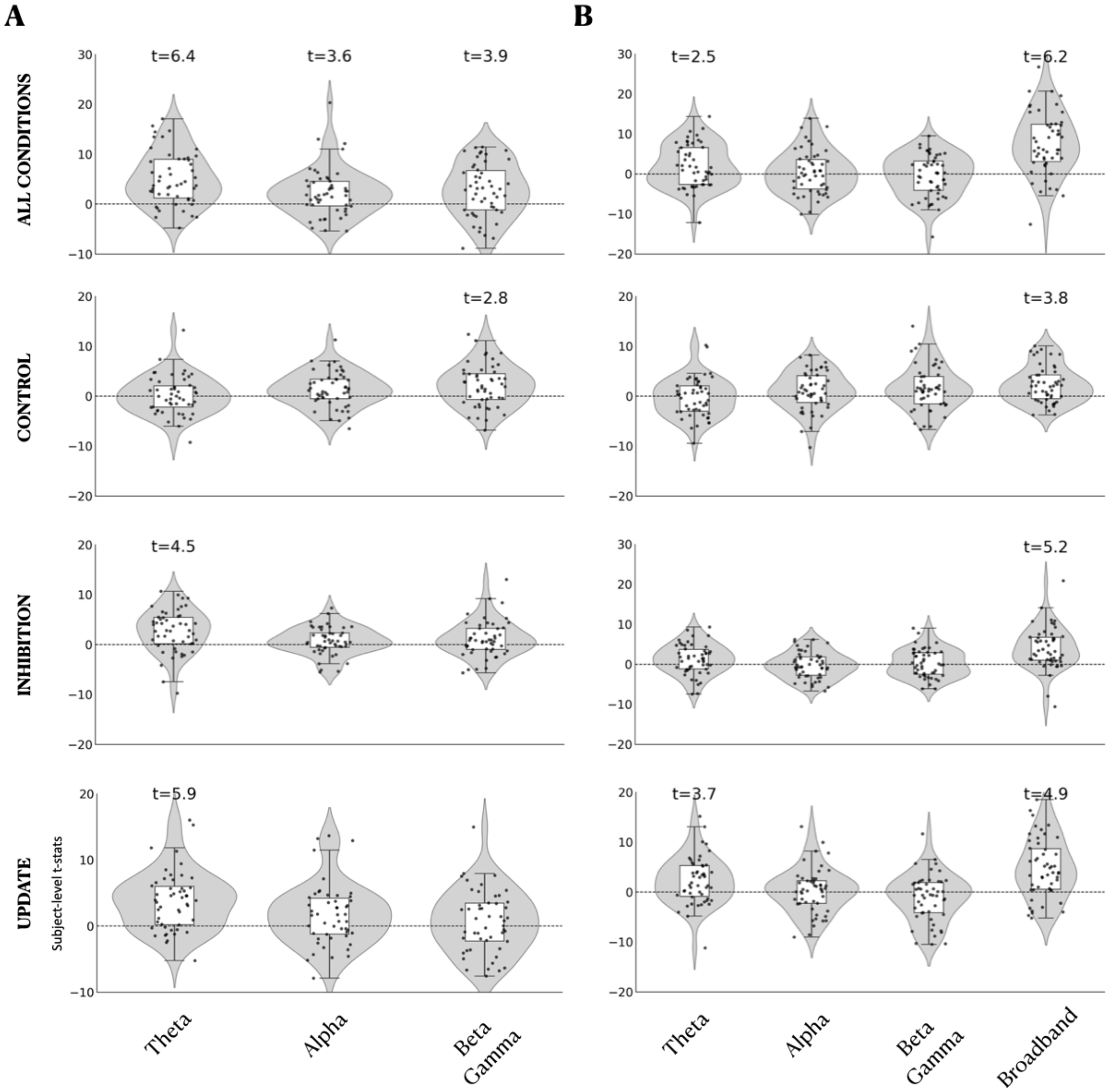


**Figure S15.** Oscillatory vs broadband geometry. Results from hierarchical linear models (A) predicting the main findings on separability from NB oscillatory components (model 1), and (B) predicting the main findings on separability from broadband component and residualized NB oscillatory components (model 2). Results derived from the amplitude envelopes of the complex Morlet wavelets, and frequency bands defined based on clustering-derived frequency boundaries at the subject level. Predictor variables are shown on the x-axis. First-level t-statistics are shown on the y-axis, and second-level t-statistics are reported in the text for regressors that remained significant after FDR correction ($p < 0.05$, two-tailed).

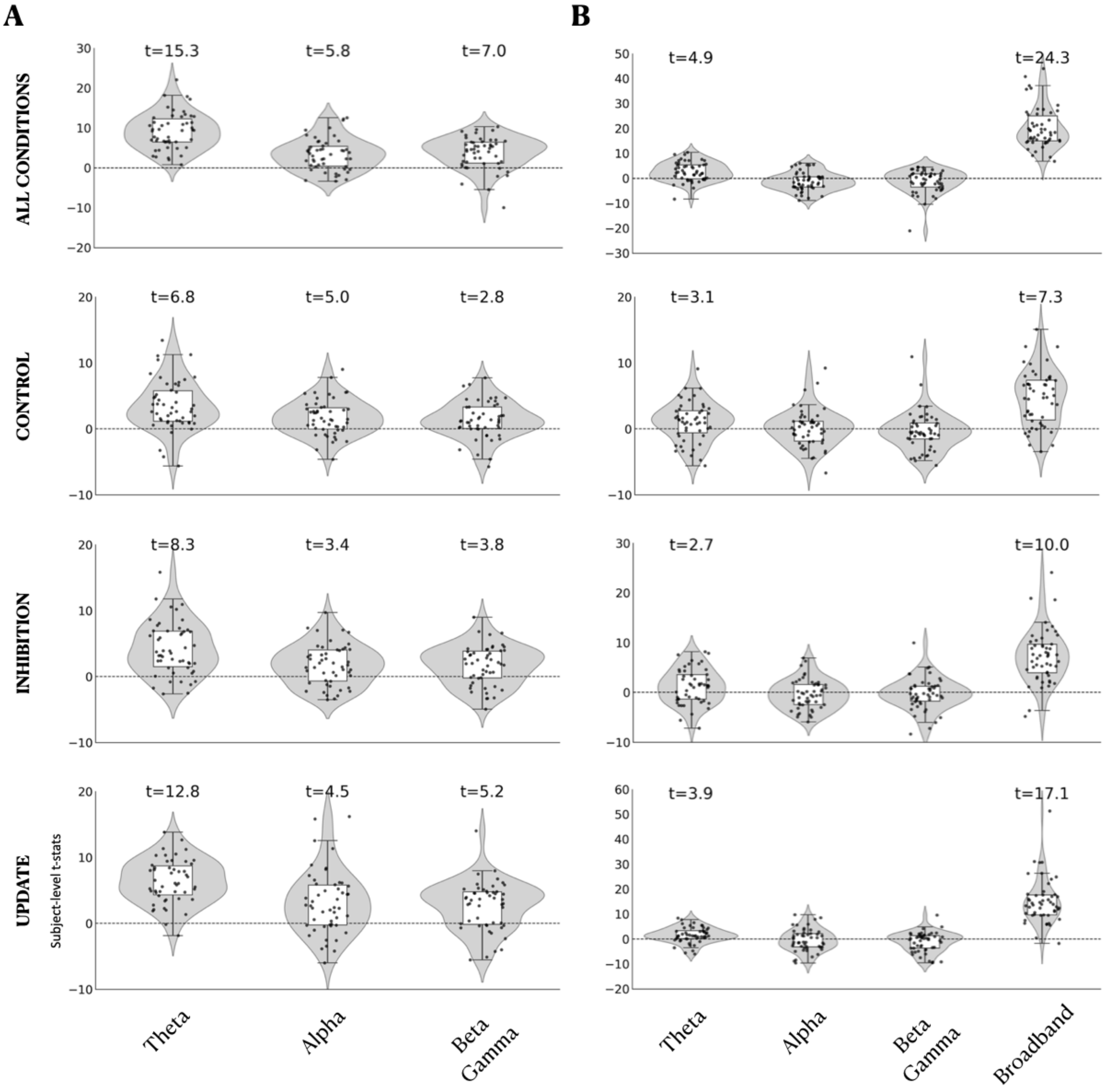


**Figure S16.** Oscillatory vs broadband geometry. Results from hierarchical linear models (A) predicting the main findings on separability from NB oscillatory components (model 1), and (B) predicting the main findings on separability from broadband component and residualized NB oscillatory components (model 2). Results derived from the real part of the complex Morlet wavelets, frequency bands defined based on clustering-derived frequency boundaries at the subject level, and hyperalignment transformation derived from broadband data. Predictor variables are shown on the x-axis. First-level t-statistics are shown on the y-axis, and second-level t-statistics are reported in the text for regressors that remained significant after FDR correction ($p < 0.05$, two-tailed).

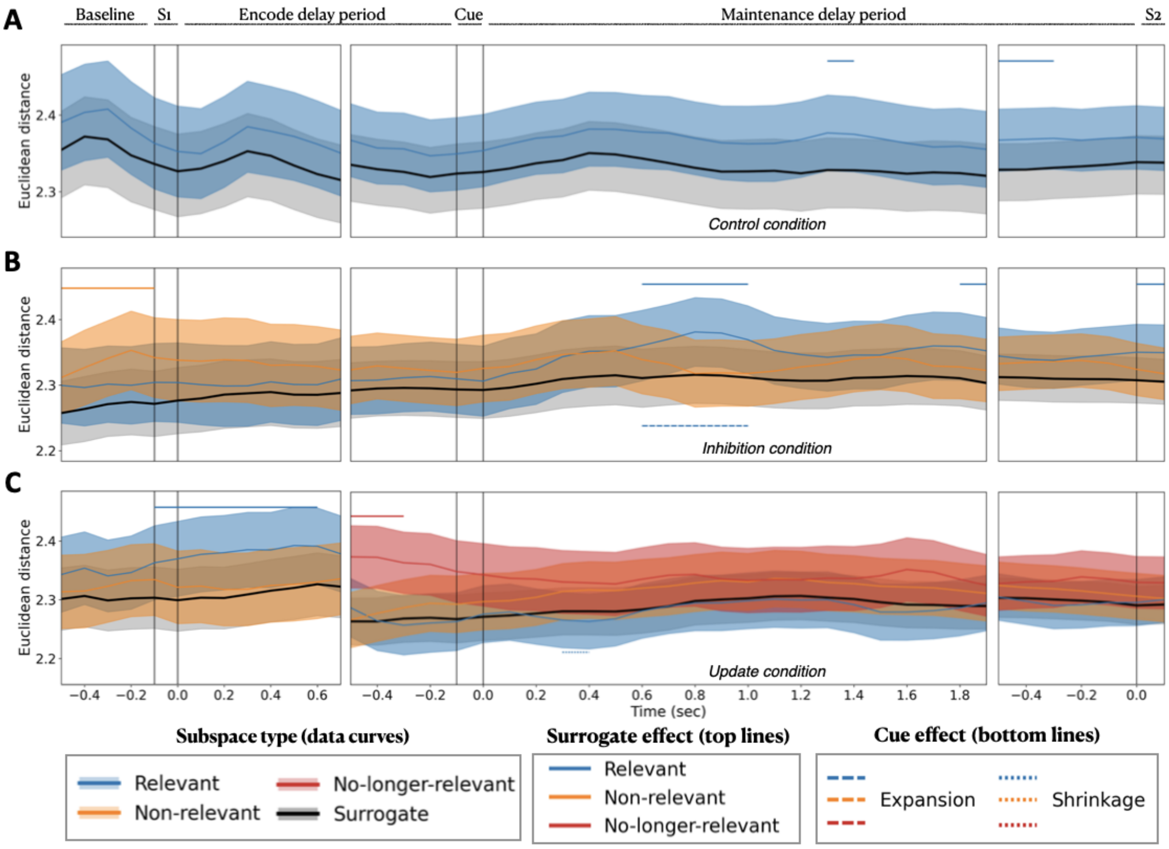


**Figure S17.** Supplementary geometric analysis performed without hyperalignment. Separability of memory representations within feature-specific neural subspaces for correct trials, shown for A) Control condition, B) Inhibition condition, and B) Update condition. Significant timepoints of surrogate effect (top lines) and cue effect (bottom lines) are uncorrected for multiple comparisons. No significant timepoints remained after FDR-correction. Panel layout, time axis, and significance markers follow the conventions described in Figure S2.

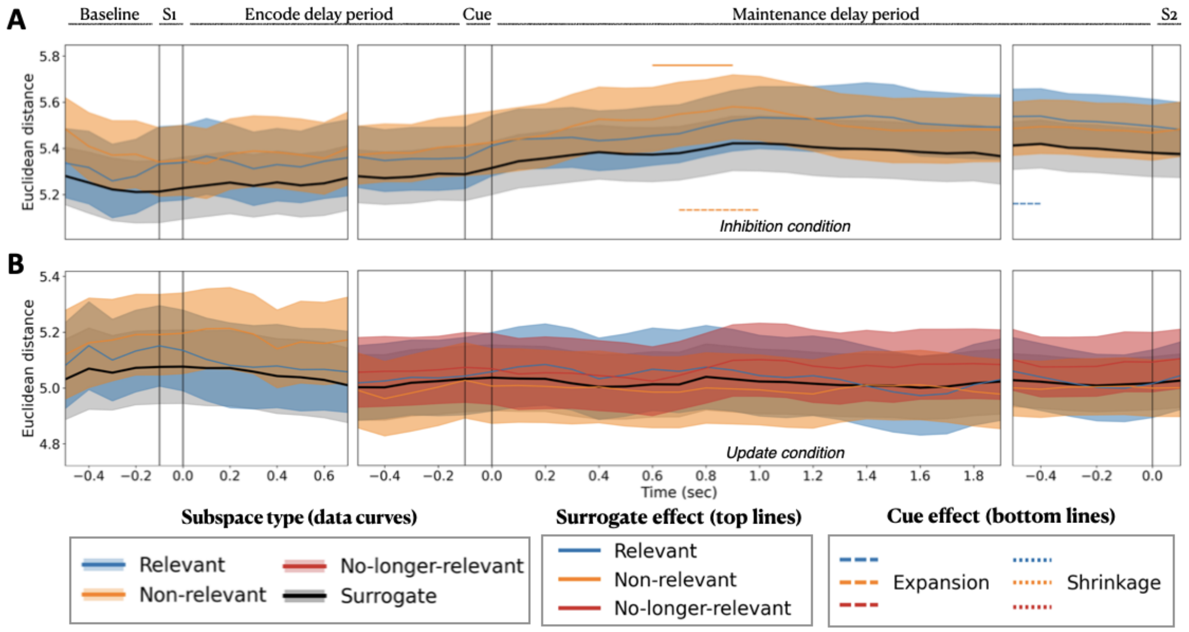


**Figure S18.** Supplementary geometric analysis performed without hyperalignment. Separability of memory representations within feature-specific neural subspaces for incorrect trials at stimulus load 2, shown for (A) Inhibition condition and (B) Update condition. Significant timepoints of surrogate effect (top lines) and cue effect (bottom lines) are uncorrected for multiple comparisons. No significant timepoints remained after FDR-correction. Panel layout, time axis, and significance markers follow the conventions described in Figure S2.

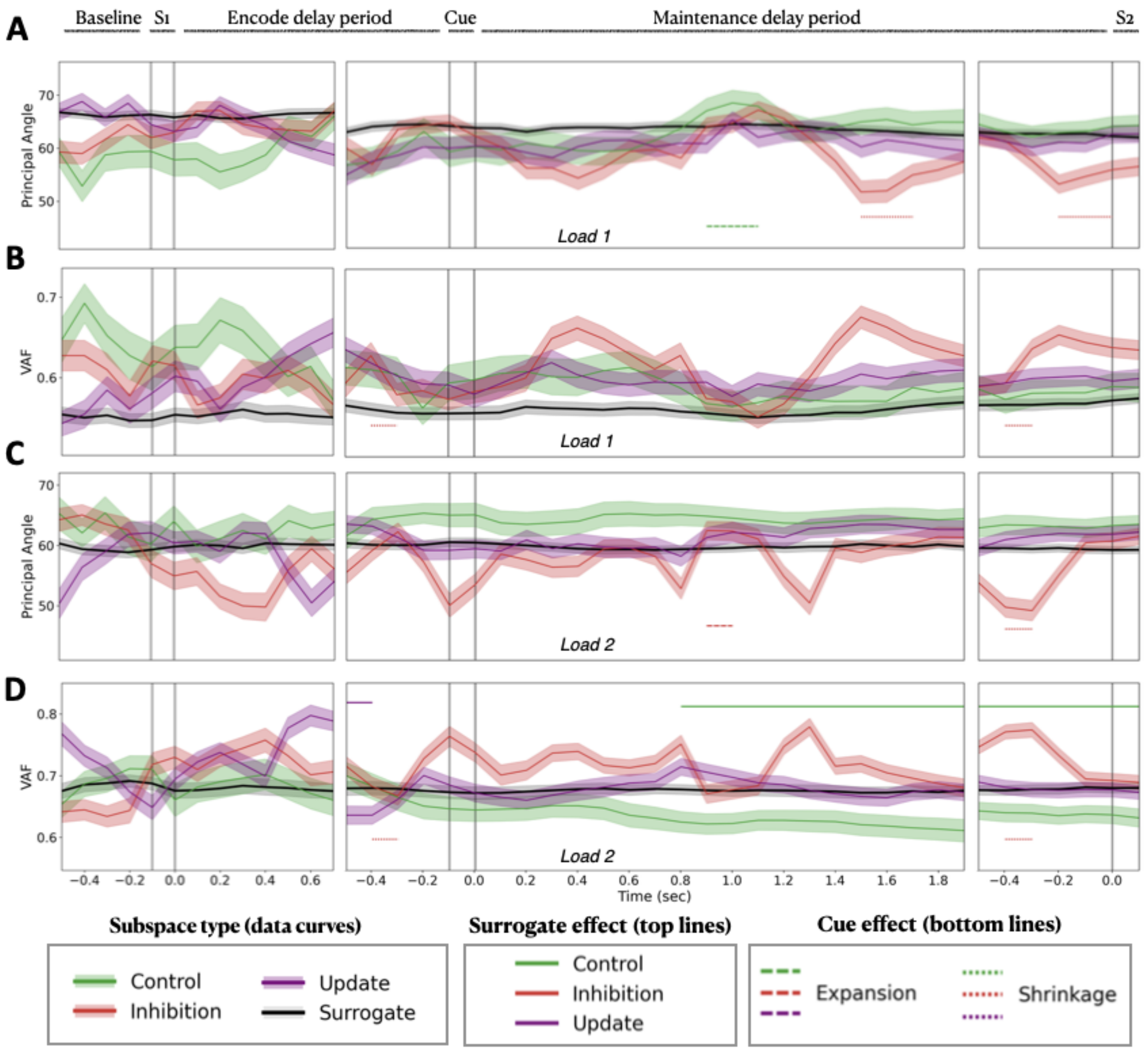


**Figure S19.** Supplementary analysis performed without hyperalignment for correct trials. Results are shown for (A) PA, stimulus 1; (B) VAF, stimulus 1; (C) PA, stimulus 2; (D) VAF, stimulus 2. Panel layouts, time axes, and markers follow the same conventions as in Figure S1.

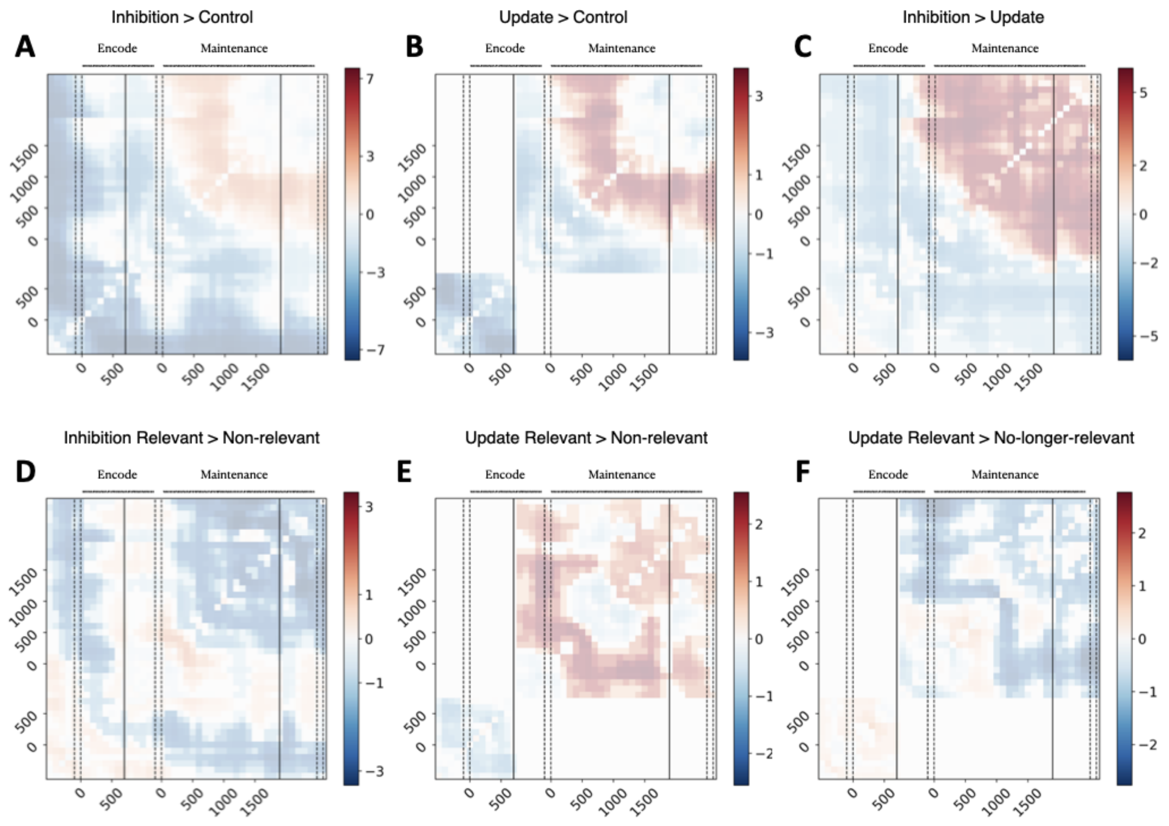


**Figure S20.** Supplementary analysis performed without hyperalignment for correct trials. Time-generalization of Procrustes distances without scaling. Panels correspond to the contrasts: A) Inhibition > Control, B) Update > Control, C) Inhibition > Update, D) Inhibition Relevant > Non-relevant, E) Update Relevant > Non-relevant, and F) Update Relevant > No-longer-relevant. Panel layouts, time axes, and markers follow the same conventions as in Figure 5.

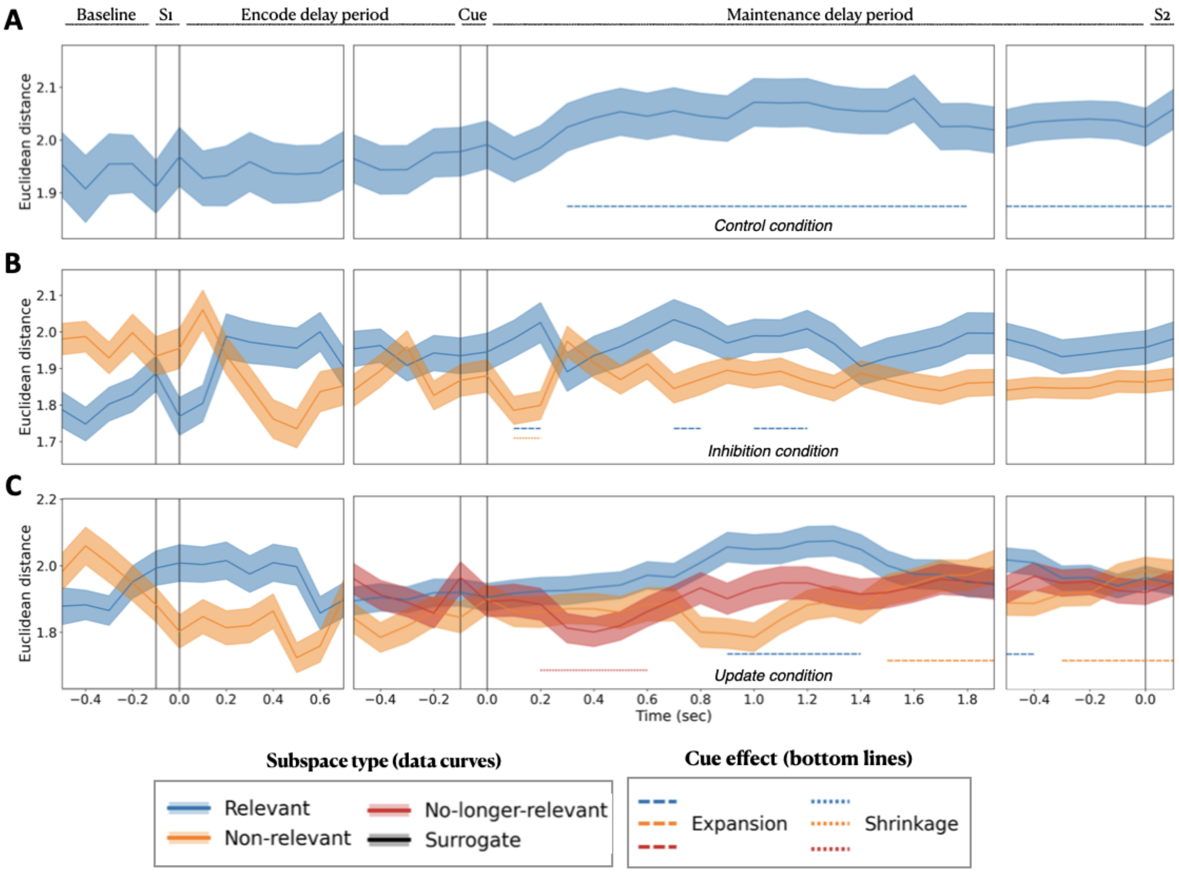


**Figure S21.** Supplementary analysis of neural geometry with hyperalignment applied on the 5 PCs of Z matrices of PC scores, instead of the 10 PCs used in the main analysis. Separability of memory representations within feature-specific neural subspaces for correct trials, shown for A) Control condition, B) Inhibition condition, and B) Update condition. Significant timepoints of cue effect (bottom lines) are FDR-corrected for multiple comparisons. Panel layout, time axis, and significance markers follow the conventions described in Figure S2.

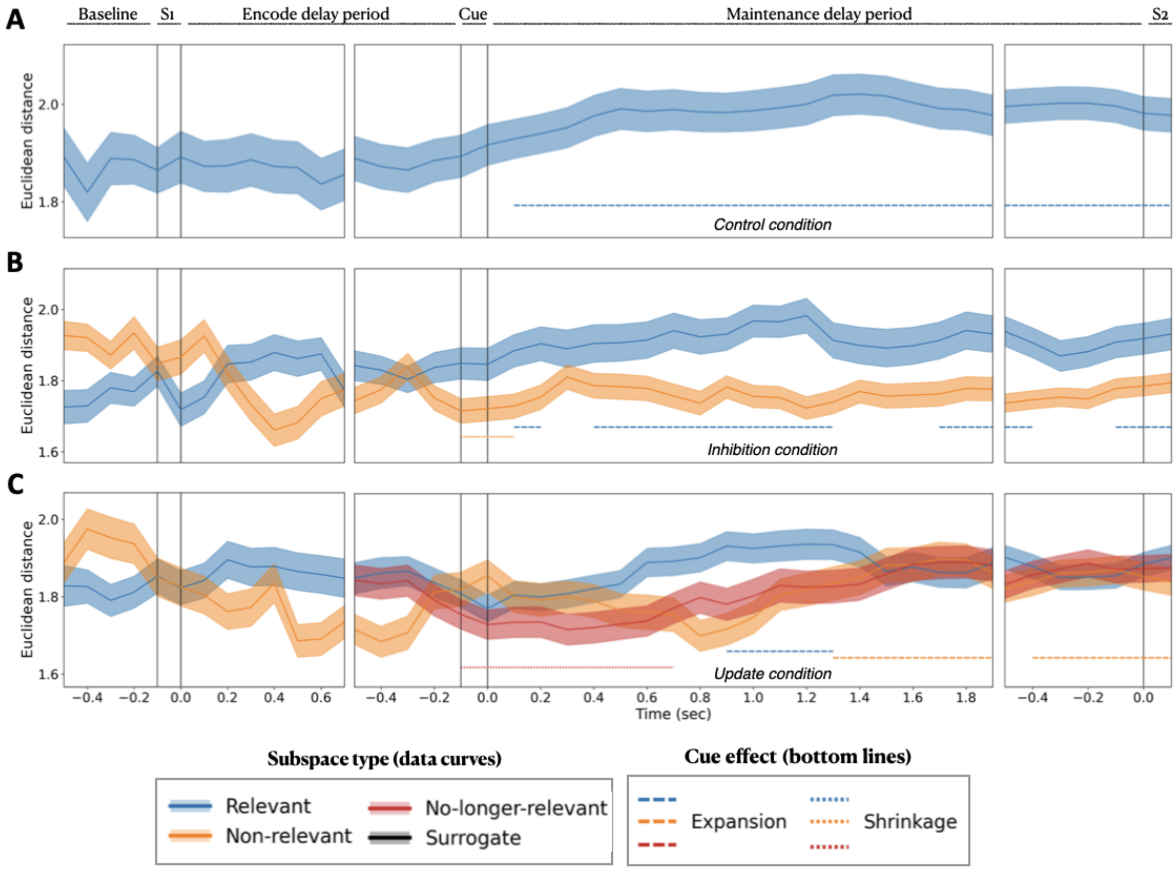


**Figure S22.** Supplementary analysis of neural geometry with hyperalignment applied on the 7 PCs of Z matrices of PC scores, instead of the 10 PCs used in the main analysis. Results are identical to those of the main analysis (Figure 2) and remain unchanged for higher hyperalignment dimensionalities (p = [15, 20, 100, 400]). Separability of memory representations within feature-specific neural subspaces for correct trials, shown for A) Control condition, B) Inhibition condition, and B) Update condition. Significant timepoints of cue effect (bottom lines) are FDR-corrected for multiple comparisons. Panel layout, time axis, and significance markers follow the conventions described in Figure S2.

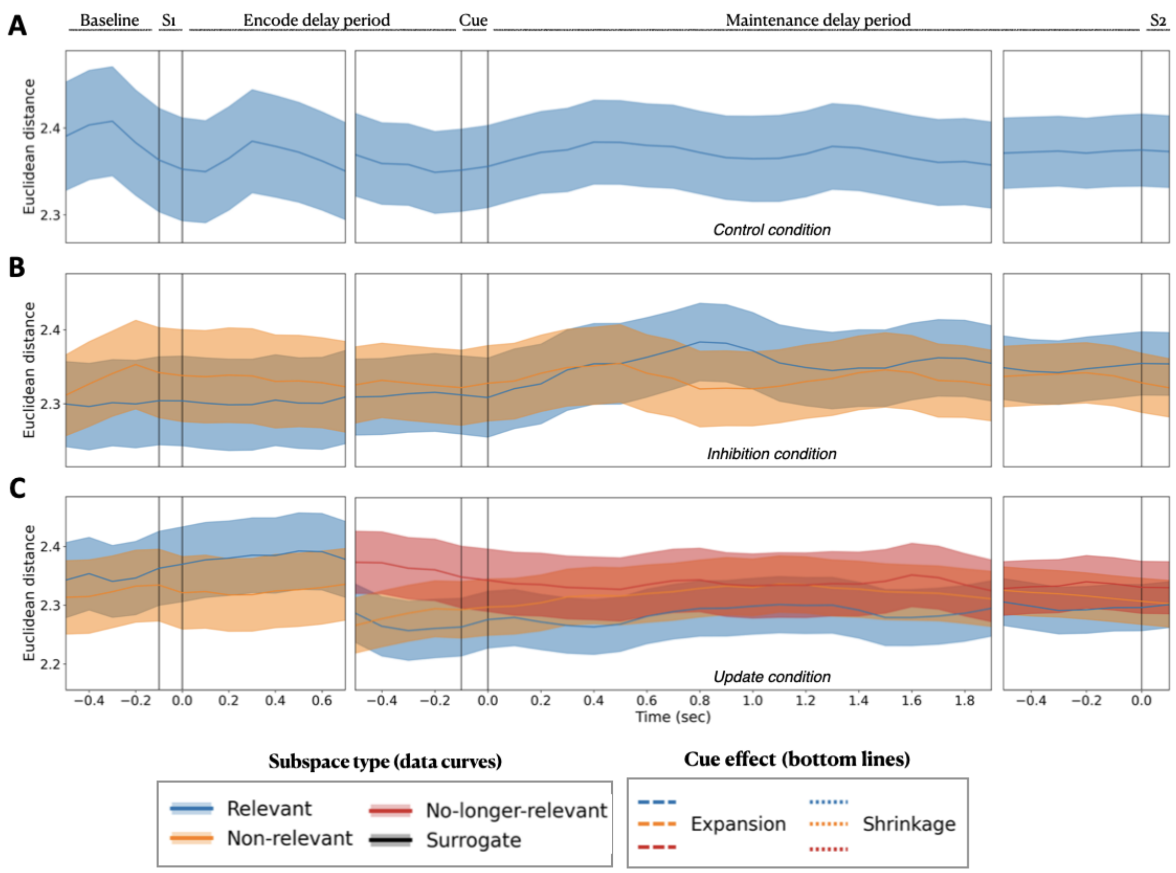


**Figure S23.** Supplementary analysis of neural geometry with hyperalignment applied on the 3 PCs of Z matrices of PC scores, instead of the 10 PCs used in the main analysis. Separability of memory representations within feature-specific neural subspaces for correct trials, shown for A) Control condition, B) Inhibition condition, and B) Update condition. Significant timepoints of cue effect (bottom lines) are FDR-corrected for multiple comparisons. Panel layout, time axis, and significance markers follow the conventions described in Figure S2.

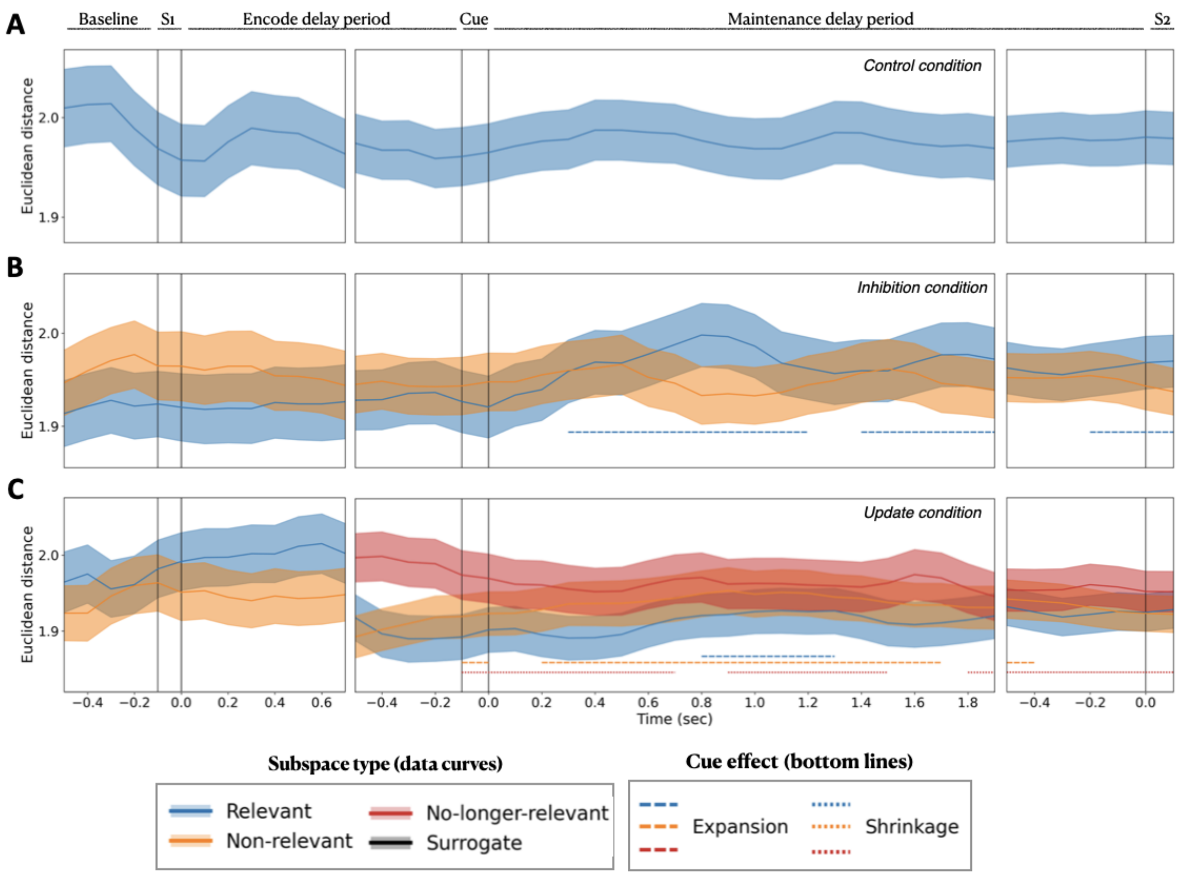


**Figure S24.** Supplementary analysis of neural geometry using iterative hyperalignment across subjects. Separability of memory representations within feature-specific neural subspaces for correct trials, shown for A) Control condition, B) Inhibition condition, and B) Update condition. Significant timepoints of cue effect (bottom lines) are FDR-corrected for multiple comparisons. Panel layout, time axis, and significance markers follow the conventions described in Figure S2.

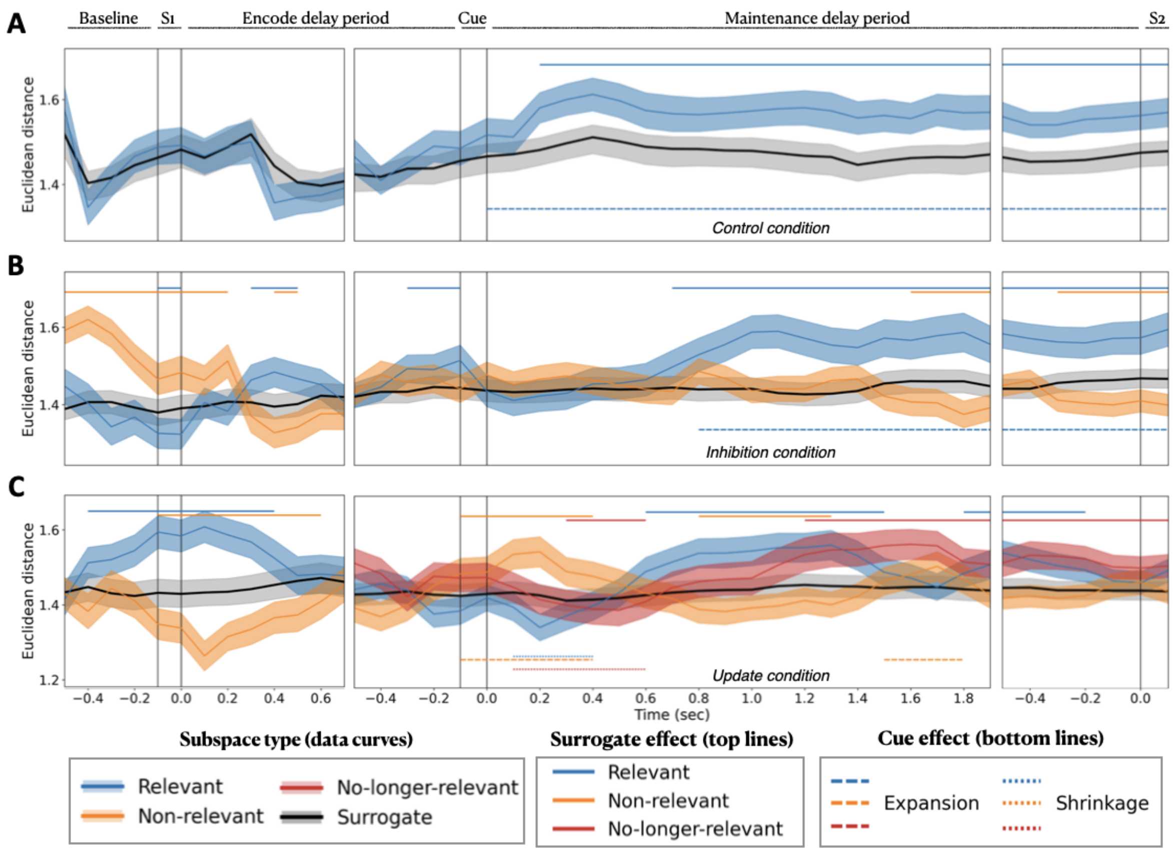


**Figure S25.** Supplementary analysis of neural geometry where subspaces where defined with leading 2 PCs. Separability of memory representations within feature-specific neural subspaces for correct trials, shown for A) Control condition, B) Inhibition condition, and B) Update condition. Significant timepoints of surrogate effect (top lines) and cue effect (bottom lines) FDR-corrected for multiple comparisons. Panel layout, time axis, and significance markers follow the conventions described in Figure S2.

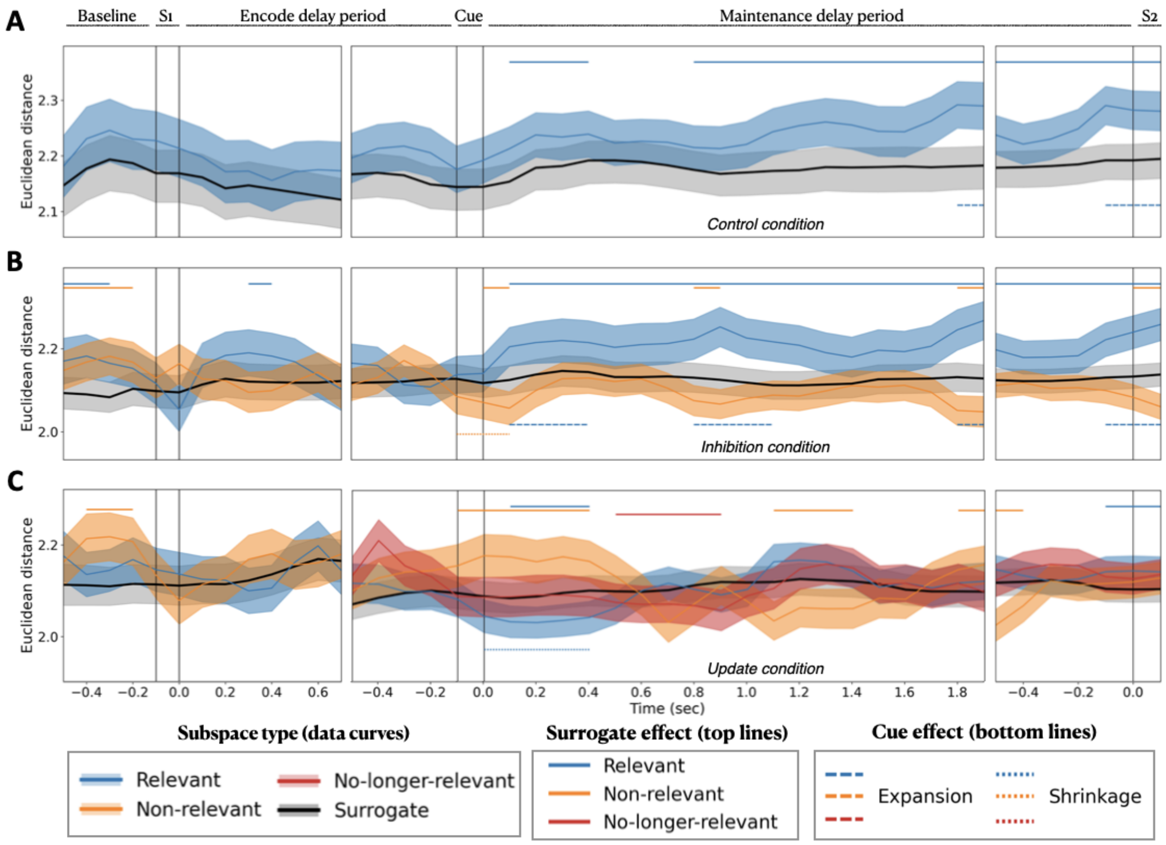


**Figure S26.** Supplementary analysis of neural geometry where subspaces where defined with leading 4 PCs. Separability of memory representations within feature-specific neural subspaces for correct trials, shown for A) Control condition, B) Inhibition condition, and B) Update condition. Significant timepoints of surrogate effect (top lines) and cue effect (bottom lines) are FDR-corrected for multiple comparisons. Panel layout, time axis, and significance markers follow the conventions described in Figure S2.

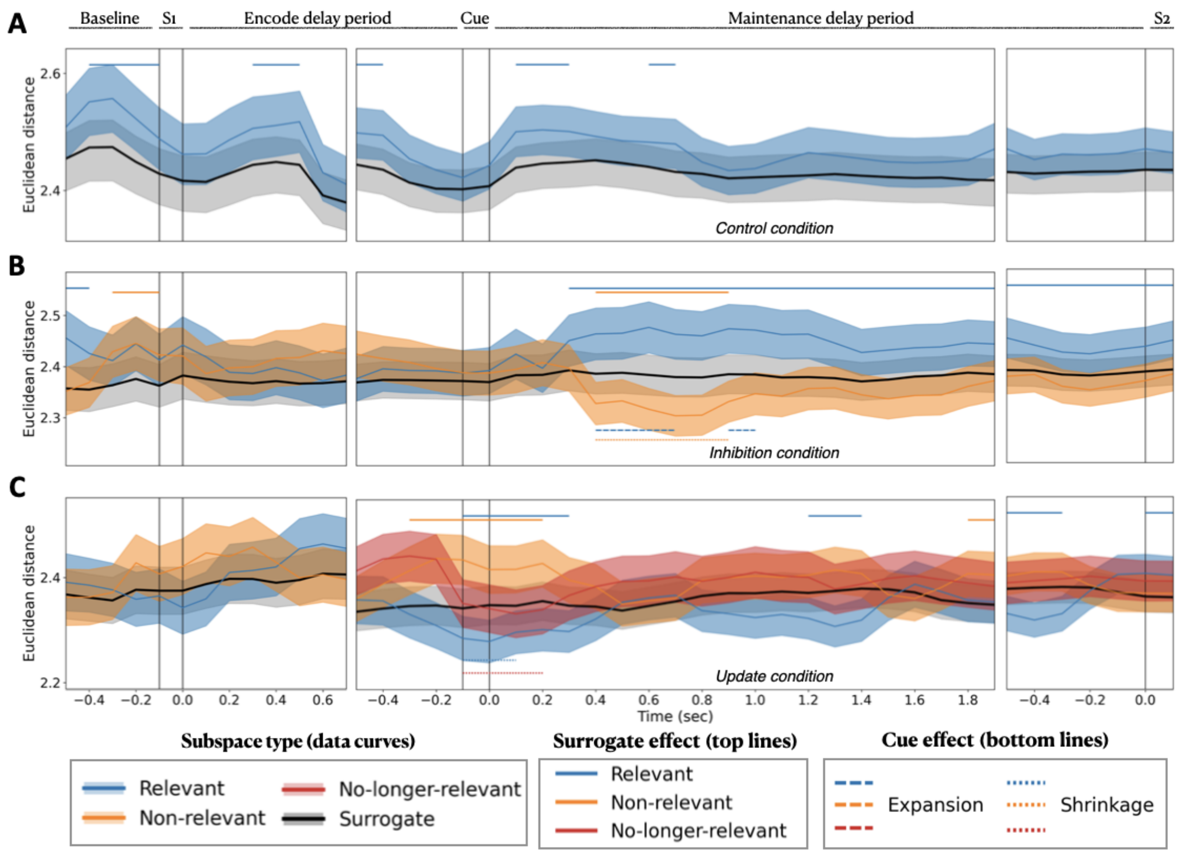


**Figure S27.** Supplementary analysis of neural geometry where subspaces where defined with leading 5 PCs. Separability of memory representations within feature-specific neural subspaces for correct trials, shown for A) Control condition, B) Inhibition condition, and B) Update condition. Significant timepoints of surrogate effect (top lines) and cue effect (bottom lines) are FDR-corrected for multiple comparisons. Panel layout, time axis, and significance markers follow the conventions described in Figure S2.

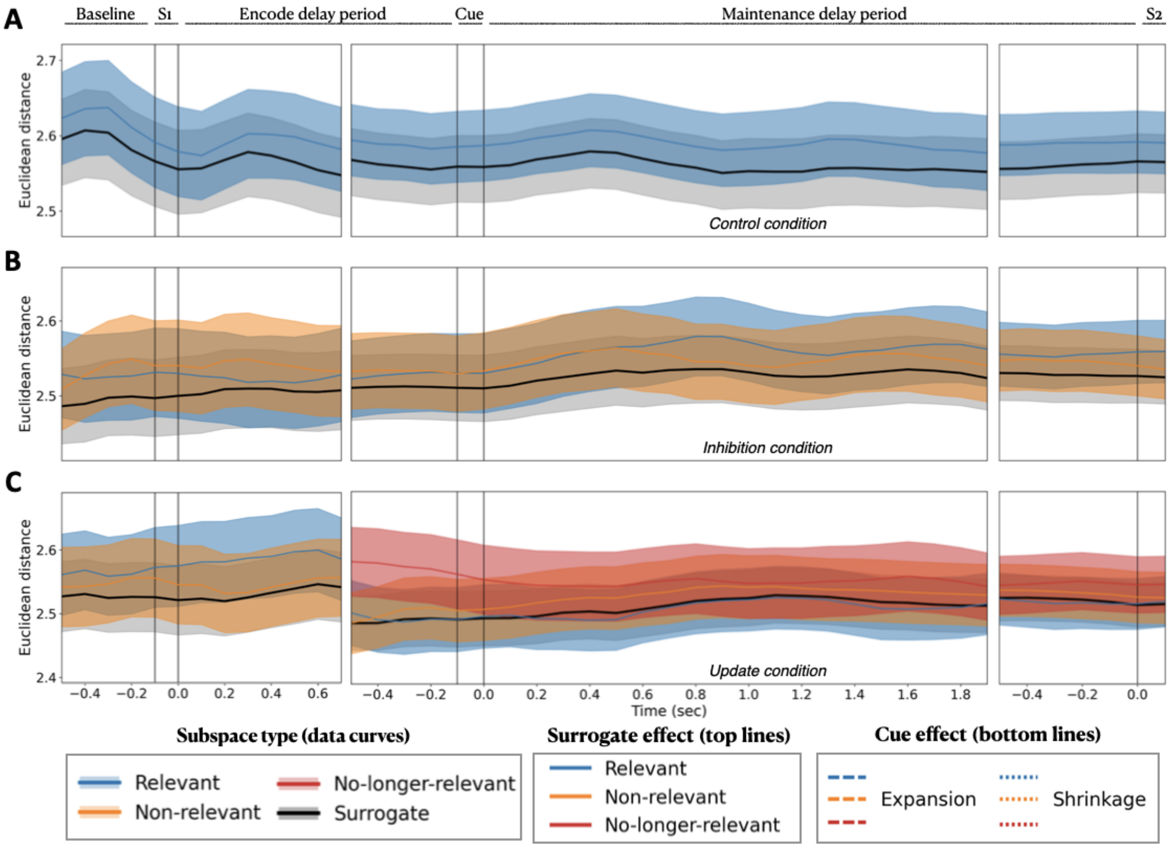


**Figure S28.** Supplementary analysis of neural geometry where subspaces where defined with leading 6 PCs. Results were identical when using 8 or 10 PCs. Separability of memory representations within feature-specific neural subspaces for correct trials, shown for A) Control condition, B) Inhibition condition, and B) Update condition. Significant timepoints of surrogate effect (top lines) and cue effect (bottom lines) are FDR-corrected for multiple comparisons. Panel layout, time axis, and significance markers follow the conventions described in Figure S2.

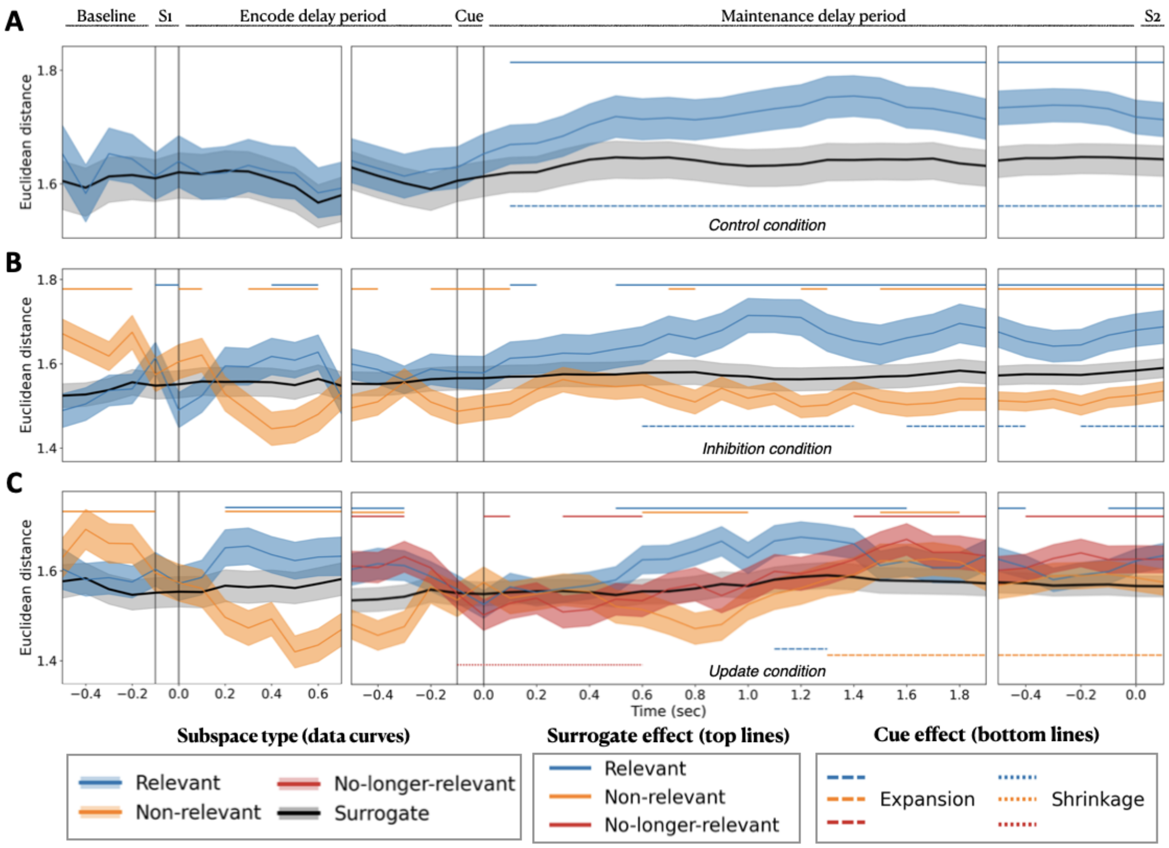


**Figure S29.** Supplementary analysis of neural subspaces estimated using a cortical parcellation of 300 areas instead of 400. Separability of memory representations within feature-specific neural subspaces for correct trials, shown for A) Control condition, B) Inhibition condition, and B) Update condition. Significant timepoints of surrogate effect (top lines) and cue effect (bottom lines) are FDR-corrected for multiple comparisons. Panel layout, time axis, and significance markers follow the conventions described in Figure S2.

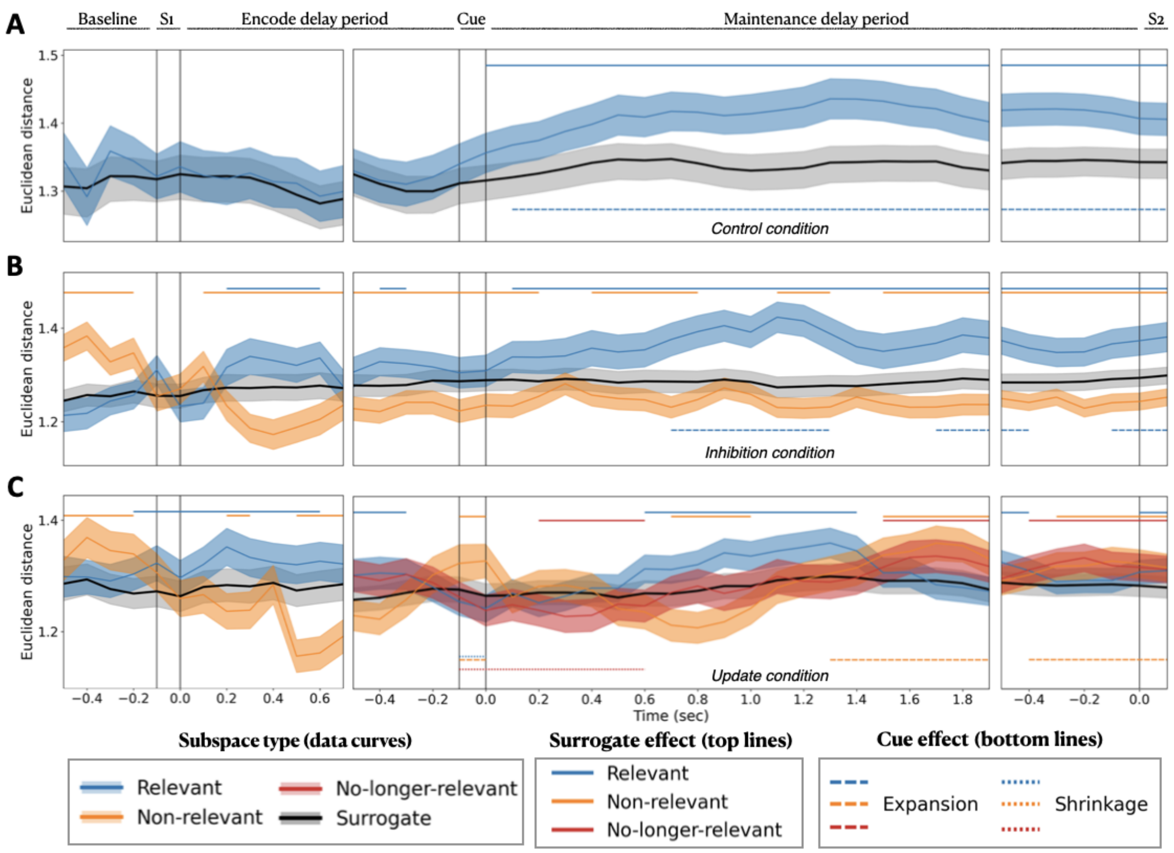


**Figure S30.** Supplementary analysis of neural subspaces estimated using a cortical parcellation of 200 areas instead of 400. Separability of memory representations within feature-specific neural subspaces for correct trials, shown for A) Control condition, B) Inhibition condition, and B) Update condition. Significant timepoints of surrogate effect (top lines) and cue effect (bottom lines) are FDR-corrected for multiple comparisons. Panel layout, time axis, and significance markers follow the conventions described in Figure S2.

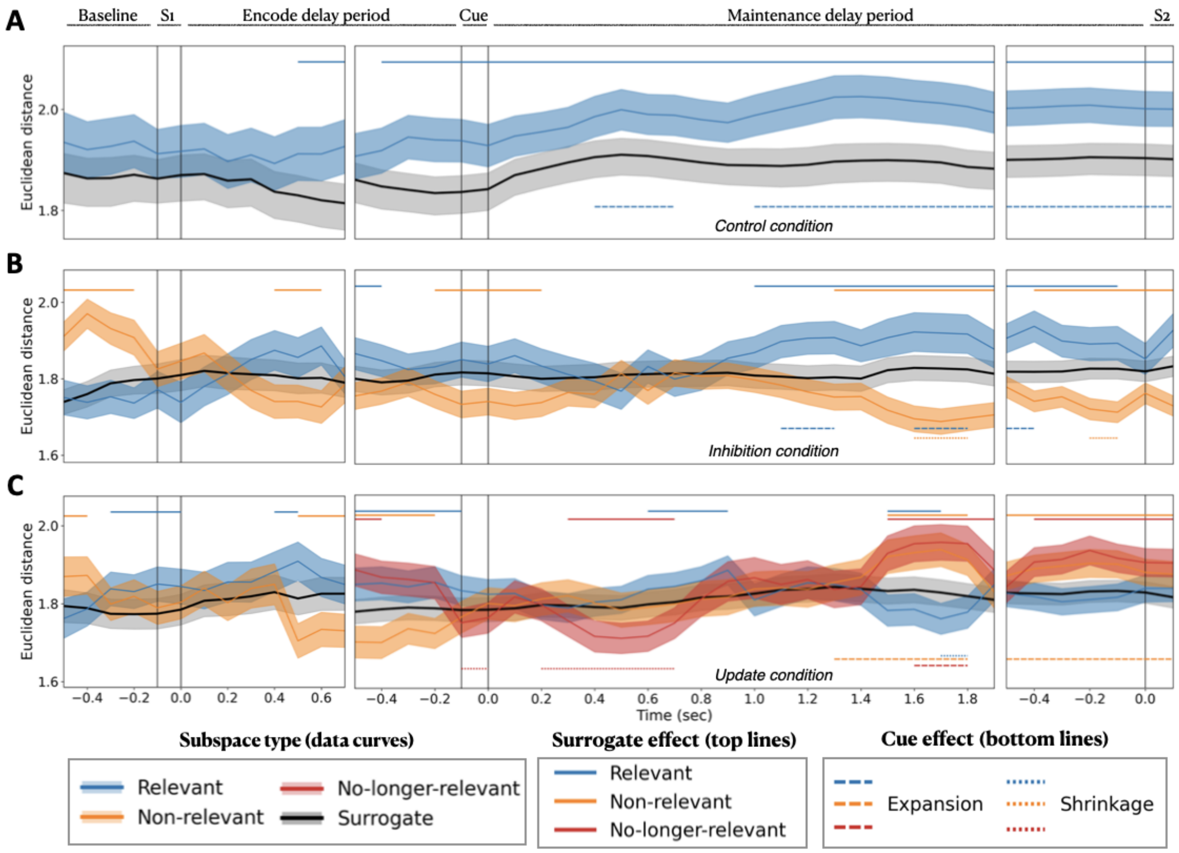


**Figure S31.** Supplementary analysis of neural geometry using the same preprocessing pipeline as in the main analyses, but without applying fidelity weighting to the source time series. Separability of memory representations within feature-specific neural subspaces for correct trials, shown for A) Control condition, B) Inhibition condition, and B) Update condition. Significant timepoints of surrogate effect (top lines) and cue effect (bottom lines) are FDR-corrected for multiple comparisons. Panel layout, time axis, and significance markers follow the conventions described in Figure S2.

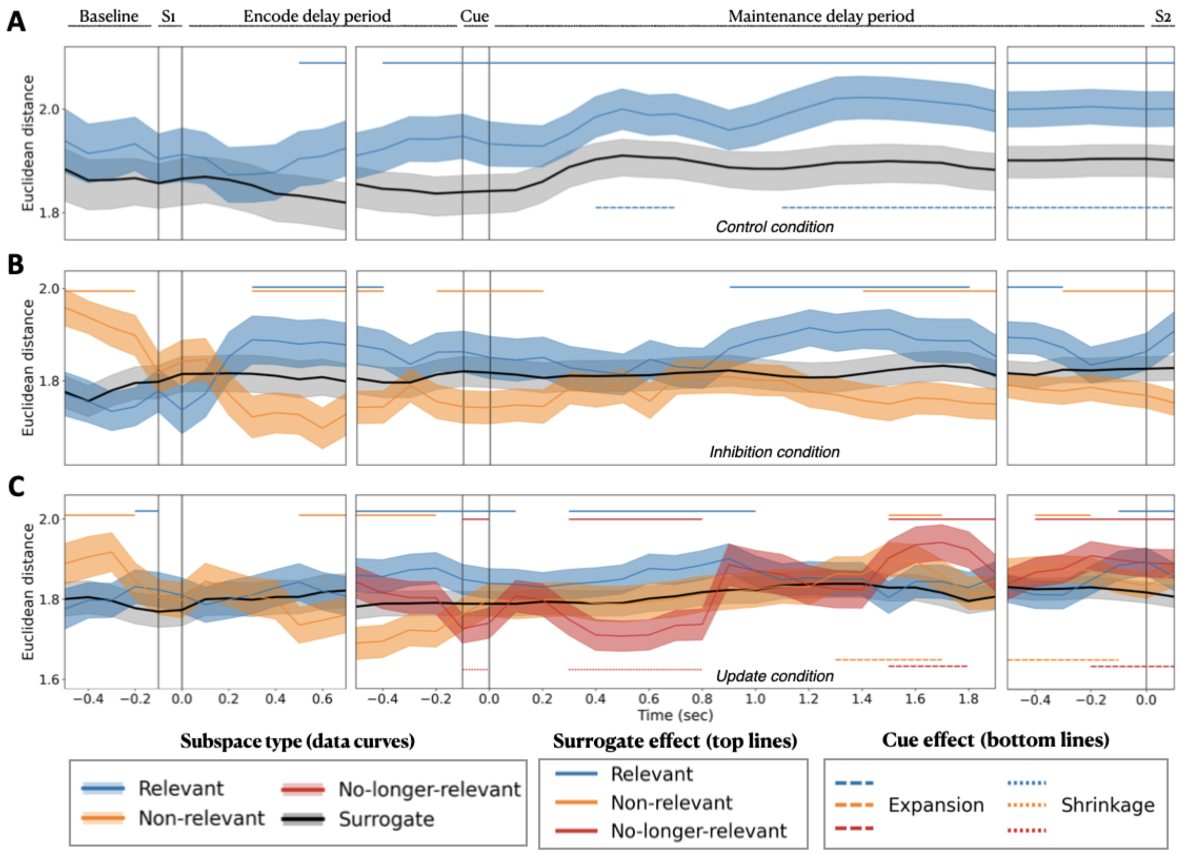


**Figure S32.** Supplementary analysis of neural geometry using a standard pipeline with MNE method. Separability of memory representations within feature-specific neural subspaces for correct trials, shown for A) Control condition, B) Inhibition condition, and B) Update condition. Significant timepoints of surrogate effect (top lines) and cue effect (bottom lines) are FDR-corrected for multiple comparisons. Panel layout, time axis, and significance markers follow the conventions described in Figure S2.

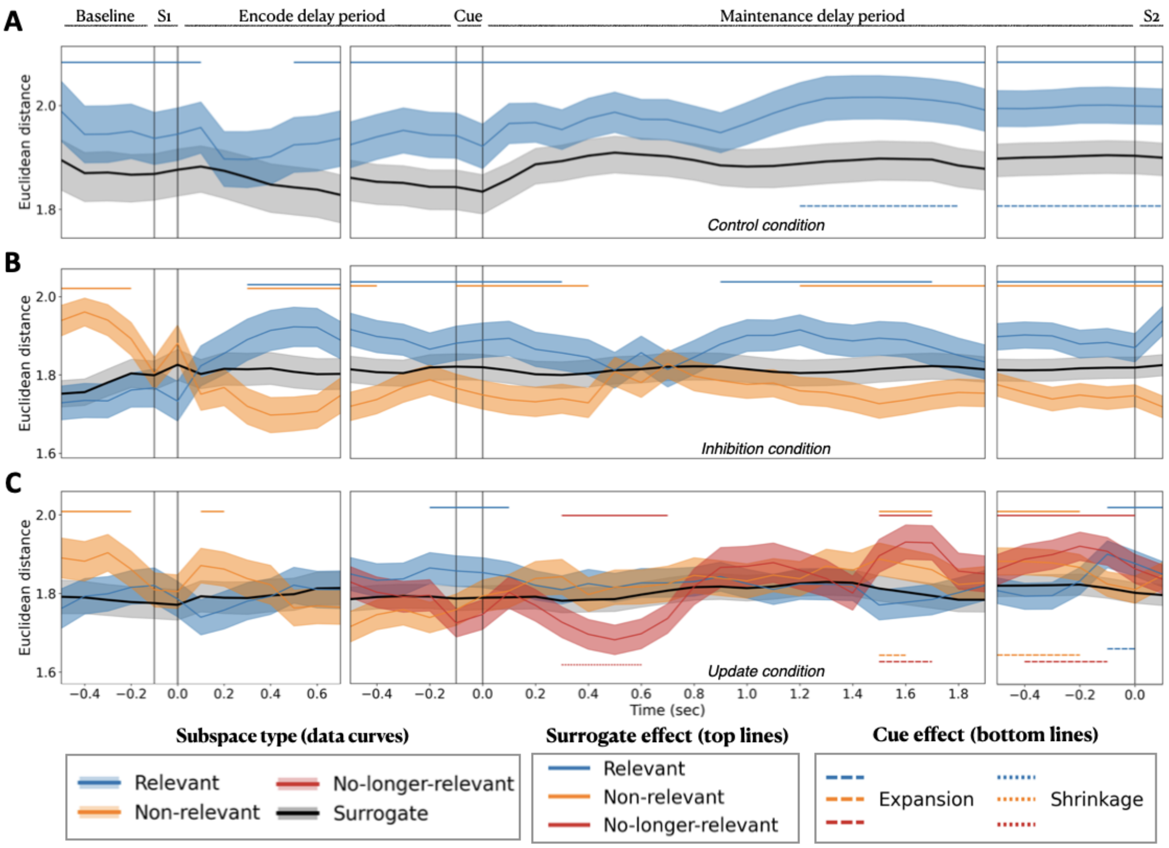


**Figure S33.** Supplementary analysis of neural geometry using a standard pipeline with dSPM method. Separability of memory representations within feature-specific neural subspaces for correct trials, shown for A) Control condition, B) Inhibition condition, and B) Update condition. Significant timepoints of surrogate effect (top lines) and cue effect (bottom lines) are FDR-corrected for multiple comparisons. Panel layout, time axis, and significance markers follow the conventions described in Figure S2.

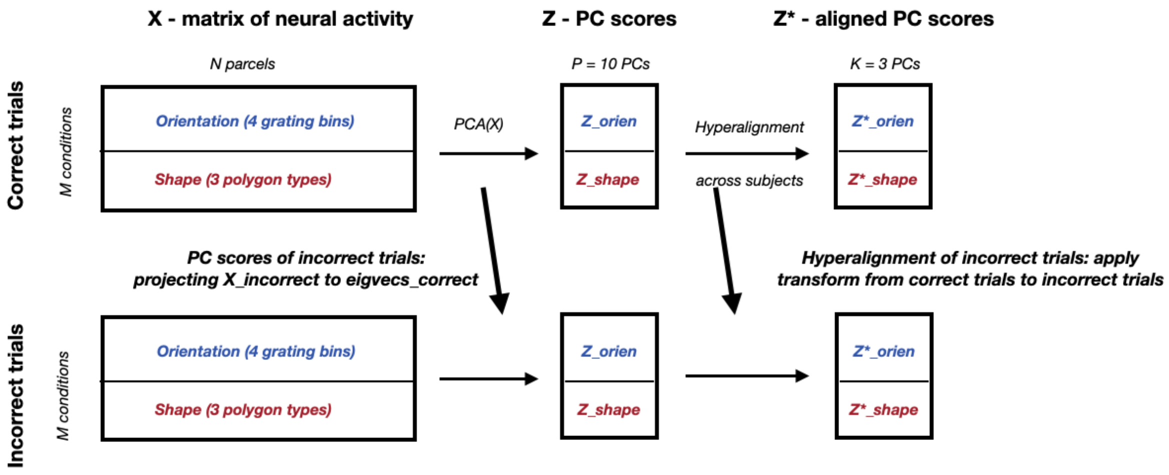


**Figure S34.** Geometric analysis of neural subspaces. Estimation of subspaces for incorrect trials: X_incorrect_trials is projected onto the eigenvectors derived from correct trials, and the hyperalignment computed for correct trials is applied to incorrect trials.

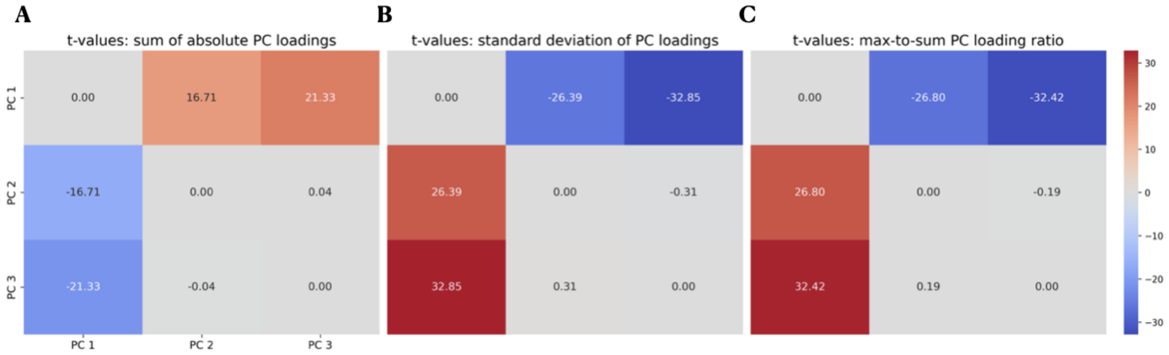


**Figure S35.** Identification of the shared broadband PC from NB separability metrics. Pairwise t-values across subjects are shown for three metrics quantifying the distribution of PC loadings across frequency bands: (A) sum of absolute loadings, which was used to select the broadband-like (aperiodic) PC; (B) standard deviation of loadings; and (C) maximum-to-sum loading ratio. The first PC consistently exhibited the most distributed pattern across bands compared with the other components. Results are shown for NB components derived from the real part of the complex Morlet wavelet time series using subject-level frequency clustering. No qualitative differences were observed when using group-level clusters or amplitude envelopes.
